# The basal cell state maintains pancreatic cancers by controlling an immunosuppressive circuit

**DOI:** 10.64898/2026.07.13.738221

**Authors:** Anupriya Singhal, Kate Ryan, Samuel A. Rose, Hannah C. Styers, Jung Yun Kim, Nikhita Pasnuri, Ashlyn Moore, Jose Meza-Llamosas, Emma Chen, Josephine Adams, Ananya Nandula, Roshan Sharma, Zhuxuan Li, Tal Nawy, Yan Yan, Nuray Tezcan, Olca Basturk, Mara H. Sherman, Dana Pe’er, Tuomas Tammela

## Abstract

Intra-tumoral heterogeneity is a cardinal feature of solid tumors, yet how distinct cancer cell states functionally contribute to malignant and stromal diversity *in situ* remains poorly understood. Using mouse models to lineage-trace or genetically ablate the two predominant cancer cell states in autochthonous pancreatic ductal adenocarcinoma (PDAC), we discover that basal cancer cells are highly plastic, whereas classical cancer cells exhibit limited plasticity. Strikingly, ablation of the basal, but not the classical, state induced rapid and durable tumor collapse, driven by loss of immunosuppressive cancer-associated fibroblasts, macrophage repolarization, and reprogramming of the tumor cytokine milieu, culminating in tumor destruction by cytotoxic lymphocytes. Knockout of a single cytokine, GM-CSF, specifically in basal cells recapitulated macrophage repolarization and lymphocyte recruitment observed upon basal state ablation and shrank tumors. These results reveal the basal cell state controls an immunosuppressive cell circuit critical for PDAC maintenance, motivating therapeutic targeting of the basal cells.

---

Even within genetically homogenous tumors, malignant cells adopt recurrent phenotypic states that are conserved across cancer types (*1, 2*). These states are often viewed through a cell-intrinsic lens—as programs of epithelial-to-mesenchymal transition (EMT), lineage differentiation, or stress adaptation. However, increasing evidence suggests that malignant states may also encode distinct capacities to interact with and organize the tumor microenvironment (TME). Spatial profiling studies across malignancies and disease stages show that defined cancer cell states associate with distinct stromal and immune niches (*3–5*). A central question is whether specific malignant states actively instruct TME architecture, or instead passively occupy TMEs shaped by other host-derived factors.

Intratumoral heterogeneity is a defining feature of pancreatic ductal adenocarcinoma (PDAC), a common and highly lethal solid tumor with a 5-year survival rate of only 13% (*6*). Despite sharing core driver alterations—including *KRAS*, *TP53*, *SMAD4*, and *CDKN2A*—PDAC cells adopt stereotyped transcriptional programs. Two dominant states have emerged: a classical state associated with glandular differentiation and epithelial identity, and a basal state characterized by basal epithelial and quasi-mesenchymal features(*7, 8*). Although individual tumors may be enriched for one state, classical and basal programs invariably coexist within the same tumor(*9–13*). How specific cancer cell states persist, transition, and interact with the TME during cancer progression and under therapeutic pressure represents a key knowledge gap in PDAC biology and clinical management. To address this critical question, new experimental strategies and model systems enabling elucidation of the dynamic nature of plastic transitions in tumors *in situ* are needed.

Basal state-dominant PDAC tumors are consistently linked to the poorest outcomes, yet the basal state-specific properties that drive aggressive behavior remain poorly understood. A central challenge in PDAC is its dense, fibrotic, and profoundly immunosuppressive TME, marked by cancer-associated fibroblasts (CAFs) and myeloid populations that suppress antitumor immunity while cytotoxic lymphocytes are excluded or exhausted. Human PDAC atlases show that basal-dominant tumors harbor reduced immune diversity, fewer CD4^+^ and CD8^+^ T cells, and abundant CAFs with elevated MAPK signaling, a subset associated with T cell exhaustion (*9, 14*). Human spatial profiling studies have begun to define recurrent local niches in PDAC, including niches in which basal cancer cells are spatially associated with myofibroblastic CAFs (*5, 15, 16*). However, these data remain largely correlative and derive from single-timepoint profiling. Without direct perturbation, it remains unknown whether basal malignant cells directly instruct this immunosuppressive ecosystem.

A major obstacle has been the lack of experimental systems for direct functional interrogation of cancer cell states *in situ*. First, it is not known how selective ablation of a defined cancer cell state—essentially subtraction of the entire functional program it encodes—remodels the tumor ecosystem. Thus, the dependencies of specific stromal and immune subsets on defined cancer cell states remain unknown. Second, because cytokine signaling operates within multidirectional cancer cell–TME circuits, conventional genetic and pharmacologic perturbations cannot determine whether the cellular source of a cytokine controls specific stromal phenotypes. Third, transitions between the basal and classical cell states have not been elucidated in time-dynamic experiments *in situ* in PDAC tumors, particularly when one state is selectively perturbed.

To address these limitations, we developed genetic systems for functional interrogation of cancer cell states in autochthonous PDAC tumors that develop in immunocompetent hosts. Our platform builds on genetically engineered mouse models (GEMMs), in which PDAC develops through oncogenic *KrasG12D* activation and *Trp53* loss in the pancreatic epithelium induced by Cre or Flp recombinase activity to the pancreatic epithelium (*KPC* or *KPF* models, respectively) (*17, 18*). This gold-standard model recapitulates key features of human PDAC, including classical and basal states and a desmoplastic, immunosuppressive TME. By engineering state-specific perturbation, our tools enable longitudinal fate mapping, temporally controlled state elimination, and cell state-specific gene perturbation. We combine these approaches with longitudinal spatial profiling to define how basal and classical states contribute to tumor maintenance, plasticity, and architecture of the tumor ecosystem in PDAC.

## RESULTS

### Reporter systems enable functional interrogation of PDAC basal and classical states *in situ*

The basal and classical transcriptional programs define major epithelial states in PDAC, yet their roles in shaping tumor behavior and the TME remain unresolved due to a lack of tools for selectively manipulating them *in vivo*. To directly interrogate the functional contributions of cancer cell states *in situ*, we developed GEMMs that enable cell state-specific labeling, lineage tracing, and ablation in autochthonous PDAC tumors (**Fig. 1A**). Our *Hipp11-BG* reporter system includes a Flp-inducible blue fluorescent protein (tagBFP) and Cre-inducible GFP that can be used to mark PDAC lineages *in vivo* via state-specific Cre recombinase activity (**Fig. 1A; Fig. S1A-C**).

**Figure 1.**
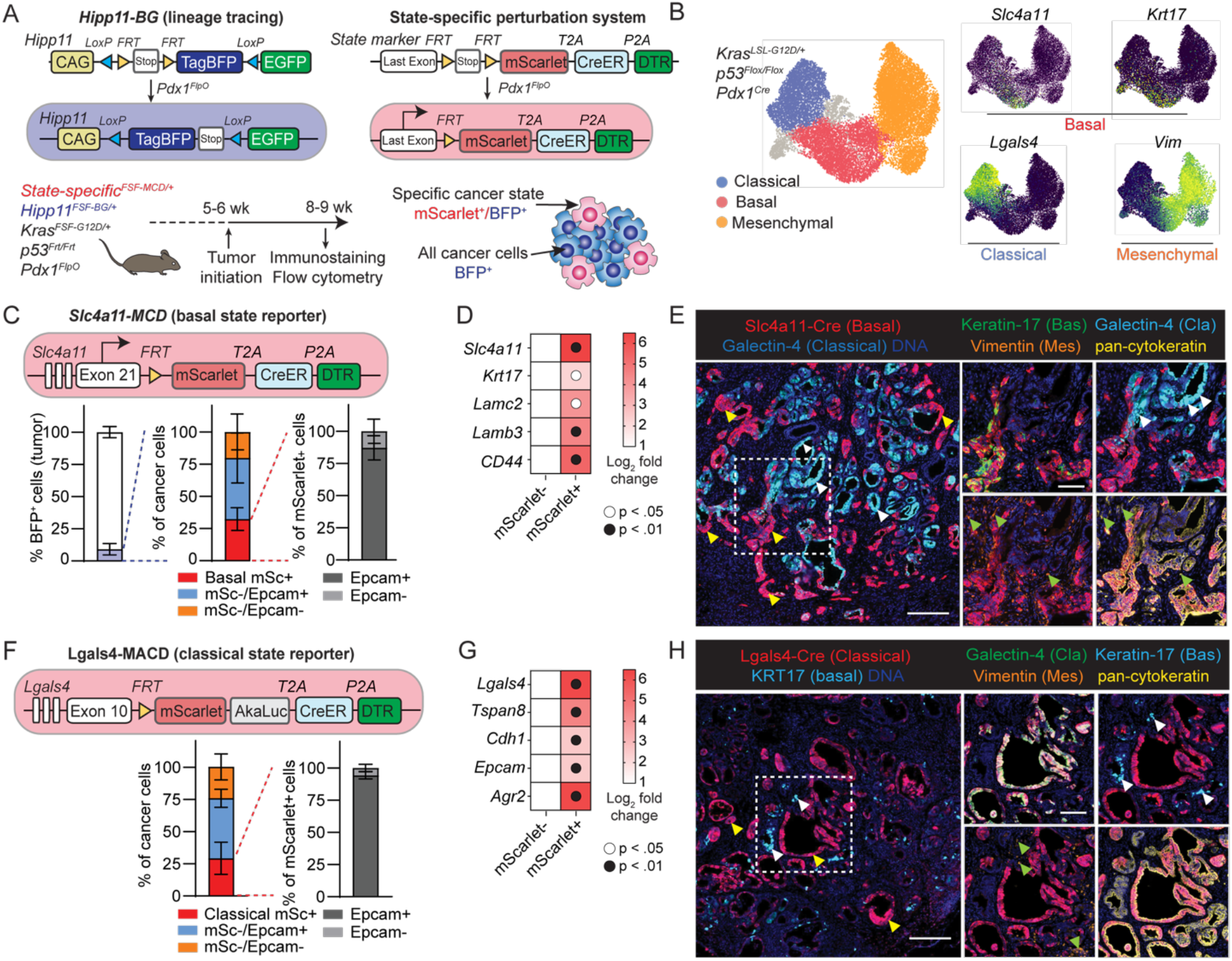
Reporter systems for *in situ* functional interrogation of basal and classical states in autochthonous PDAC tumors. **(A)** Basal and classical reporter systems each consist of two components. (i) The *Hipp11^FSF-BG/+^*allele, upon pancreas-specific recombination mediated by *Pdx1-*driven *FlpO* (upper left), provides baseline tagBFP expression and Cre-dependent switch to GFP for lineage tracing. (ii) A state-specific perturbation reporter (upper right) is activated in the pancreatic epithelium following *Pdx1^FlpO^*-mediated FSF cassette excision, such that cells of the corresponding state express mScarlet for visualization, CreER^T2^ for lineage tracing, and DTR for ablation (MCD cassette). *KPF* (*Kras^FSF-^_G12D/+_;Trp53^frt/frt^;Hipp11^FSF-BG/+^*) mice harboring *state-specific^FSF-MCD/+^* alleles develop autochthonous PDAC and reach tumor burden endpoint at 8–9 weeks (lower left). In these mice, BFP marks all PDAC cells and mScarlet marks the cell state of interest (lower right). **(B)** Uniform manifold approximation and projection (UMAP) embedding of autochthonous PDAC cell transcriptomes from Pitter *et al*. (*12*), colored by cancer cell state (left) or key cell state markers (right).**(C)** Flow cytometric quantification of tumor cell composition (bottom) in *KPF-basal* mice harboring the *Slc4a11^FSF-MCD/+^* reporter (top). *n* = 5 mice; mSc, mScarlet. **(D)** Basal marker gene expression in cells isolated by flow cytometry from *KPF-basal* tumors based on mScarlet fluorescence. *n* = 3 independent tumors per cell population; *p* values, Welch’s t-test. **(E)** Immunofluorescence of *KPF-basal* tumors. Cre (basal cell) and Galectin-4 (classical) localization is largely mutually exclusive (left). Scale bar, 100 µm. Magnifications of boxed region reveal co-localization of Cre with basal marker Keratin-17. Scale bar, 50 µm. Yellow arrowheads, basal cells; white arrowheads, classical cells; green arrowheads, Vimentin^+^ mesenchymal cancer cells. **(F)** Flow cytometric quantification of tumor cell composition (bottom) in *KPF-classical* mice harboring the *Lgals4^MACD/+^*reporter (top), which contains the MCD cassette with bright AkaLuc bioluminescent report (MACD). *n* = 5 mice. mSc, mScarlet. **(G)** Classical marker gene expression in cells isolated by flow cytometry from *KPF-classical* tumors based on mScarlet fluorescence. *n* = 3 biological replicates per group; *p* values, Welch’s t-test. **(H)** Immunofluorescence of *KPF-classical* tumors. Cre (classical cell) and Keratin-17 (basal) localization is largely mutually exclusive (left). Scale bar, 100 µm. Magnifications of boxed region reveal co-localization of Cre with Galectin-4. Arrowheads highlight basal (yellow), classical (right), or mesenchymal (green) cancer cells. Scale bar, 50 µm.

To generate state-specific reporters, we used our previous single-cell RNA sequencing (scRNA-seq)-based characterization of the *KPC* model, which identified classical, basal, and mesenchymal states that are enriched for glandular epithelial differentiation, human-conserved laminins and keratins, and fibroblast-like transcriptional features, respectively (*12*). For state-specific knock-in loci, we chose *Slc4a11—*a basal state marker in PDAC GEMMs that correlates strongly with *Krt17*, a well-established basal marker in human disease and mouse models (*7, 8*)—and *Lgals4* (encoding galectin-4)—a core classical marker in mice and the most highly correlated marker of classical identity in a recent human scRNA-seq study (**Fig. 1B; Fig. S1D–G**) (*7–9*). For *Slc4a11*, we resorted to our recently reported *MCD* reporter allele (*19*), comprised of mScarlet for visualizing and isolating basal cells within tumors; tamoxifen-inducible Cre (CreER^T2^) for lineage tracing; and the diphtheria toxin receptor (DTR) suicide gene for cytoablation (**Fig. 1C-E**). To mark classical cells, we knocked in a *MACD* allele into the *Lgals4* locus containing the same components in addition to a super-bright AkaLuc bioluminescence reporter, enabling longitudinal tracking of classical cells *in vivo* (**Fig. 1F-H; Fig. S1H,I**) (*20*). We introduced each allele into *KPF* mice together with *Hipp11-BG*, generating *KPF; Hipp11-BG; Slc4a11-MCD* (hereafter, the *KPF-basal* model), and *KPF; Hipp11-BG; Lgals4-MACD* (hereafter *KPF-classical*) mice. In both models, the target cell state is marked by mScarlet, all PDAC cells are initially BFP⁺, and a single pulse of tamoxifen leads to a heritable genetic switch of BFP to GFP for lineage tracing (**Fig. 1A**). Mice harboring either reporter developed aggressive autochthonous PDAC with reproducible mixtures of basal, classical, and mesenchymal populations (**Fig. 1C,F; Fig. S1J,K**) and reached humane endpoint at approximately 8–9 weeks (**Fig. 1C**), indicating that the reporter constructs do not impact PDAC progression.

The reporter systems label their intended cell types based on marker expression, marker localization and morphology. In *KPF-basal*, mScarlet⁺ basal cells comprised 32.4 ± 8.6% of BFP⁺ cancer cells and 3.2 ± 2.1% of all cells within the tumor, and nearly 90% of mScarlet^+^ cells expressed EpCAM, consistent with epithelial identity (**Fig. 1C, Fig. S1J**). *KPF-basal* and *KPF-classical* tumors contained similar proportions of basal and classical cancer cell states, indicating comparable underlying state composition across the models (**Fig. 1C**, **Fig. 1F, Fig. S1J,K**). Reporter-positive cells isolated from *KPF-basal* and *KPF-classical* were enriched for their respective canonical markers (**Fig. 1D,G**). mScarlet^+^ cells in *KPF-basal* expressed KRT17, lacked galectin-4 and vimentin, and displayed a flattened basal epithelial morphology, with some localizing basally to the classical cells and some showing invasive features (**Fig. 1E**). In contrast, mScarlet^+^ *KPF-classical* cells expressed Cre and galectin-4, lacked KRT17, and formed columnar epithelia lining gland-like lumina (**Fig. 1H**). Vimentin⁺ mesenchymal cells were spatially distinct and interdigitated with stromal compartments (**Fig. 1E**). Basal cells were relatively quiescent compared to classical cells, consistent with our previous work (*19*) (**Fig. S1L**). These data establish *KPF-basal* and *KPF-classical* as complementary systems that enable functional interrogation of the two dominant cancer states in PDAC, which differ in transcriptional programs, spatial organization, and proliferative capacity.

### The basal state harbors high plasticity *in situ* in PDAC tumors

To determine the differentiation potential of basal PDAC cells *in vivo*, we performed lineage tracing in established *KPF-basal* PDAC tumors (**Fig. 2A**). A single dose of tamoxifen selectively labeled basal cells and their progeny with GFP. While GFP^+^ cells at day 3 faithfully labeled basal cells as expected, by day 14, GFP^+^ basal-derived cells had frequently adopted non-basal identities, including classical identity with glandular epithelial morphology and galectin-4 expression, as well as mesenchymal features characterized by spindle-like morphology (**Fig. 2B**). Consistent with this, the fraction of lineage-traced cells that exited the basal state (mScarlet^-^ among total GFP^+^ cells) increased sharply from negligible on day 3 to 41.5% ± 10.3% on day 14 (**Fig. 2C,D**).

**Figure 2.**
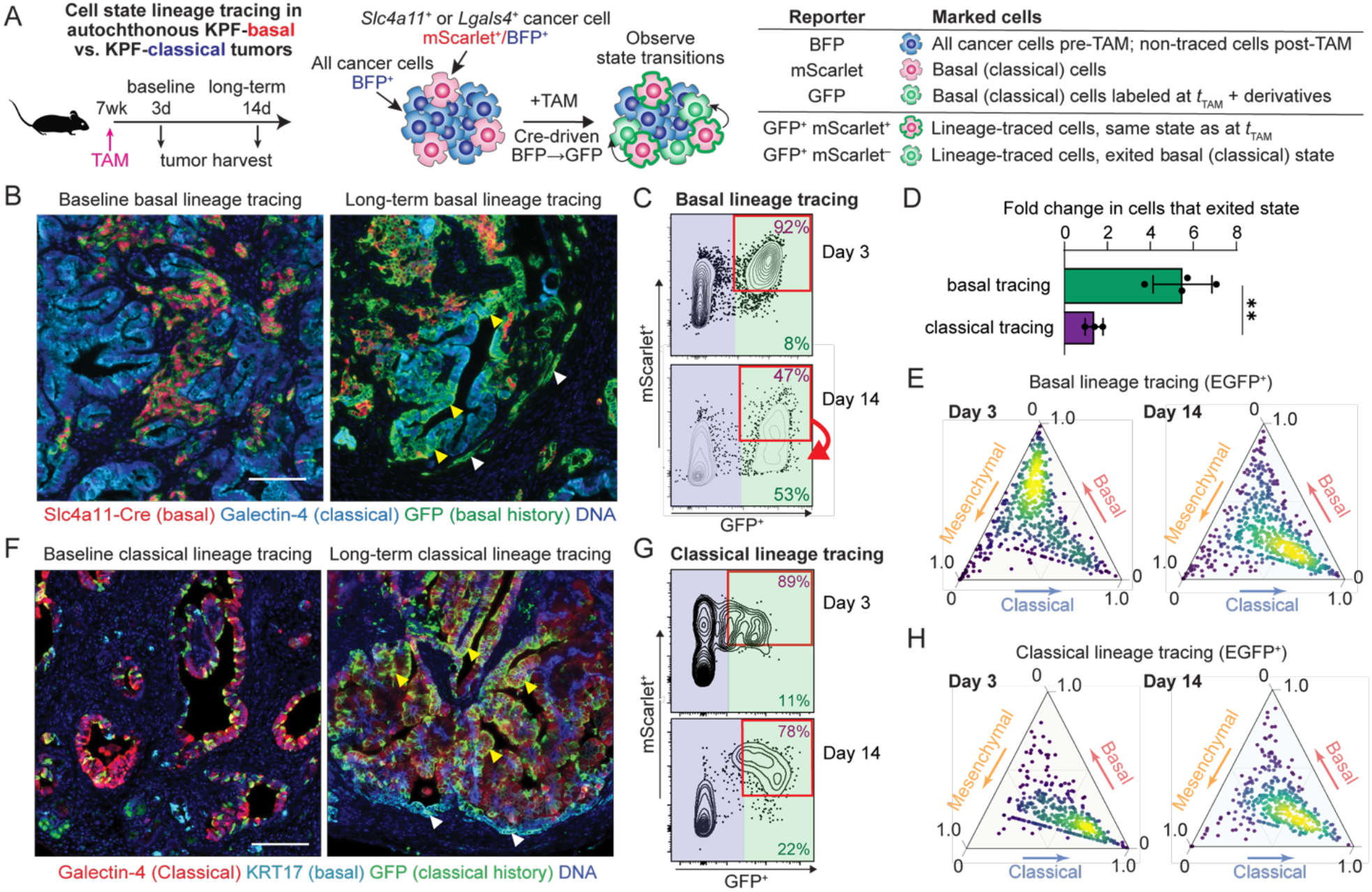
The basal PDAC cell state, but not the classical state, undergoes frequent and rapid transdifferentiation *in situ*. **(A)** Experimental design for tracing basal cells and their descendants to test for state transitions. Tamoxifen (TAM) administration triggers a permanent switch from BFP to GFP expression via Cre activity in *KPF-basal* mice harboring established autochthonous PDAC tumors. Tumors are harvested 3 days (baseline) and 14 days (long-term) after initiating lineage tracing (time *t*_TAM_) to test whether descendants of the traced state (marked by GFP) retain or lose their original identity (marked by mScarlet). (**B,C**) *KPF-basal* tumors induced for lineage tracing and subjected to immunofluorescence (B, scale bar 100 μm) and cell-state quantification by live tumor cell flow cytometry (C). **(D)** Fraction of lineage-traced cells that exited the initially labeled state, calculated as fold change in mScarlet⁻GFP⁺ cells/total GFP⁺ cells at day 14 relative to day 3. Basal cells undergo cell-state exit at much higher rates than classical cells. **, *p* = 0.0019, Welch’s t-test. **(E)** Probabilistic prediction of cell states from scRNA-seq show a shift from pure basal states at baseline to basal exit and acquired classical and mesenchymal identity in long-term basal-lineage-traced cells (Methods). (**F–H**) *KPF-classical* tumors induced for lineage tracing and subjected to immunofluorescence (F, scale bar 100 μm), live cell flow cytometry (G), and scRNA-seq (H) show stable maintenance of classical states in baseline and long-term lineage-traced cells.

To interrogate cell states derived from the basal cells at high resolution, we performed scRNA-seq of lineage-traced cells in autochthonous *KPF-basal* tumors. We reasoned that state transitions may not occur as abrupt switches between discrete identities, but instead involve movement through a continuous malignant state phenotypic space. We therefore scored each lineage-traced cell for basal, classical, and mesenchymal transcriptional programs derived from an independent *KPF* PDAC reference dataset (**Table S1**), normalized these scores to make them comparable across programs, and transformed them into fractional cell state weights (**Methods**). Plotting these weights confirmed the selective capture of the basal state at day 3, and revealed a broad distribution spanning basal, classical, mesenchymal, and mixed/intermediate identities by day 14 (**Fig. 2E**). Taken together, these findings indicate that basal cells harbor high plasticity and lineage potential in PDAC tumors *in vivo*.

### The classical state harbors low plasticity in PDAC

To test whether the high capacity for lineage-switching (plasticity) is unique to the basal cells, we next interrogated classical state differentiation potential *in situ* in established autochthonous *KPF-classical* PDAC tumors (**Fig. 2F**). In contrast to the basal state, the classical state showed restricted state transitions: over a 2-week interval, only 15.7% ± 5.7% of the classical cells adopted alternative identities, as opposed with the significantly higher plasticity of basal cells (41.5% ± 10.3%) (**Fig. 2D,G)**. Following tracing, labeled classical cells stably maintained glandular morphology and expression of key classical differentiation markers (**Fig. 2F**). We applied the same continuous state-space analysis to scRNA-seq profiles of lineage-traced cells from *KPF-classical* tumors. In contrast to basal-derived cells, classical-lineage cells remained concentrated near the classical pole over the 14-day tracing interval, with limited redistribution toward basal, mesenchymal, or intermediate identities (**Fig. 2H**). Collectively, these findings indicate that classical PDAC cells represent a subset of malignant cells with a relatively fixed, epithelial identity and limited plasticity when compared to the basal PDAC cells.

### Selective ablation of the basal state leads to rapid collapse of PDAC tumors

We next used the *KPF-basal* model to move from lineage potential to function, asking what happens to an established PDAC tumor when the basal state is selectively removed. Basal cells in *KPF-basal* PDAC tumors were ablated by repeated systemic administration of diphtheria toxin (DT) once tumors became palpable (approximately 7 weeks of age) (**Fig. S2A**). Strikingly, basal state ablation induced rapid and profound tumor regression as measured by whole-animal magnetic resonance imaging (*21*) and with pancreas weights returning to the range of age-matched non-tumor-bearing controls within 7 days (**Fig. 3A; Fig. S2B**).

**Figure 3.**
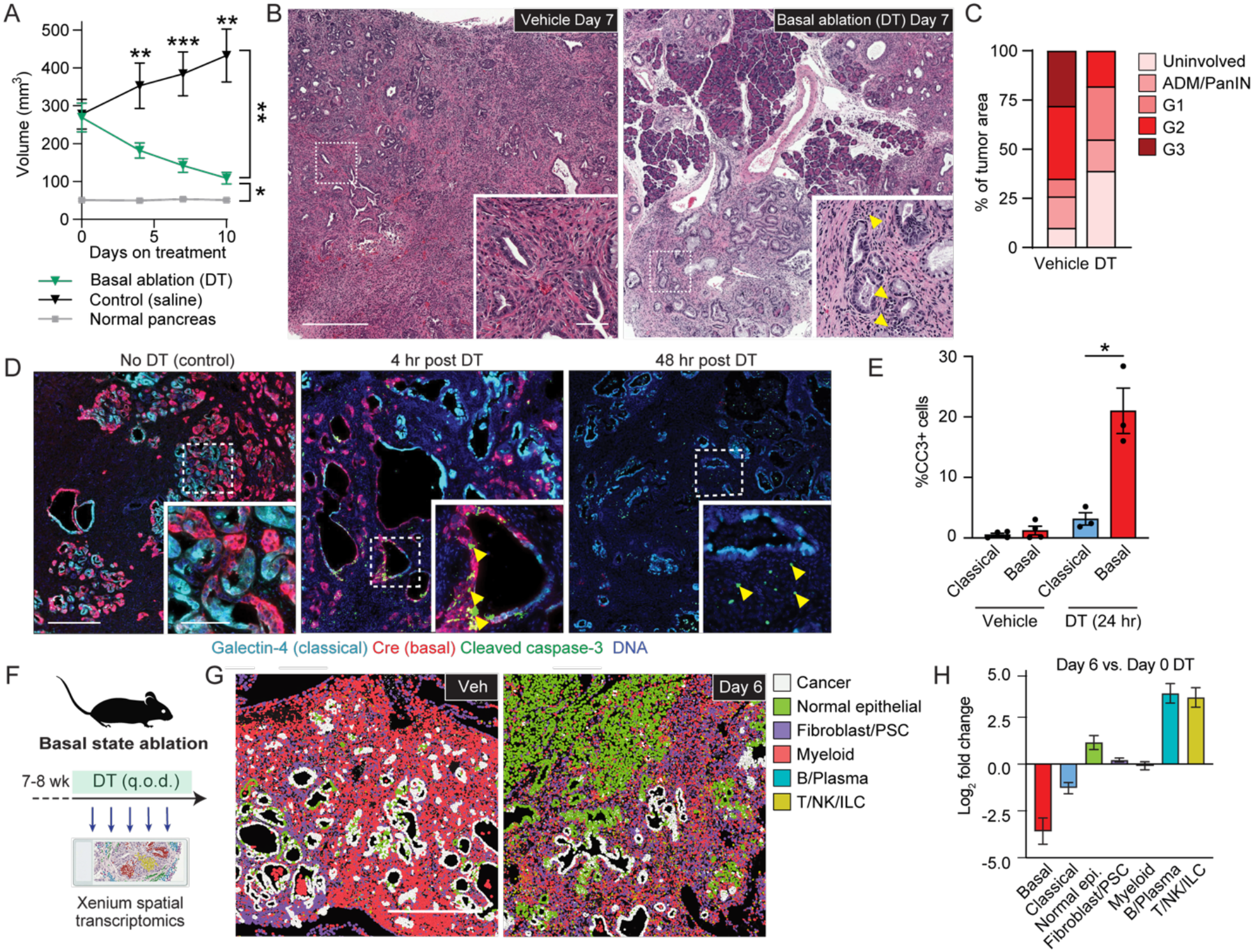
Selective ablation of the basal cell state leads to rapid remodeling of the tumor ecosystem. **(A)** Mean tumor volume in vehicle- and DT-treated basal reporter mice (*n* = 5–7 mice per group), as measured by MRI. Error bars, s.e.m.; *, *p* < 0.05; **, *p* < 0.005; ***, *p* < 0.0005, Welch’s t-test. **(B)** Representative H&E staining of untreated and 1-week DT-treated *KPF-basal* tumors. Scale bar, 1 mm. Inset, 50 μm. Arrowheads indicate lymphocyte infiltration. **(C)** Percentage of H&E-stained tissue assigned to each histologic category. *n* = 3 biological replicates per condition. **(D)** Immunofluorescence staining for cleaved Caspase-3 (apoptotic cells) and cancer state markers in untreated *KPF-basal* tumors and tumors collected 4 or 48 hr after DT treatment. Arrowheads, cleaved Caspase-3⁺ cells. Scale bar, 200 µm. Inset, 40 µm. **(E)** Mean percentage of cancer cells that are cleaved Caspase-3⁺ in untreated (*n = 4*) or 24 hr post DT (*n = 3*) tumors. Error bars, s.e.m.; *, *p* = 0.034, Welch’s *t*-test. **(F)** Experimental design for DT-mediated ablation of *Slc4a11-MCD*⁺ cells in *KPF-basal* tumors. Xenium spatial transcriptomics was performed before DT administration and 2, 4, and 6 days after initiation. **(G)** Representative Xenium data showing tissue remodeling by day 6 after DT treatment, including an enrichment of infiltrating T, NK and normal pancreatic cells compared to vehicle (Veh) control. Polygons represent 10X cell segmentation boundaries colored by cell type, as determined by clustering, and cell type annotation (Methods). **(H)** Log_2_-fold changes in tumor cell-state composition at day 6 versus baseline following DT-mediated basal cell ablation. Fold changes are computed from the fraction of each cell state out of all parenchymal cells in each sample.

Residual disease following basal cell state ablation exhibited a shift toward lower-grade (G) histology, with G2/G3 adenoma/adenocarcinoma regions dramatically reduced in size and an enrichment of G1 (pancreatic intraepithelial neoplasia, PanIN) or earlier lesions (**Fig. 3B,C)**. Notably, basal state-ablated tumors demonstrated robust infiltration by lymphocytes that was immediately obvious in H&E sections—cells that are largely excluded in PDAC tissues (**Fig. 3B**, arrowheads).

To confirm specificity of DT-mediated suicide gene activation to the basal state, we assessed apoptosis over the course of ablation. Early after the first DT dose (∼4 hours), apoptosis was confined to the basal cells, whereas classical and mesenchymal cancer cells as well as adjacent non-epithelial cells were spared, confirming selective targeting (**Fig. 3D,E; Fig. S3C,D**). By 48 hours, basal cell elimination was complete. Interestingly, at this time point, we observed a secondary wave of apoptosis within neighboring stromal populations and other cancer cell states that was not seen in early time points, suggesting this wave of cell death was secondary to the loss of the basal cell state (**Fig. 3D**). Importantly, DT-mediated cell death has previously been shown to be minimally immunogenic and lacking in bystander toxicity (*22, 23*). Together, these findings indicate that the basal cell state is required for the maintenance of the malignant and stromal components of the PDAC tumor ecosystem.

To test whether tumor maintenance is uniquely dependent on the basal state, we directly compared ablation of the basal and classical cell states. We generated parallel cell lines from autochthonous murine *KPF-basal* and *KPF-classical* tumors and conducted orthotopic transplantation into syngeneic hosts to generate parallel cohorts and compare state-specific requirements. Despite similar baseline cell state composition (**Fig. 1C,F**), selective ablation of classical cells did not induce tumor regression, whereas basal state ablation produced deep responses that paralleled our findings in the autochthonous PDAC tumors (**Fig. S2E-H**). These findings establish a unique requirement for the basal state in tumor maintenance in orthogonal model systems.

### Basal cell state ablation leads to rapid stromal remodeling and cytotoxic lymphocyte influx

The rapid regression and immune infiltration observed after seven days of basal state ablation suggested that selective loss of basal cells initiates a broader tissue-level remodeling response. To define this response over time, we collected tumors at days 2, 4, and 6 on continuous DT and profiled them by Xenium spatial transcriptomics using a custom 480-gene panel capturing cancer cell state identity, immune activation states, cognate cytokine–receptor pairs, and key pathway activity markers resolved in our single-cell reference data (**Fig. 3F, Table S1, Methods**). High-quality spatial transcript segmentation using Segger (*24*) enabled robust spatially resolved cell phenotyping of individual cells, allowing us to reconstruct state-rich tissue maps across the ablation time course rather than reducing the response to broad cell type abundance alone. Given substantial phenotypic heterogeneity in the mesenchymal cancer state, as well as variability in fraction of mesenchymal cells in the autochthonous tumors, we excluded mesenchymal cancer cells from the cell state analysis and focused on the classical and basal malignant compartments and on stromal cells.

This analysis revealed dramatic dynamic remodeling of the tumor ecosystem in response to basal cell state ablation. At baseline, tumors recapitulated stereotypic PDAC architecture described in multiple human and mouse PDAC atlas studies (*25, 26*), with abundant cancer cells, CAFs, and tumor-associated macrophages (TAMs), and sparse lymphocyte infiltration. By day 6, this architecture was profoundly altered: the tumors showed marked cancer cell depletion, enrichment of residual early-stage neoplastic lesions and normal acinar cells, and prominent cytotoxic lymphocyte infiltration within the tumor bed (**Fig. 3F-H**). Basal cancer cells were rapidly depleted, whereas classical epithelial cells persisted and eventually became the dominant residual neoplastic compartment (**Fig. 3G, H**), consistent with *de facto* eradication of adenocarcinoma in the pancreas. Together, these findings show that basal-state ablation triggers a tissue-wide response that extends well beyond the targeted cells, coupling collapse of aggressive malignant compartments with stromal remodeling and cytotoxic lymphocyte influx.

### A classical–basal tumor cell axis organizes tumor microenvironment niches in PDAC

The rapid and profound remodeling of the tumor microenvironment and lymphocyte influx triggered by basal cell state ablation suggested that basal cancer cells may actively instruct immunosuppressive stromal features. We therefore asked whether basal and classical cancer cells occupy distinct local ecosystems in PDAC. Addressing this requires a representation that preserved the measured transcriptional richness of each neighborhood rather than reducing it to cell type counts (*27*). We defined cancer cell-centered spatial units as local tissue neighborhoods anchored on individual cancer cells, retaining for each unit the spatially defined expression matrix of 32 neighboring cells in physical space captured by Xenium rather than collapsing the neighborhood to summary statistics (**Fig. 4A**).

**Figure 4.**
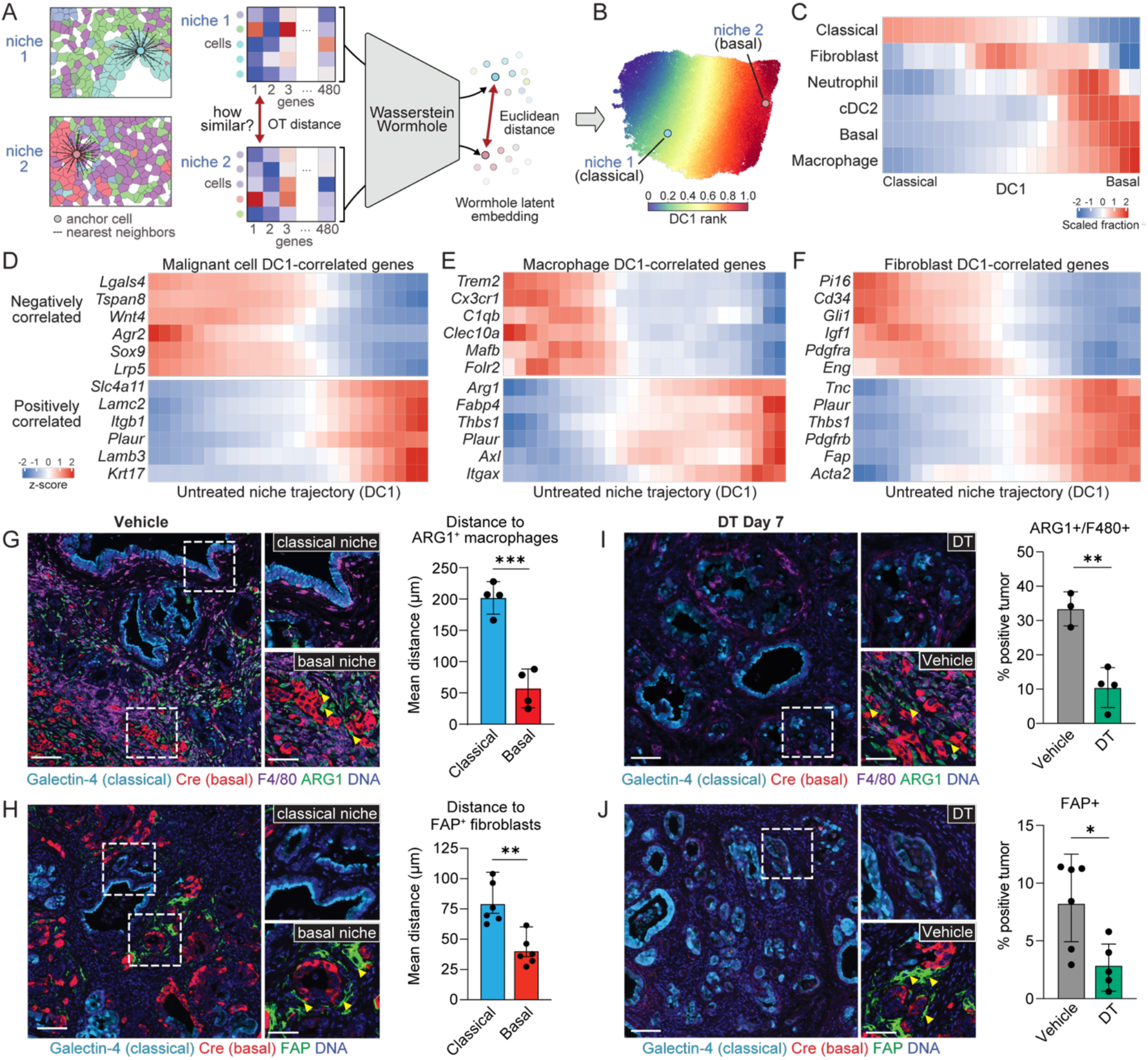
A classical–basal tumor cell axis organizes tumor microenvironment niches and is disrupted by basal cell ablation. **(A)** Schematic of spatial neighborhood analysis using Xenium spatial transcriptomics data. The local tissue niche of each cancer cell is defined as the full gene expression matrix of the 32 most proximal cells around that anchor cancer cell. The Wasserstein Wormhole transformer embeds each niche matrix as a point in a latent space wherein nearby points correspond to neighborhoods with similar transcriptional niche states. Niche similarity in the Wormhole latent is determined by Euclidean distance between points—an easily computable, scalable metric compatible with existing single-cell analysis tools such as diffusion component analysis. **(B)** Diffusion component analysis identified the dominant axis of niche variation (DC1) in untreated *Slc4a11-*MCD tumors as separating classical-anchored (classical niche) from basal-anchored (basal niche) tumor neighborhoods (Methods). **(C)** Coarse cellular composition of spatial neighborhoods ordered along the classical–basal niche axis (DC1). **(D)** Cancer cell genes associated with DC1, highlighting canonical epithelial, basal-state, matrix-adhesion, and invasive program genes (Methods). **(E)** Macrophage genes associated with DC1, highlighting tissue resident (classical) or immunosuppressive and wound healing (basal) features. **(F)** Fibroblast genes associated with DC1, highlighting tissue resident and inflammatory CAF (classical) or myofibroblast and matrix remodeling (basal) features. **(G)** Immunofluorescence staining and quantification of average distance from Galectin-4^+^ or *Slc4a11*-Cre^+^ cells to the nearest Arg1^+^ F4/80^+^ macrophages in untreated *Slc4a11*-MCD tumors. Scale bar, 100 μm. *n* = 4; error bars, s.d.; ***, *p* = 0.0004, Welch’s *t*-test. **(H)** Immunofluorescence staining and quantification of average distance from Galectin-4^+^ or *Slc4a11*-Cre^+^ cells to the nearest FAP^+^ fibroblasts in untreated *Slc4a11*-MCD tumors. Scale bar, 100 μm. *n* = 6; error bars, s.d.; **, *p* = 0.001, Welch’s *t*-test. **(I)** Immunofluorescence staining and quantification of Arg1^+^ F4/80^+^ cells in *Slc4a11*-MCD tumors 7 days after treatment with vehicle (*n* = 3) or DT (*n* = 4). Scale bar, 100 μm. Error bars, s.d.; **, *p* = 0.003, Welch’s *t*-test. **(J)** Immunofluorescence staining and quantification of FAP^+^ cells in *Slc4a11*-MCD tumors 7 days after treatment with vehicle (*n* = 6) or DT (*n* = 5). Scale bar, 100 μm. Inset, 60 μm. Error bars, s.d.; *, *p* = 0.018, Welch’s *t*-test.

Single-cell genomics became powerful because cells could be placed in a phenotypic manifold where distance reflected biological relatedness. We sought the analogous representation for tissue neighborhoods. Wasserstein Wormhole creates such a niche geometry by embedding spatial units into a shared space in which nearby points represent local multicellular ecosystems with similar transcriptional and cellular organization (**Methods**)(*28*). This allowed us to visualize, compare, and order PDAC niches by their overall tissue state, preserving the rich transcriptional context, continuous gene programs, and multicellular organization of each neighborhood rather than relying on discrete cell labels alone (**Fig. 4A,B**).

We next used the Wormhole niche map to ask which biological features explain the largest differences among local PDAC neighborhoods. Diffusion component analysis (*29*) revealed that the dominant axis of variation, DC1, corresponded to a classical-to-basal niche axis: spatial units anchored on classical cancer cells occupied one end of the axis, whereas basal-anchored niches occupied the other (**Fig. 4B-D; Fig. S3A,B**). This organization indicates that malignant cell state is not merely a property of individual cancer cells but is tightly coupled to the surrounding stromal and immune context (**Fig 4A,B**).

We next examined the cellular composition of neighborhoods along the classical–basal niche axis. Classical neighborhoods were epithelial-rich, containing classical cancer cells and acinar, ductal, and acinar-to-ductal metaplasia-like (*30*) compartments consistent with a comparably more differentiated and benign tumor tissue architecture, often surrounded by fibroblasts (**Fig. 4C; Fig. S3C**). In contrast, basal neighborhoods were stromal-rich, enriched for macrophages and monocytes (hereafter, macrophages; Methods), other myeloid cells, fibroblasts, and endothelial cells (**Fig. 4C**; **Fig. S3C, D**). Thus, the dominant niche axis separates epithelial-rich, differentiated classical neighborhoods from stromal-rich basal neighborhoods.

We next used this ordered niche axis to move beyond cell type composition and interrogate how the functional programs of niche-resident cells vary across the classical–basal continuum. Within cancer cells, macrophages, and fibroblasts, we identified genes most strongly associated with DC1 and displayed representative markers in the main figure, with full ranked lists provided in the supplement (**Fig. 4D-F; Fig. S3E-G; Table S3)**. This revealed coordinated remodeling across all three compartments, with cancer cell identity, macrophage polarization, and fibroblast activation shifting progressively along the same niche axis.

In cancer cells, the classical pole was defined by canonical classical state markers, including *Lgals4*, *Agr2*, and *Tspan8*, together with differentiated epithelial programs marked by *Sox9* (**Fig. 4D; Fig. S3E**). This pole also showed evidence of an active WNT program, including *Wnt4* and *Lrp6*, consistent with our recent work showing that spatially organized WNT signaling helps drive classical cell differentiation in PDAC (*31*). In contrast, progression toward the basal pole was marked by expression of human-conserved basal state genes, including *Krt17 and Lamc2*—as well as *Slc4a11*, the marker used in our basal reporter and recovered here through an unbiased niche-level analysis. Importantly, this shift was accompanied by increasing expression of laminins, integrins (*Lamb3, Itgb1, Itga5*), and *Plaur*—typically associated with matrix engagement, adhesion, and invasion (**Fig. 4D; Fig. S3E**). Thus, the classical-basal niche axis captures an active transition from a differentiated epithelial identity toward a plastic and invasive malignant program.

Macrophages similarly underwent a pronounced state shift along the classical–basal niche axis. Classical-proximal macrophages expressed tissue-resident features marked by *Folr2* and *C1qb*, whereas basal-proximal macrophages progressively acquired immunosuppressive and wound-healing programs marked by *Arg1*, *Axl*, *Thbs1*, and *Plaur* expression ((*32*) **Fig. 4E; Fig. S3F**). Fibroblasts showed a parallel remodeling pattern along the classical–basal niche axis. Classical-proximal fibroblasts were enriched for tissue-resident programs (*Pi16*, *Gli1*) and inflammatory CAF (iCAF)-like features (*Pdgfra*, *Igf1)*—consistent with CAF programs that have been linked to epithelial-derived cytokine signaling in PDAC (*33–35*). In contrast, basal-proximal fibroblasts expressed activated myofibroblast and matrix-remodeling genes, including *Acta2*, *Fap*, *Tnc*, *Thbs1*, and *Plaur* (**Fig. 4F; Fig. S3G**). Thus, progression toward the basal niche is accompanied by a coordinated shift from inflammatory/tissue-resident fibroblast programs toward myofibroblast activation and ECM remodeling. Notably, *Plaur* was enriched at the basal pole across cancer cells, macrophages, and fibroblasts, suggesting a convergent pro-invasive and ECM-remodeling program within the basal niche ecosystem.

### Basal cell state ablation dismantles basal-associated stromal niches

Given the tight spatial coupling between basal cancer cells and immunosuppressive stromal populations, we next asked whether these stromal niches require the continued presence of basal cells. Immunofluorescence confirmed that, at baseline, basal tumor regions were enriched with FAP⁺ myofibroblastic CAFs—a matrix-remodeling fibroblast state associated with PDAC desmoplasia (*33–35*)—as well as ARG1⁺ macrophages, confirming the basal-associated stromal architecture identified by spatial transcriptomics (**Fig. 4G,H**). Following basal state ablation, both stromal populations were markedly reduced (**Fig. 4I,J**), indicating that these niches are not merely spatially associated with basal cells but are actively maintained by them.

Consistent with this targeted immunofluorescence, Xenium analysis revealed broader remodeling of the stromal compartment after basal state ablation. The myofibroblastic and matrix-remodeling CAF states were depleted, whereas inflammatory, tissue-resident, and interferon (IFN)-high fibroblast states were enriched. Macrophages showed a depletion of suppressive ARG1^+^ phenotypes and an enrichment of an interferon (IFN)-high inflammatory state and tissue resident states (**Fig. S4A**). This remodeling was specific to loss of the basal state: in parallel orthotopic *KPF-basal* and *KPF-classical* models, basal cell ablation reduced ARG1⁺ macrophages, whereas classical cell ablation did not dismantle the same stromal programs and instead was associated with relative enrichment of ARG1⁺ macrophages (**Fig. S4B-G**). Together, these data indicate that basal cancer cells are required to maintain the immunosuppressive and fibrotic niche that surrounds them, establishing the basal state as an active organizer—not merely a marker—of key pro-tumorigenic components of the PDAC tumor ecosystem.

### The basal cancer cell state is a source of key immunosuppressive cytokines in PDAC

The enrichment of activated myofibroblasts and immunosuppressive myeloid cells around basal cancer cells, coupled with their rapid depletion after basal state ablation, suggested that basal cells may actively sustain their local TME through paracrine signaling. We therefore asked which cancer-derived cytokines and ligands were positioned to organize basal-associated stromal niches. To identify candidate mediators, we focused on cancer-derived cytokines and ligands that satisfied two spatial criteria. First, we prioritized ligands whose expression increased toward the basal pole of the classical–basal niche axis—reasoning that these factors may help drive the signaling programs that organize basal- vs. classical-associated niches. Second, we asked whether ligand expression in cancer cells coincided locally with enrichment of stromal or immune cells expressing the corresponding receptor (**Fig. S5A,B**). This spatial analysis revealed a basal-associated ligand module that included *Csf2*, *Lif*, *Tgfb1*, and *Pdgfb—*factors with established roles in myeloid and fibroblast regulation (*36–39*). By contrast, classical cells expressed a distinct ligand module, including *Il18* and *Wnt4* (**Fig. S3E, S5C**). These data indicate that basal and classical states differ not only in intrinsic malignant programs, but also in the paracrine signals they use to shape their local tissue ecosystems.

We next asked whether these basal-associated ligands showed evidence of local engagement with neighboring stromal populations. Among these signaling pairs, the *Csf2–Csf2ra* interaction showed particularly strong spatial support: *Csf2* expression in cancer cells was associated with enrichment of *Csf2ra*⁺ expressing macrophages and cDC2s in the same niches, together with induction of GM-CSF response and feedback genes, including *Socs2* and *Cish* (**Fig. 5A, B**) (*36, 37*). Macrophage *Mki67* was also increased in *Csf2*-high niches, suggesting *Csf2* and other basal-derived factors drive local macrophage expansion (**Fig. 5A, B**). Basal niches also showed increased *Stat3* expression, consistent with a stromal signaling context responsive to cytokines such as *Lif* and *Csf2*. In parallel, *Lif* and *Tgfb*-high associated niches showed receptor enrichment and pathway activity in neighboring stromal populations, nominating multiple basal-derived signaling pathways that may sustain the suppressive niche (**Fig. 5A**).

**Figure 5.**
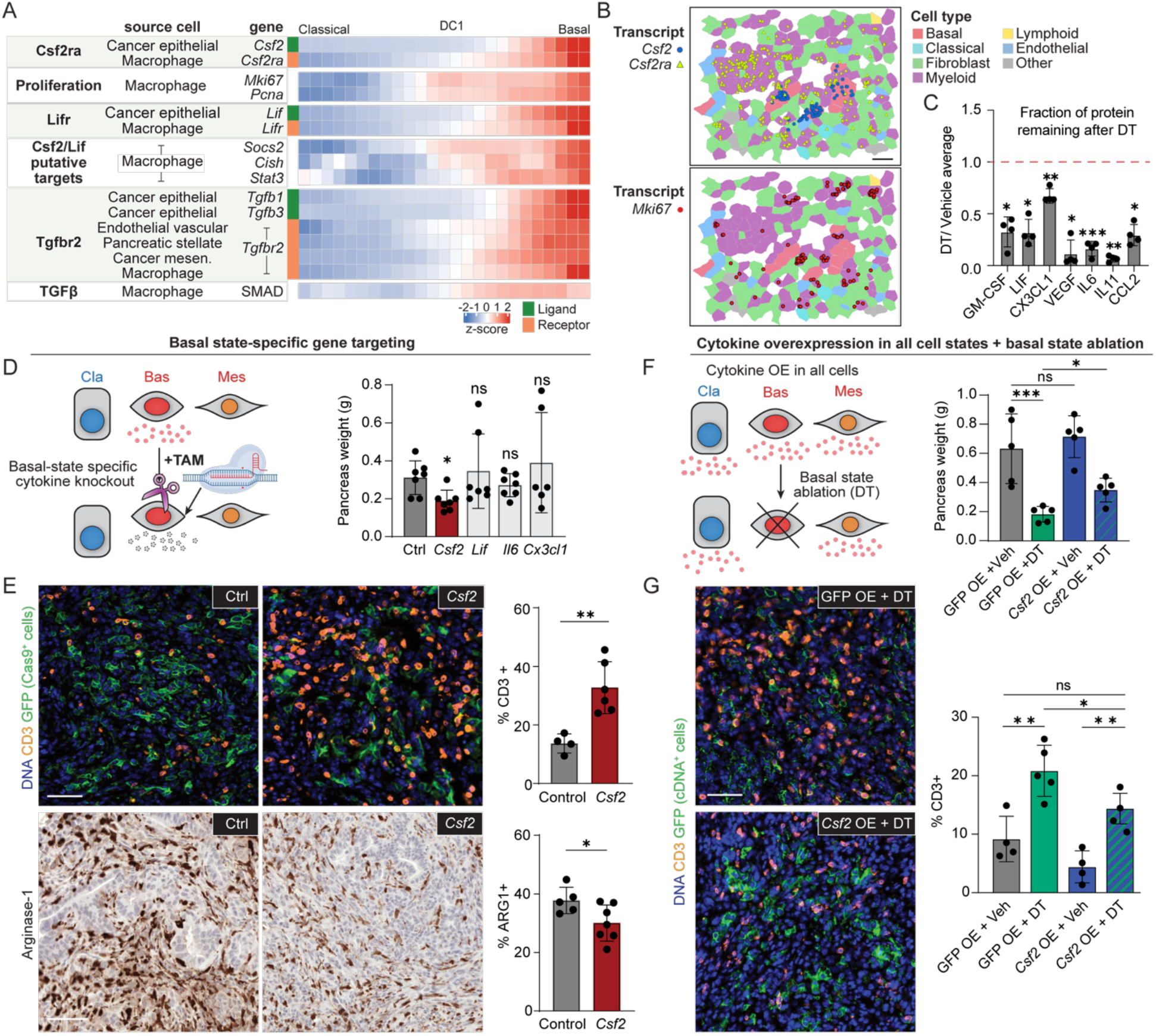
GM-CSF produced by basal cancer cells enforces an immunosuppressive niche. **(A)** Putative signaling interactions defined by cognate ligand–receptor pairs expressed in different cell types within basal niches, as well as associated expression programs. SMAD, average scaled expression of broadly induced TFGβ response genes (*Pmepa1*, *Skil*, *Smad7*, *Serpine1*). **(B)** Representative Xenium images showing an example of likely *Csf2* signaling from basal cells to neighboring *Csf2ra*-expressing myeloid cells, and activation of myeloid proliferation (*Mki67*) within basal niches. **(C)** Mean cytokine concentration fold-change in orthotopic *Slc4a11*-MCD tumors 7 days after DT (*n* = 4) or vehicle (*n* = 3) treatment. Error bars, s.d.; *, *p* < 0.05; **, p < 0.01; ***, p < 0.001, Welch’s *t*-test. **(D)** Left, *in vivo* basal-state-restricted CRISPR perturbation system: TAM administration activates CreER^T2^, and thereby Cas9 expression, only in *Slc4a11-MCD*⁺ cells, such that genome editing of targeted loci occurs selectively in basal cells. Right, mean pancreas weight at experimental endpoint following basal-state-restricted sgRNA perturbation. *n* = 6–7 biological replicates for each sgRNA; error bars, s.d.; *, *p* = 0.011; ns, *p* > 0.05, Welch’s t-test. **(E)** Immunohistochemical staining and quantification of CD3^+^ and Arg1^+^ cells in *sgCtrl* (*n* = 4) and *sgCsf2* (*n* = 6) tumors. Scale bars, 50 μm. Error bars, s.d.; *, *p* = 0.0326; **, *p* = 0.0021, Welch’s *t*-test. **(F)** Left, system for cytokine overexpression combined with basal-state ablation: *Csf2* overexpression cassette is delivered to all cancer cells via Adeno-Cre infection of orthotopic *Slc4a11-MCD* tumors, for constitutive cytokine expression during DT-mediated depletion of *Slc4a11-MCD*⁺ basal cells. Right, Mean pancreas weight of control GFP or *Csf2* overexpressing (OE) with vehicle (Veh) or basal-state ablated (DT) tumors at experimental endpoint. *n* = 5 tumors per condition; error bars, s.d.; *, p = 0.012; ***, *p* = 0.002; ns, *p* > 0.05, Welch’s t-test. **(G)** Immunofluorescence staining and quantification of CD3^+^ in tumors in (F). Scale bar, 50 μm. *n* = 5 in each GFP OE condition, *n* = 4 in each *Csf2* OE condition; error bars, s.d.; *, *p* < 0.05; **, p < 0.01; ns, *p* > 0.05, Welch’s *t*-test.

Because cytokine transcripts are often sparse in spatial transcriptomic data, and because our targeted Xenium panel may not capture the full spectrum of basal-derived ligands, we used an orthogonal protein-based approach to directly measure cytokines lost after basal cell ablation in PDAC tumors. We generated orthotopic *KPF-basal* transplant tumors in syngeneic, immunocompetent mice, ablated the basal state in established tumors, and performed 44-plex cytokine profiling on whole tumors after 7 days of continuous depletion, comparing to control-treated tumors (**Fig. 5C; Fig. S6A,B**). Multiple cytokines were robustly reduced after basal cell ablation, including GM-CSF (encoded by *Csf2*), LIF, IL-6, CX3CL1, VEGF, CCL2, and IL-11. However, the strongest changes did not simply mirror the spatial nomination. When integrated with the spatial ligand–receptor analysis above, this depletion profile converged most strongly on GM-CSF and LIF, which were both enriched in basal-associated niches, locally coupled to receptor-positive stromal populations, and lost after basal-state ablation.

Together, these complementary spatial and protein-level data nominated GM-CSF and LIF as candidate basal-derived cytokines positioned to sustain the immunosuppressive TME. We therefore prioritized *Csf2* and *Lif* as convergent candidates for functional interrogation by gain- and loss-of-function approaches and included *Il6* and *Cx3cl1* as top ablation-sensitive cytokines identified by protein profiling but not nominated by the spatial ligand–receptor analysis.

### GM-CSF produced by basal cancer cells enforces an immunosuppressive niche

To directly test the function of basal-derived cytokines, we built a basal state-specific CRISPR knockout system. We leveraged the CreER activity built into the *KPF*-*basal* (*Slc4a11*-MCD) reporter and coupled it to a Cre-dependent Cas9 cassette, thereby restricting Cas9 expression to basal cancer cells (**Fig. S6C**). For each candidate cytokine, we introduced a multiplexed sgRNA construct encoding four guides against that gene to maximize knockout efficiency (**Fig. S6D)** (*40*). We validated this approach *in vitro*, confirming robust loss of GM-CSF and CX3CL1 by ELISA (**Fig. S6E**) and genomic disruption of additional cytokine loci by targeted sequencing. This approach enabled precise, inducible deletion of cytokines from the basal state *in vivo*, while leaving other cancer cell states unedited, allowing us to test whether individual factors produced by the basal state are required to maintain the surrounding tumor ecosystem.

We next used this platform to test the functional consequences of deleting individual candidate cytokines from basal cancer cells *in vivo* (**Fig. S6F**). To do this, we selectively deleted candidate cytokines from basal cancer cells in established orthotopic tumors harboring our basal state-specific editing system. Among the cytokines tested, only basal-restricted deletion of *Csf2* produced a robust tumor phenotype. Strikingly, basal-restricted knockout of *Csf2* reduced tumor burden by 39.4% ± 18.0% compared with controls (**Fig. 5D**). This was accompanied by local tumor remodeling: edited tumor regions showed increased T cell infiltration and reduced ARG1⁺ myeloid cells (**Fig. 5E**).

We next asked whether GM-CSF was sufficient to restore features of the suppressive niche after basal cell loss. To do this, we constitutively expressed *Csf2* cDNA in all cancer cells while ablating the basal cancer cell state (**Fig. S6G-I**). *Csf2* overexpression partially rescued tumor growth after basal cell ablation (**Fig. 5F**). These tumors also had fewer infiltrating T cells, both at baseline and after basal cell ablation, suggesting that GM-CSF limits the immune activation typically induced by loss of basal cells (**Fig. 5G**).

Together, these loss- and gain-of-function data identify basal-derived GM-CSF as a key organizer of the local immunosuppressive niche. Deleting *Csf2* specifically from basal cancer cells reduced Arg1⁺ myeloid cells, increased T cell infiltration, and impaired tumor growth, while restoring GM-CSF partially rescued tumor growth and suppressed T cell infiltration after basal cell ablation. Thus, we find that a single basal-derived cytokine is sufficient to drive a cell circuit within the PDAC TME that limits T cell infiltration and supports cancer cell growth *in vivo*.

### Basal state ablation reprograms residual classical niches for T/NK cell–mediated tumor killing

We next asked how selective loss of the basal cells propagates into a broader anti-tumor response against cancer cells that are not directly targeted by DT (**Fig. 6A,B**). Basal state ablation rapidly depleted the basal cancer state, leaving classical epithelial cells as the dominant residual malignant compartment (**Fig. 6B**; **Fig. 3H**). Yet these residual classical cells did not remain in their original niche context: over the ablation time course, classical cell neighborhoods progressively shifted from cancer cell–dominated niches to tissue states containing increasing numbers of T/NK cells and other immune cell types **(Fig. S7A)**. Because classical cells are not directly ablated, these changing classical niches provided a window into the non–cell-autonomous tissue response triggered by basal-state loss.

**Figure 6.**
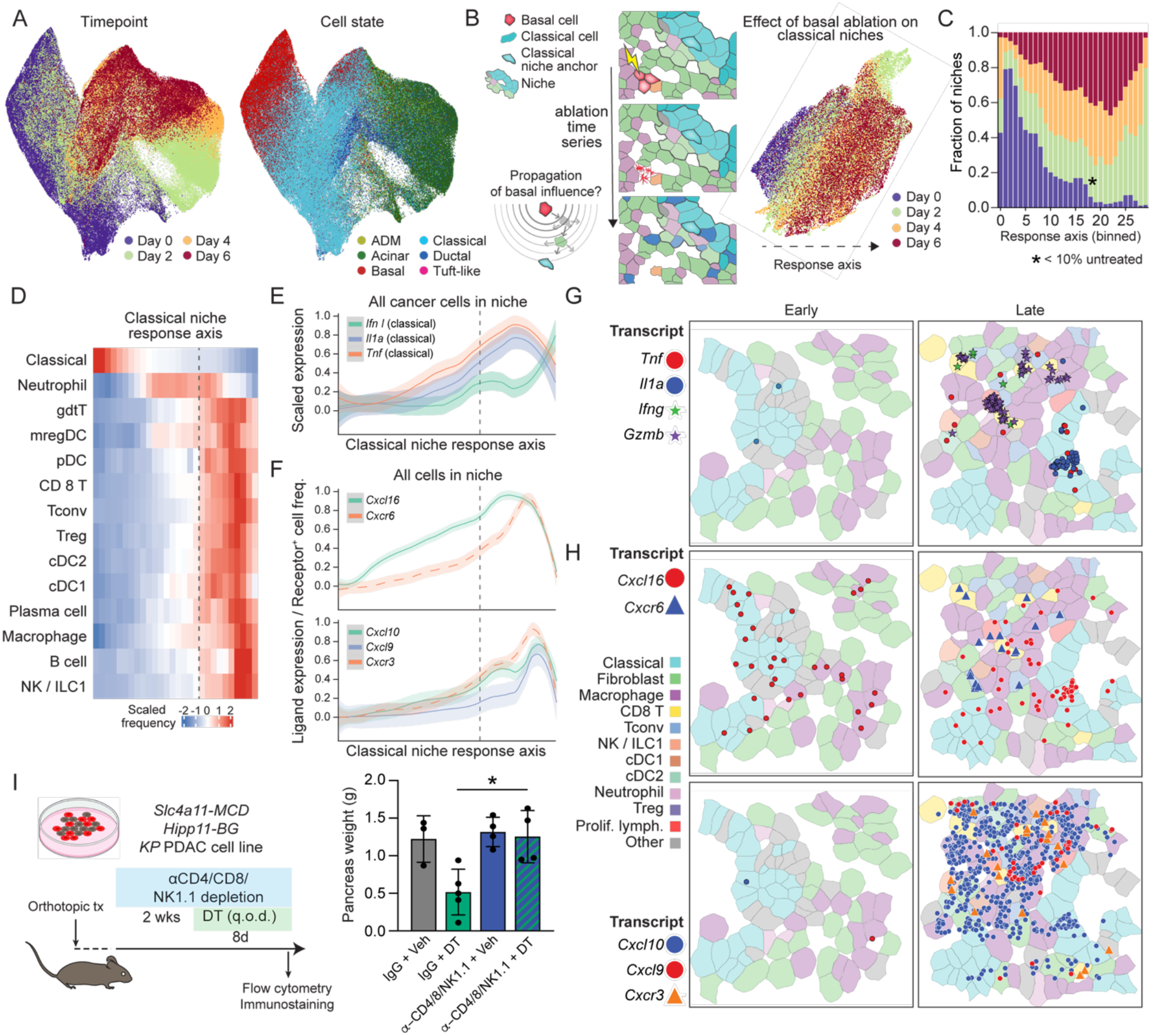
Basal state ablation induces sequential immune recruitment and T/NK cell-mediated tumor killing. **(A)** UMAP embedding of all niches in basal state ablation experiment, labeled by anchor cell state (right) or timepoint (left) before (day 0) or after (days 2–6) DT administration. **(B)** Left, schematic of rationale for assessing the role of basal cells on long-range tumor remodeling, by querying classical-anchored niches across the basal ablation time series. Right, UMAP embedding of classical niches only (Methods), colored by timepoint, which illustrates a temporally ordered classical niche response axis. **(C)** Contribution of ablation timepoints to each bin along the classical niche response axis (niche anchors colored as in (B)). Asterisk denotes the bin at which the untreated fraction drops below 10% of niche bin fraction. **(D)** Cell type composition (*z*-scaled per cell type) along the classical niche response axis. Dotted line corresponds to asterisk in (C). **(E)** Scaled per-bin expression of select cancer-cell-expressed damage-associated ligands, ordered along the classical niche response axis. Dotted line corresponds to asterisk in (C). **(F)** Scaled per-bin expression (for ligands, solid lines) or positive cell fraction (for cognate receptors, dashed lines) of chemokine–ligand receptors that are positively associated with the classical niche response axis. Dotted line corresponds to asterisk in (C). **(G,H)** Selected annotated Xenium fields of view for early (bins 4–6) and late (bins 23–26) response timepoints. Raw transcript localizations (points) of select effector molecules (G) and chemokine–receptor pairs (H) show higher expression and colocalization in later temporal bins. **(I)** Experimental design for combined lymphocyte depletion and DT-mediated basal-state ablation (left) and mean pancreas weight at experimental endpoint (right). *n* = 3–5 tumors for each condition; error bars, s.d.; *, *p* = 0.0315, Welch’s t-test.

Because basal state ablation unfolds asynchronously across each tissue, the time course contains a heterogeneous mixture of directly ablated basal regions, mixed neighborhoods undergoing local collapse, and residual classical tumor regions undergoing secondary remodeling. To isolate this secondary response from the direct local effects of basal cell elimination, we focused on highly classical-enriched niches across the ablation time course, retaining only niches in which the malignant compartment was nearly entirely classical, yielding approximately 50,000 classical niches for analysis (**Fig. 6B, Methods**). We then asked whether these classical niches followed stereotyped, temporally ordered remodeling path after basal state ablation. Wormhole analysis of these niches transformed the heterogeneous ablation time course into an ordered tissue trajectory: diffusion analysis identified a dominant axis through this space that closely tracked time after ablation and ordered classical niches from their pre-ablation state to progressively immune-infiltrated post-ablation states (**Fig. 6B,C**; **Fig. S7A**). We refer to this ordering as the “classical response axis,” (**Methods**) which provided the framework for identifying the sequential cell recruitment, signaling, and effector programs that emerge within residual classical niches after basal-state loss.

Having defined an ordered trajectory of classical niche remodeling, we next asked which immune populations appeared as niches progressed along the classical response axis. Immune influx occurred in a clear sequence: myeloid populations, including neutrophils, macrophages, and CCR7^+^ dendritic cells (mregDCs), appeared early, followed by cDC1 and cDC2, cytotoxic T cells, NK cells, ILC1-like populations, and later B and plasma cells (**Fig. 6D**). Thus, basal cell ablation triggers a staged immune recruitment within the residual classical tumor niche.

We next used the classical response axis to ask which signals accompany the earliest stages of this immune remodeling. Projecting cytokines, chemokines, and ligand–receptor pairs onto the axis (**Methods**) revealed an early injury-associated inflammatory program: residual classical cancer cells upregulated *Tnf*, *Il1a*, *Areg*, and type I interferon-associated signals, while fibroblasts induced the alarmin *Il33* (**Fig. 6E,G; Fig. S7B,C**). These signals suggest that basal cell loss is sensed as a local tissue injury—triggering alarmin release, NF-κB-driven inflammatory signaling, epithelial repair programs, and type I interferon responses that support antigen presentation and immune priming (**Fig. S7D**). This response coincided with the dissolution of the pre-existing tumor niche architecture. Basal-associated macrophage states, including *Arg1*⁺, *Axl*^+^, and *Itgax*⁺ programs, declined along the response axis, while macrophages and fibroblasts acquired increasingly inflammatory and IFN-polarized features (**Fig. S7D,E)**. Thus, basal cell ablation dismantles suppressive stromal programs and reprograms the residual classical niche into an inflammatory response state poised for immune recruitment.

Having identified an early injury–inflammatory program, we next asked how this activated niche recruits immune cells. The classical response axis allowed us to distinguish chemokines that merely changed after ablation from those whose timing and location were consistent with immune recruitment. We therefore prioritized chemokine programs in which ligand induction preceded or coincided with the influx of cognate receptor-positive immune cells along the trajectory. In parallel, we statistically tested local spatial coupling by asking whether niches with higher ligand abundance were enriched for neighboring cells expressing the cognate receptor (**Fig. 6F, Fig. S8A-C, Methods**). Chemokine axes supported by both temporal ordering and significant local ligand–receptor association were nominated as candidate mediators of immune recruitment.

This analysis nominated a sequential chemokine program for lymphocyte recruitment. Early in the response, *Cxcl16* expression by residual cancer cells preceded accumulation of *Cxcr6*⁺ lymphocytes, consistent with entry of tissue-resident or tissue-tropic T cell populations into the remodeling classical niche (**Fig. 6F,H; Fig. S8A,B**). At later stages, *Cxcl9* and *Cxcl10* increased together with *Cxcr3*⁺ immune cells, nominating the canonical *Cxcl9/Cxcl10–Cxcr3* axis as a major recruitment program for activated cytotoxic T cells, NK cells, and cDC1s (**Fig. 6F,H)** (*41*). These chemokine changes were accompanied by the appearance of activated effector populations: late response niches contained increased CD8⁺ and CD4⁺ T cells and NK cells expressing *Gzmb*, *Prf1*, *Ifng*, and *Tbx21*, together with late-peaking interferon response programs (**Fig. S8C; Fig. S9A,B**). Thus, the classical response axis resolves a progression from inflammatory niche remodeling to chemokine-associated lymphocyte recruitment and cytotoxic effector activation.

The trajectory analysis predicted that basal state ablation culminates in cytotoxic lymphocyte recruitment and activation. To test whether this immune response was required for tumor regression, we depleted CD4⁺ T cells, CD8⁺ T cells, and NK cells during DT-mediated basal state cytoablation (**Fig. 6I; Fig. S9C-G)**. Combined lymphocyte depletion blunted the tumor-regressive effect of basal state ablation and rescued tumor growth (**Fig. 6I**). These results establish that tumor collapse after basal cell ablation is not simply a direct consequence of eliminating basal cancer cells or disrupting stromal support. Instead, basal cell loss unleashes an immune-dependent anti-tumor response that requires T cells and NK cells as effector populations.

### Basal cell plasticity enables persistence under therapeutic pressure

Having established that basal cells actively organize an immunosuppressive tumor ecosystem, we next asked how this state behaves under clinically relevant therapeutic pressure. Our lineage-tracing studies showed that basal cells are highly plastic and can generate diverse malignant cell states within PDAC (**Fig. 2**), raising the possibility that therapy may not eliminate basal lineage cells, but instead redirect them into alternative residual states. Because, plastic transitions to drug-tolerant states are increasingly recognized as a central mechanism of adaptive resistance to cancer therapies (*42–44*), we lineage traced basal-derived cells during KRAS(G12D) inhibition, FOLFIRI chemotherapy, or their combination—a promising therapeutic strategy currently being evaluated in clinical trials (*45–47*). To enable synchronized treatment studies and longitudinal monitoring of tumor burden, we established an orthotopic *KPF-basal* reporter allograft model in immunocompromised NSG mice (**Fig. 7A**), in which tumors expressed secreted *Gaussia princeps* luciferase (GLuc), allowing tumor burden to be measured longitudinally via small-volume sampling of the peripheral blood (*48*).

**Figure 7.**
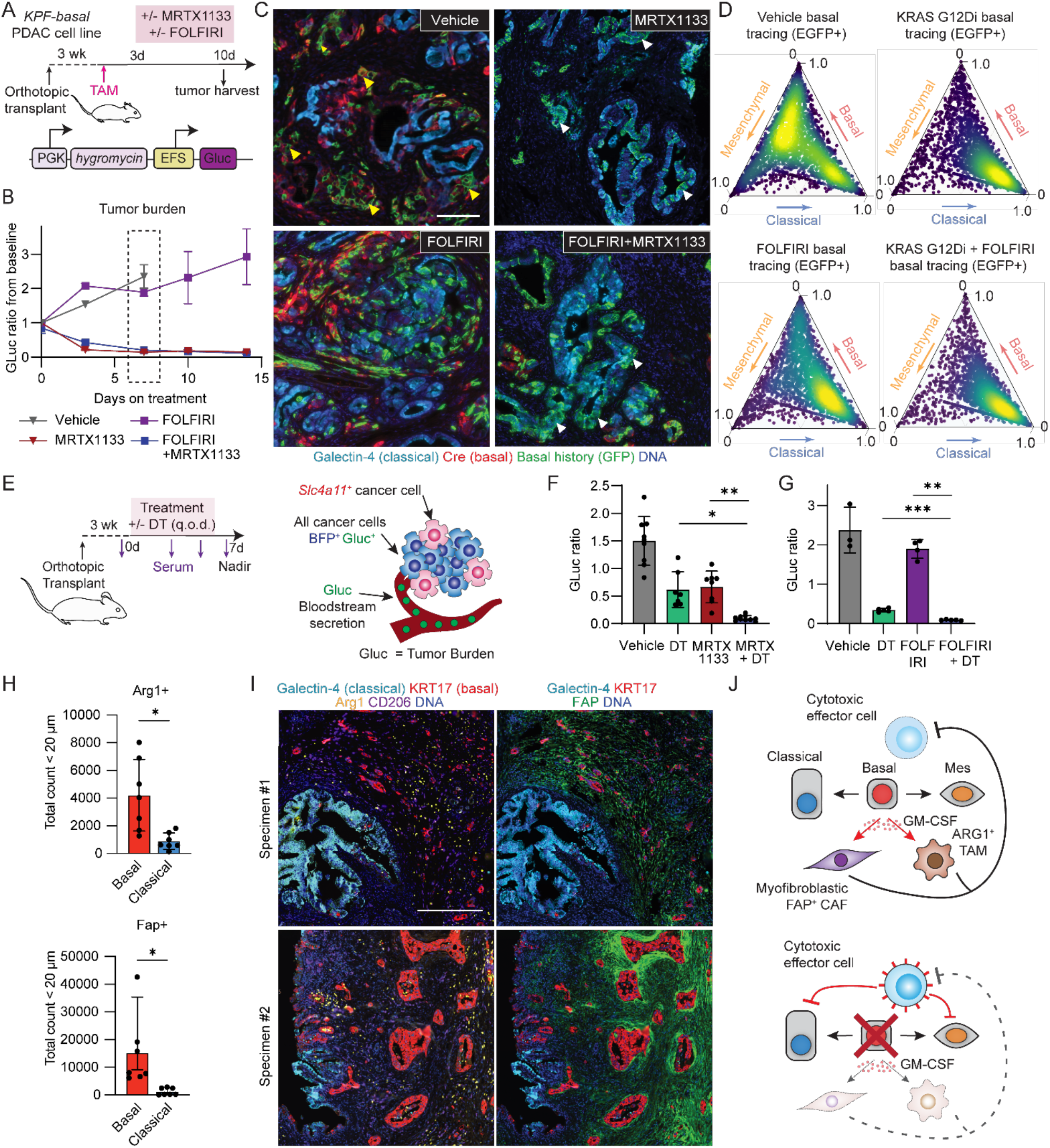
Therapeutic targeting of the basal state and conserved niche organization in PDAC. **(A)** Experimental design for lineage tracing basal-derived cells during therapeutic pressure in orthotopic *KPF-basal* reporter allograft tumors. *KPF-basal* cells were engineered to express secreted GLuc for longitudinal monitoring of tumor burden. After tumor establishment, basal cells were lineage traced, and mice were treated with vehicle, MRTX1133, FOLFIRI, or MRTX1133+FOLFIRI. **(B)** Mean tumor burden in treated orthotopic *KPF-basal* tumors over time, measured as the ratio of circulating GLuc levels to baseline. Error bars, s.e.m.; *n* = 3–6 mice per group. Shaded rectangle indicates tumor harvest timepoint for downstream analysis, including scRNA-seq (day 7). **(C)** Representative immunofluorescence images showing GFP overlap with *Slc4a11*-Cre or Galectin-4 in basal lineage-traced cells in autochthonous GEMM tumors after 7 days of KRAS inhibitor treatment. Note the emergence of GFP⁺ basal-derived cells with classical identity following treatment. Scale bar, 100 μm. **(D)** Predicted cell state probabilities for individual lineage-traced GFP⁺ cells with a basal history after treatment. KRAS inhibition with or without chemotherapy shifted basal-derived cells toward a classical phenotype, compared to controls. **(E)** Left, experimental design for comparing DT-mediated ablation of *Slc4a11-MCD*⁺ basal cells during treatment with MRTX1133, FOLFIRI, and vehicle control in orthotopic tumors. Right, secretion of GLuc into the blood enables longitudinal measurement of tumor burden. **(F)** Mean tumor burden at day 7 of treatment, measured by GLuc in orthotopic KPF-*basal* tumors for KRAS inhibitor or combination therapy. *n* = 7–10 mice per group; error bars, s.d.; **, adjusted *p* = 0.0015; **, adjusted *p* = 0.0052, Welch’s t-test with Holm’s multiple testing correction. **(G)** Mean tumor burden at day 7 of treatment, measured by GLuc in orthotopic KPF-*basal* tumors for chemotherapy or combination therapy. *n* = 3–5 mice per group; error bars, s.d.; **, adjusted *p* = 0.0012, ***, adjusted p = 0.00026. Welch’s t-test with Holm’s multiple testing correction. **(H)** Mean Arg1⁺ or Fap⁺ cell counts within 20 μm of KRT17⁺ basal cancer cells or Galectin-4⁺ classical cancer cells in human PDAC specimens. *n* = 7 patients; error bars, s.d.; *, *p* = 0.0141 (Arg1^+^) and 0.0303 (Fap^+^), Welch’s t-test. **(I)** Representative immunofluorescence images of human PDAC specimens showing basal and classical regions, and their associated stromal populations, within the same tumor. **(J)** Model of basal state-mediated immunosuppression and induction of antitumor immunity

After tumor establishment, we lineage traced basal cells, followed by systemic administration of the allele-specific KRAS(G12D) inhibitor MRTX1133, 5-FU–based chemotherapy (FOLFIRI), or the MRTX1133+FOLFIRI combination (**Fig. 7A)**. The two MRTX1133-containing regimens induced rapid tumor regression, with GLuc decreasing to ∼20% of baseline by day 7, whereas FOLFIRI alone produced only modest disease slowing (**Fig. 7B; Fig. S10A**). We next asked how these therapies affected the fate of basal-lineage cells within residual disease. In untreated tumors, ∼30% of traced basal cells exited basal identity over 7 days. By contrast, MRTX1133 alone or MRTX1133/FOLFIRI drove nearly 90% of traced cells out of the basal state (**Fig. S10B,C**), whereas FOLFIRI produced state transitions similar to untreated controls (**Fig. S10C**).

We and others have shown that basal PDAC cells are highly sensitive to KRAS inhibition (*49, 50*), yet residual tumors under MRTX1133 still contained abundant GFP⁺ basal lineage descendants, indicating persistence of basal-derived cells under KRAS inhibitor pressure (**Fig. 7C)**. Notably, these GFP⁺ cells—and the residual tumor overall—displayed glandular epithelial morphology, strong galectin-4 expression and reduced Slc4a11-Cre expression, consistent with therapy-induced reprogramming of basal cells toward a classical identity under KRAS inhibition (**Fig. 7C; Fig. S10D)**. A similar phenotype was observed with the MRTX1133+FOLFIRI combination: despite the combined, orthogonal selective pressures of targeted KRAS inhibition and cytotoxic chemotherapy on basal and classical states, respectively, residual basal lineage descendants persisted and predominantly adopted classical features (**Fig. 7C)**. ScRNA-seq of lineage-traced cells confirmed these therapy-induced state transitions at the transcriptional level (**Fig. 7D**). Thus, KRAS inhibition does not fully eliminate basal cells; instead, basal-lineage cells persist by adopting classical features as a mechanism of escape, even when KRAS inhibition is combined with chemotherapy.

### Targeting the basal state enhances therapeutic response

Basal lineage tracing showed that KRAS inhibition can drive basal-derived cells into classical residual states rather than eliminating them. We next asked whether the basal cell state nevertheless remains functionally important during therapy. To test this, we treated GLuc-expressing orthotopic *KPF-basal* tumors with MRTX1133 or FOLFIRI, with or without DT-mediated basal cell ablation (**Fig. 7E**). Although MRTX1133 and FOLFIRI each reduced tumor burden, both left substantial residual disease. In contrast, addition of basal state ablation produced markedly deeper tumor regressions (**Fig. 7F,G; Fig. S10E,F**). Thus, basal state elimination can further reduce therapy-persistent disease, supporting the idea that basal state programs and the niches they maintain remain important barriers to complete therapeutic response.

### Basal-state associated TME niches are conserved in human PDAC

Finally, we asked whether the basal-associated stromal architecture defined in our mouse models is conserved in human PDAC. We profiled seven primary PDAC resection specimens without prior treatment, selected to contain both basal and classical cancer cell states, allowing local comparison of stromal organization around distinct malignant states within the same human tumors. At the level of local TME architecture, ARG1⁺ myeloid cells and FAP⁺ fibroblasts were significantly enriched near KRT17⁺ basal cancer cells compared with galectin-4⁺ classical cancer cells (**Fig. 7H,I**). Thus, the basal state-associated immunosuppressive niche identified functionally in mouse tumors is also evident as a conserved spatial architecture in human PDAC.

Together, these findings support a model in which basal cancer cells act as plastic, niche-organizing cells that sustain immunosuppressive myeloid and fibroblast populations and block cytotoxic lymphocyte access (**Fig. 7J**). Loss of basal state function dismantles this suppressive circuit, remodels residual classical niches, and enables T/NK cell–mediated tumor killing. This conserved association of basal tumor regions with suppressive stromal compartments in human PDAC motivates therapeutic strategies aimed at disrupting the basal state and the associated ecosystem.

## DISCUSSION

Intra-tumoral phenotypic heterogeneity of cancer cells is a defining feature of PDAC, but whether distinct malignant cell states encode specialized functions in tumors has remained unresolved. Using genetic systems for lineage tracing, selective ablation, and cell state-restricted gene perturbation *in vivo*, together with spatial computational approaches that reconstruct multicellular niche organization and remodeling trajectories, we directly interrogated the basal and classical cell states and uncovered a striking functional asymmetry: ablation of the basal cell state, but not the classical cell state, triggered rapid and durable tumor collapse. These analyses identify the basal state as a driver of tumor maintenance through two linked properties: phenotypic plasticity and the enforcement of a distinct multicellular niche enriched for immunosuppressive macrophages and fibroblast programs, which is dismantled upon basal state loss.

It is notable that the induction of plasticity in a subset of epithelial cells, repair-associated immunosuppressive macrophages, and contractile myofibroblasts are cardinal features of wound healing across different epithelia (*51–53*). Interestingly, our analyses of large-scale scRNA-seq datasets of human patient tumors showed that a highly plastic malignant state that closely resembles the basal state in PDAC is present in multiple common and lethal carcinomas (*19*), and correlates strongly with the *Stress* archetype state—a cell state that is conserved across most human cancers independently of lineage (*1, 2, 19*)—suggesting the function of such regenerative plastic states in driving stromal remodeling and immunosuppression may be conserved across many human cancers. An important direction for future work will be to functionally define whether these recurrent regenerative plastic/basal-like programs similarly organize tumor-promoting malignant-stromal interactions across cancer types, as suggested by several recent studies (*54–57*).

In the pancreas, injury induces epigenetic and transcriptional states linked to neoplastic transformation (*58, 59*), and epithelial plasticity is a hallmark of normal wound repair, where it is typically transient and resolves once tissue integrity is restored (*60, 61*). Our results indicate that the basal cancer cells co-opt and stabilize this regenerative state—converting a transient repair response into a persistent cancer cell state—that continuously recruits and sustains immunosuppressive fibroblast and myeloid populations, thereby promoting tumor growth. Previous work has described association of basal PDAC cells in myofibroblastic and myeloid-rich niches across the disease continuum (*9, 15, 62*), including in metastases (*5*). These studies primarily focused on how stromal populations induce or reinforce basal or related wound-like malignant states, which include tumor necrosis factor-a (TNFa) produced by the basal-associated macrophages (*62*) and myofibroblastic CAFs that activate EGFR signaling in basal-like PDAC cells (*57*). In our study, we address the reciprocal direction: whether basal malignant cells actively remodel and maintain the stromal and immune niche.

By combining spatial association with state-restricted perturbation, our study moves this model from correlation to mechanism. Basal PDAC cells were consistently associated with ARG1^+^ macrophages and myCAFs, a spatial relationship that also emerges during formation of the precursor to the basal state in recent work (*16, 27*). Here we tested whether basal cells actively maintain this niche in established tumors using genetic systems we developed for cell state-restricted and temporally controlled gene perturbation *in situ*. Using this system, we identify the basal-derived cytokine GM-CSF/*Csf2*—an established driver of PDAC immunosuppression(*36, 37*)—as a key mediator of myeloid remodeling, positioning the basal state and its secreted programs upstream of a suppressive myeloid circuit in PDAC. Notably, secreted-factor expression in basal cells appears organized as a broader co-regulated module that includes myCAF-associated factors such as *Pdgfb* and *Tgfb* (*38, 39*), raising the possibility that multiple basal-derived signals act in parallel on distinct stromal compartments.

The conservation of archetypal malignant programs across cancers suggests that tumor evolution preserves a core repertoire of cell states rather than converging on a single optimal phenotype (*1, 2*). One explanation is division of labor, in which distinct malignant states perform complementary functions that benefit the tumor as a whole. In PDAC, classical cells may support proliferation, basal cells facilitate plasticity and immune suppression, and mesenchymal cancer cells may mimic fibroblast function, providing support to epithelial cancer cells. However, our cytoablation results indicate that classical cells are dispensable for short-term maintenance of fully formed, aggressive PDAC tumors. We and others have shown that classical cells resist oncogenic KRAS inhibition and can serve as a reservoir for relapse (*63, 64*), but whether they have other functions in PDAC tumors remains an important area of future inquiry. A prior study found that suicide gene-mediated ablation of vimentin^+^ PDAC cells robustly suppressed tumor growth and extended survival (*65*), implicating EMT-like programs in PDAC maintenance. However, vimentin is broadly expressed across non-classical PDAC states, including most basal PDAC cells (**Fig. 1B**), and therefore vimentin-directed ablation cannot be interpreted as a mesenchymal-specific perturbation, which confounds the interpretation of this data. Future studies employing faithful reporters are needed to delineate the function of the mesenchymal cells in PDAC evolution, maintenance, and stromal interactions.

Our findings have several therapeutic implications. First, they identify the basal state as a source of phenotypic adaptation under therapy. Although basal cells are highly sensitive to KRAS inhibition, lineage tracing shows that basal-derived cells can persist by transdifferentiating toward a classical phenotype, thereby contributing to residual disease. This supports a model in which acute resistance arises not only through selection of pre-existing resistant states, but also through therapy-induced state transitions of plastic cells. Second, combining basal state cytoablation with KRAS inhibition or chemotherapy yields deeper responses, likely by simultaneously limiting plasticity and stromal remodeling.

Our findings further raise the possibility that tumor microenvironment remodeling in response to KRAS inhibition is mediated, at least in part, through selective depletion of the basal state and the associated TME-modifying cytokine module rather than direct effects on KRAS signaling and cell survival alone. Consistent with this, recent studies of allele-selective and multi-RAS inhibitors in immunocompetent PDAC models demonstrate increased CD8⁺ T cell infiltration, reduced myeloid-mediated immunosuppression, CAF reprogramming, and synergy with checkpoint blockade (*66–69*), paralleling the effects observed following basal ablation. Our findings suggest a highly malleable model where plasticity of basal cells permits evasion of KRAS inhibition rather than resulting in selective elimination of the basal cell state. In adapting to classical identity, tumors under KRAS inhibition have a lower ability to promote immunosuppression, allowing for lymphocyte infiltration and increased efficacy of checkpoint inhibitors. However, even transient basal state ablation produced a stronger therapeutic response than KRAS inhibition alone, suggesting that RAS inhibition may incompletely suppress the basal program. Identifying additional basal state dependencies—particularly those controlling plasticity, immunosuppression, or residual niche maintenance—may therefore provide a strategy to deepen responses and limit adaptive escape.

Several limitations should be considered. First, our data do not establish that the classical state lacks function in PDAC. Rather, in short-term ablation experiments in established tumors, classical cell loss did not produce a comparable growth defect or the same pattern of tumor microenvironment remodeling observed after basal cell ablation. Classical cells may have important functions in other settings, including tumor initiation, immune evasion, or relapse after therapy. Second, our reporters and perturbation systems focus on major basal and classical programs and do not fully address the roles of intermediate, transitional, or rarer malignant populations. Third, although we identify basal-derived *Csf2* as a functional mediator of myeloid remodeling, it is likely that other basal-derived cytokines or secreted factors contribute independently or cooperatively to stromal remodeling, immune suppression, or tumor maintenance. Future studies will be required to systematically dissect how distinct components of this broader secreted factor module contribute to stromal architecture, immune exclusion, and malignant cell persistence.

In conclusion, our findings identify the basal cell state as a central driver of cell state heterogeneity and immunosuppression in PDAC. Much like the conductor of an orchestra, the basal state coopts regenerative plasticity and paracrine signaling interactions to organize a persistent wound healing-like response, which drives cancer progression and therapeutic resistance. Targeting the basal state ecosystem may therefore provide a strategy to dismantle immune suppression and deepen therapeutic responses in one of the most aggressive and treatment-refractory human cancers.

## ACKNOWLEDGEMENTS

We thank the members of the Tammela and Pe’er laboratories for discussions; the members of the Sloan Kettering Institute’s Single Cell Analytics and Innovation Lab (SAIL) computational unit and R. Chaligne; W. Kang, M. Tipping and the members of the Molecular Cytology Core for histology support; A. Santella and E. Rosiek for help with image analysis and quantification; E. de Stanchina and the Antitumor Assessment Core Facility for support with drug administration and tumor transplant experiments; R. Gardner, M. Kweens and A. Longhini for FACS support; N. Cruz for laboratory management; J. Christensen and J. Hallin for providing MRTX1133. We also thank J. Reyes, J. Chan, C. H. Pan for sharing experimental protocols and guidance.

## MATERIALS AND METHODS

### Animal studies

All animal studies were approved by the Memorial Sloan Kettering Cancer Center (MSKCC) Institutional Animal Care and Use Committee (protocol, #17-11-008). MSKCC guidelines for the proper and humane use of animals in biomedical research were followed. All mice were housed and monitored in a 12-hour light/12-hour dark cycle at 20-25°C and 30-70% humidity with food and water provided by the listed investigators and veterinary staff under the Research Animal Resource Center at MSKCC.

All genetically engineered mice were maintained in C57/BL6 and Sv129 mixed backgrounds. Autochthonous *Kras^frt-stop-frt(FSF)-G12D/+^; Trp53^frt/frt^; Pdx1-FlpO* (*KPF)* mice were crossed to new and previously published reporter alleles. For basal cell lineage tracing experiments, *KPF* mice carrying either *Rosa26^lox-stop-lox(LSL)-mTmG/+^* or *Hipp11^FSF-BG/+^* were crossed with the previously-generated reporter *Slc4a11^FSF-MCD/+^* containing a mScarlet fluorescent reporter, CreER, and ablation cassette DTR (*19*). For classical cell lineage tracing experiments, *KPF* mice carrying either *Rosa26^LSL-mTmG/+^* or *Hipp11^FSF-BG/+^* were crossed with the classical state reporter *Lgals4^FSF-MACD/+,^* which contains an mScarlet fluorescent reporter, AkaLuc bioluminescent reporter, CreER, and DTR ablation cassette.

Expression of “*MCD*” or “*MACD*” is restricted to the cancer cells through excision of the frt-stop-frt (FSF) cassette by *Pdx1-*FlpO, which concomitantly induces expression of KRAS(G12D) and loss of p53 (**Fig. 1A**). Normal control mice used in experiments did not harbor the pancreas-specific Flp recombinase allele, and therefore did not form autochthonous tumors. Mouse genotyping was performed at two weeks of age. Tumors were harvested at planned endpoints or until they reached a humane endpoint (weight loss, distended abdomen, and hunching). Mice were euthanized by CO_2_ asphyxiation.

*NOD.Cg-Prkdc^scid^;Il2rg^tm1Wjl^/SzJ* (NSG; The Jackson Laboratory, 005557) or B6129SF1/J F1 hybrid mice (F1) mice (The Jackson Laboratory, 101043) were used as recipients in orthotopic studies.

### Generation of reporter models

For the generation of the Hipp11-FSF-BG donor vector, ∼5,000-bp 5′ and ∼7,000-bp 3′ homology arms targeting the *Hipp11* safe-harbor intergenic region, located between the *Eif4enif1* and *Drg1* genes, were amplified from genomic DNA of C57BL/6 mouse embryonic stem cells using high-fidelity PCR (KME-101, Toyobo). A homology-directed repair template donor vector was constructed by flanking the CAG-LoxP-Frt-Neomycin-PGKpA-SV40pA-Stop-Frt-TagBFP-3xFlag-bGlobinpA-LoxP-mEGFP-WPRE-gGHpA (BG) cassette with the 5′ and 3′ homology arms, followed by cloning into the pUC19 plasmid backbone using Gibson assembly (#638949, Takara).

For the generation of the *Lgals4-MACD* donor vector, ∼1,500-bp 5′ and 3′ homology arms flanking the end of *Lgals4* exon 10 were amplified from C57BL/6 genomic DNA using high-fidelity PCR (KME-101, Toyobo). A homology-directed repair template donor vector was constructed by flanking the frt-PGK-Hygro-pA-frt-T2A-mScarlet-AkaLuc-T2A-CreER-P2A-DTR (MACD) cassette with the 5′ and 3′ homology arms and cloned into the pUC19 plasmid backbone using Gibson assembly (#638949, Takara).

### Embryonic stem cell targeting, genotyping and chimera generation

A *Kras^FSF-G12D/+^;Trp53^frt/frt^* (*KPfrt*) mouse embryonic stem cell (mESC) line in the C57BL/6J background was generated by crossing a hormone-primed C57BL/6J *Trp53^frt/frt^* female mouse with a *Kras^FSF-G12D/+^ ; Trp53^frt/frt^* male mouse. At 3.5 days after coitum, blastocysts were flushed out from the pregnant uterus, isolated and cultured on a mouse embryonic fibroblast (MEF) feeder layer. Individual ES cell lines were genotyped by PCR detection of *Kras^FSF-G12D/+^, Trp53^frt/frt^* alleles and Y chromosome (*Zfy*).

For the generation of *Hipp11^FSF-BG^*^/+^ knock-in mESCs, the donor vector (*Hipp11-FSF-BG*) was targeted to the ubiquitously active *Hipp11* locus (*70*) in *KPfrt* mESCs, which harbor Flp recombinase-activatable *Kras^FSF-G12D^; Trp53* loss-of-function (*Trp53^frt/frt^*) alleles (**Fig. S1A-C**) (*18*). The donor vector and ribonucleoprotein (RNP) complex containing HiFi Cas9 nuclease V3 (IDT, 1081061) and a crRNA–tracrRNA duplex (IDT) were co-transfected into *KPfrt* mESCs by electroporation using the Lonza 4D-Nucleofector system.

*KPfrt* mES cells were thawed 2 days before targeting, and the media were changed 1 day and 2 hr before electroporation. Before electroporation, sequence-specific crRNA and universal tracrRNA were resuspended in IDTE buffer (IDT) at a concentration of 200 µM and the crRNA–tracrRNA duplex was then formed (final concentration, 44 µM) by combining an equimolar concentration of crRNA and tracrRNA and annealing at 95 °C for 5 min (followed by cooling down to room temperature at ramp rate of 0.1 °C s^−1^). RNP complexes were formed by combining 22 pmol of crRNA–tracrRNA duplex and 22 pmol HiFi Cas9 nuclease and incubating at room temperature for 20 min. For each electroporation, 500,000 mESCs, 1 µl donor vector (3 µg µl^−1^), 1 µl RNP complex, 2 µl electroporation enhancer (10 µM, IDT), 16.4 µl Nucleofector P3 primary cell solution and 3.6 µl Nucleofector Supplement 1 were combined and loaded into electroporation cuvette. Following electroporation, mESCs were plated onto feeder MEFs. After 48 h, cells were selected with neomycin for 1 week. Resistant clones were manually picked, expanded, and validated by genotyping and sequencing using primers specific for DTR, mScarlet, and the targeted homology arm junction. Primers are listed in **Table S4**.

Chimeric F_0_ mice were obtained by injecting genotype-verified mESCs into host embryos at the eight-cell stage. F0 mice were genotyped at 2 weeks of age using the primers listed in **Table S4**. F_0_ mice were crossed into the *Slc4a11^FSF-MCD/+^* or *Lgals4^FSF-MACD/+^*; *Pdx1-FlpO* background to generate lineage tracing reporter mice appropriate for the listed experiments.

### Direct embryo targeting (for Lgals4-MACD)

For generation of the *Lgals4^FSF-MACD/+^* mouse model, female donor mice in the *KPfrt* background were superovulated, and one-cell embryos were harvested. Zygotes were injected into the male pronucleus with a Cas9/crRNA ribonucleoprotein (RNP) complex targeting the *Lgals4* locus, together with the donor construct containing the FSF-MACD cassette. Following injection, embryos were cultured to the two-cell stage and transferred into pseudopregnant recipient females, which carried the embryos to term.

Founder pups were screened by PCR for insertion of the MACD cassette downstream of exon 10 of *Lgals4* using primers listed in **Table S4**. Founders validated by PCR were then further validated by copy number assay and genotyping. Validated founders were bred with *KPfrt* mice to confirm germline transmission and establish the colony for subsequent validation studies.

### Cell line generation

Cell lines were generated by isolating tumor cells, as described in the methods section describing isolation of primary pancreas tumor cells below, from autochthonous reporter mice from the following backgrounds and used with *in vitro* assays and orthotopic transplantation models.

- *Kras^G12D/+^ Trp53^Δ/ Δ^ Rosa26^LSL-mTmG/+^Slc4a11^MCD/+^ Pdx1-FlpO*
- *Kras^G12D/+^ Trp53 ^Δ/ Δ^ Hipp11^BG/+^Slc4a11^MCD/+^ Pdx1-FlpO*
- *Kras^G12D/+^ Trp53 ^Δ/ Δ^ Hipp1^-BG/+^Lgals4^MACD/+^ Pdx1-FlpO*

Additionally, a lentivirus construct (pPHGmiR_PGK-Hygro-EFS-GLuc_pLT) was introduced to cells lines to allow for non-invasive tumor measurements during orthotopic experiments via submandibular blood collection. After transfection of cells with this lentivirus construct, cells were selected for using hygromycin. Cell lines were cultured in 2D in RPMI (Gibco) supplemented with 1% GlutaMAX (#35050061, Thermo Fisher Scientific), 1% Pen/Strep (#15070063, Thermo Fisher Scientific), and 10% heat-inactivated FBS (#SH30910.03, HyClone). All mouse cell lines were tested for Mycoplasma every 2 months using a PCR assay and confirmed to be negative.

### Design of GM-CSF overexpression vector

Plasmids expressing wild-type (WT) murine cDNAs encoding GM-CSF (*Csf2*), were generated. All constructs were cloned into a lentiviral backbone of the form pLV-rev(PGK>Blast):loxP:STOP:loxP:P2A:GFP:P2A:cDNA.

### Generation of PGK-Blast-LoxP-frameshift-LoxP-P2A-Cas9 (inducible Cas9)

A lentiviral construct encoding PGK-Blast-loxP-frameshift-loxP-P2A-Cas9 was designed and cloned into a pLenti backbone. The construct was designed with the following features: a Blasticidin resistance gene lacking a stop codon at its 3′ end, a stop codon immediately downstream of the first loxP site to prevent read-through, a ∼100 bp frameshift sequence flanked by loxP sites that is in-frame following Cre-mediated LoxP excision, and a nuclear localization sequence (NLS) appended to the C-terminus of Cas9, which carries a stop codon at its 3′ end.

To assemble the full-length construct, the entire insert was divided into five fragments, each designed as a geneblock (Integrated DNA Technologies). Gene-specific oligonucleotides containing Gibson assembly overhangs were synthesized and used as primers to amplify each geneblock by high-fidelity PCR (Q5 polymerase, NEB), generating five PCR fragments with compatible overlapping ends. To facilitate efficient final assembly, fragments were cloned in two intermediate steps: three fragments were first assembled into an intermediate vector, and the remaining two fragments were assembled into a separate intermediate vector. The verified intermediate constructs were then digested to release the respective inserts, and all five fragments were assembled simultaneously into the linearized pLenti backbone by Gibson assembly. The final construct was verified by Sanger sequencing.

### Generation of HAMMER sgRNA library

Mouse HAMMER sgRNA libraries were constructed using the ALPA (Array-based Lentiviral Production of sgRNA Arrays) cloning strategy as previously described (*40*), with the identical backbone plasmid (pYJA5). Briefly, twenty-nucleotide sgRNA sequences targeting mouse protein-coding genes were incorporated into oligonucleotides synthesized (Bio Basic Inc.). Three PCR fragments (C1, M, and C2) were amplified using high-fidelity PCR (Q5 polymerase, NEB), where each fragment was amplified using sgRNA-containing oligonucleotides as primers, thereby embedding up to four sgRNA sequences per construct. The three fragments were then assembled into BbsI-linearized pYJA5 vector by Gibson assembly. The assembled constructs were transformed into competent cells for library propagation. For detailed cloning procedures, refer to the original ALPA cloning publication (*40*).

### Generation of lentivirus

HEK293T cells were transfected with lentiviral transfer plasmids and the second-generation lentiviral packaging plasmid psPAX2 (Addgene, 12260) and the envelope plasmid pMD2.G (Addgene, 12259) using either the TransIT-LT1 kit (Mirus Bio, MIR 6000) or PEI (MedChem Express, HY-K2014). At 24 hr after transfection, the medium was discarded and replaced with fresh complete medium. Viral medium was collected and filtered through 0.45-µm PES filters (Cytiva, 6780-2504) at 48 hr and 72 hr after transfection. All viral media collected were concentrated using an ultracentrifuge with rotor speed set at >130,000g for 2 hr at 4 °C. The supernatant was discarded into bleach and viral pellets were allowed to solubilize overnight at 4 °C. Concentrated virus was gently mixed and aliquoted. The aliquots were immediately placed on dry ice and stored at −80 °C.

### Orthotopic models

Mouse orthotopic pancreas transplants were performed in NSG or B6129SF1/J mice (JAX stock #101043). Orthotopic tumors were generated by implanting 50,000 cells from a 2D culture into the tail of the pancreas. Mouse PDAC cell lines, containing *Slc4a11-MCD* or *Lgals4-MACD in the KPF background with either lineage tracing construct (Hipp11-BG or Rosa26-mTmG),* were thawed and passaged for 1 to 2 months prior to orthotopic transplantation. Animal studies began approximately 3 weeks after tumor implantation. All animals were monitored weekly, and tumors were harvested at a maximum size of 1 cm^3^ or if the animals reached a humane endpoint for euthanasia.

### In vivo lineage tracing

Tumor bearing mice were treated orally with one dose of tamoxifen (200 mg/kg via oral gavage) at approximately 6 weeks after birth or 3 weeks after transplantation and monitored daily until end point (3, 7, or 14 days). Tamoxifen was dissolved in corn oil at 20 mg/mL for one hour at 60°C.

### In vivo genetic perturbation experiments

In genetic perturbation experiments, murine *KPF*; *Slc4a11-MCD* and *KPF*; *Hipp11-BG* cell lines were transduced with lentiviruses containing loxP-frameshift-loxP-Cas9 and sgRNAs and selected for antibiotic resistance. These cell lines were transplanted orthotopically into B6129SF1/J F1 generation mice or NSG mice, as described above, and tumors were allowed to develop. Mice were administered 4 doses of 100 mg/kg tamoxifen resuspended in corn oil by oral gavage for 10 days until the endpoint.

In genetic overexpression experiments, murine *KPF*; *Slc4a11-MCD* and *KPF*; *Hipp11-BG* cell lines were transduced with lentiviruses containing LSL-cDNAs and Cre recombinase and selected for antibiotic resistance. These cell lines were transplanted orthotopically into B6129SF1/J F1 generation mice and tumors were allowed to develop.

### Therapeutic administration (KRAS inhibitors, chemotherapy, diphtheria toxin)

*KPF* mice bearing autochthonous PDAC tumors or NSG mice bearing orthotopic *KPF* cell line allografts were intraperitoneally administered freshly prepared MRTX1133 in Captisol (#HY-17031, MedChemExpress) at 30 mg/kg twice a day starting at 7 weeks after birth or 3 weeks after transplantation of cell lines, or in both cases when tumors were palpable. Mouse body condition was monitored daily, and no significant cytotoxic side effects were observed during treatment. In genetic ablation experiments, mice with established tumors were treated with DT (#D0564, Sigma) resuspended in sterile PBS at 50 µg/kg by intraperitoneal injection every other day starting at the beginning of treatment. For autochthonous *KPF* experiments, treatment was initiated in mice with palpable tumors at approximately seven weeks after birth. For orthotopic transplantation experiments, treatment was initiated in NSG or B6129SF1/J F1-generation mice approximately three weeks after transplantation of murine PDAC cell lines, once tumors were established. In chemotherapy experiments, NSG mice bearing *KPF* cell line transplants were treated FOLFIRI experiments intraperitoneally three times weekly for the duration of treatment, which consists of: 5-fluorouracil (30 mg/kg), irinotecan (73 mg/kg), and leucovorin (90 mg/kg). Vehicle controls were used for all drug treatments.

### Administration of blocking antibodies

For T and NK cell depletion, CD8 (clone 2.43, BioXcell #BE0061), CD4 (clone GK1.5, BioXcell #BP0003), and NK1.1 (clone PK136, BioXcell #BE0036) neutralizing antibody or rat IgG2b isotype control antibody (clone LTF-2 BioXcell #BP0090) were administered at 200 µg per antibody per mouse every third day starting two weeks prior to DT treatment for 8 total doses.

### In vivo bioluminescence imaging, plasma sampling, and luciferase assays

Recipient mice with AkaLuc-expressing tumors were dosed with 100µl of 30mM of AkaLumine-HCl substrate resuspended in PBS (TokeOni, 808350) or Firefly Luciferase expressing tumors were dosed with 100 µl of 15.7 mM of D-luciferin potassium salt resuspended in PBS (Gold Biotechnology, LUCK) and imaged on an IVIS Lumina II (Perkin Elmer). Signals were recorded for 20 5-second exposures to capture peak luminescence. Luciferase signals were acquired through analysis on the Living Image software (Revvity). The total flux (p/s) for each mouse was calculated for the 20 exposures, and the peak value was used for each timepoint in individual animals. A fold-change over time was generated by dividing the calculated value at each time point from the pre-treatment value.

For mice containing the Gaussia luciferase reporter, whole venous blood was harvested by puncturing the submandibular vein and collecting 40µl in blood collection vials (#02–675-185, Fisher Scientific). Plasma was separated by centrifugation at >8000 g for 10 minutes at 4°C and diluted 1:10 in PBS. Plasma was treated with 50 µl of the Gaussia luciferase substrate Coelenterazine-h (#301, NanoLight, diluted 1:10 in PBS) at 200uM and luminescence was immediately measured with the BioTek Cytation 5 (Agilent) at a working concentration of 20 µM. When calculating the tumor burden from Gaussia luciferase measurements, two technical replicates from the serum measurement were averaged and background (control well with PBS only) was subtracted. A fold-change from the pre-treatment value was calculated at each time point for each individual mouse within the treatment groups.

### MRI Longitudinal in vivo imaging

Mice were anesthetized with 1-2% isoflurane gas in air and scanned on a 9.4-Tesla Bruker Biospec scanner (Bruker Biospin Corp., Billerica, MA) with a 12-cm gradient coil (maximum gradient strength 530 mT/m). An ID 40 mm Bruker volume coil was used for RF excitation and detection. Mouse respiration during the scan was monitored by a physiological monitoring unit (SA Instruments, Stony Brook, NY). Mice were positioned prone in the scanner. First T1-weighted scout images along 3 orthogonal orientations were acquired. T2-weighted mouse axial abdominal images were then acquired using RARE(Rapid Acquisition with Relaxation Enhancement) fast spin-echo sequences with the following acquisition parameters: slice thickness of 0.5 mm, repetition time 2.35 s, echo time, 26 ms, FOV 30 x 25 mm and 256×256 matrix.

MRI images were analyzed using 3D Slicer software (*71*). ROI was manually annotated for each FOV, and volumes (mm^3^) were calculated with 3D Slicer at each time point for individual mouse scans. Pancreas and tumor volumes were recorded and averaged for each time point.

### Isolation of primary pancreatic cancer cells for scRNA-seq

Orthotopically transplanted and primary tumors were dissociated into single-cell suspensions, and cancer cells were isolated for scRNA-seq by FACS. To dissociate tumors into single-cell suspensions, tumors were finely chopped with scissors and incubated in a digestion buffer of Collagenase IV (17104019, ThermoFisher Scientific, 0.1 U/ml), Dispase (#354235, Corning, 0.6 U/ml), DNase I (#69182–3; Sigma Aldrich, 10 U/ml), and Soybean Trypsin Inhibitor (#T9003, Sigma Aldrich, 0.1 mg/ml) dissolved in HBSS with Mg^2+^ and Ca^2+^ (#14025076, Thermo Fisher Scientific) in gentleMACS Tubes (Miltenyi Biotec) for 42 minutes at 37°C using the gentleMACS Octo Dissociator (#130–096-427, Miltenyi Biotec). After mechanical and enzymatic dissociation, tumor cells were washed with HBSS + Mg^2+^ and Ca^2+^ and filtered through a 100 µm cell strainer (#352360, Falcon) and spun at 300 *g* for 5 minutes at room temperature. Cells were then washed with S-MEM media with 2% of heat inactivated FBS and pelleted at 300 *g* for 5 minutes at 4°C. The supernatant was removed, and the pellet resuspended in Fluorescence-Activated Cell Sorting (FACS) buffer (200 mM EDTA with 2% of heat-inactivated FBS in PBS) before being passed through a 40 µm strainer (#352340, Falcon).

For mouse cell suspensions to prepare for FACS, cell suspensions were blocked at 4°C for 5 minutes with rat anti-mouse CD16/CD32 (Mouse BD Fc Block, #553142, BD Biosciences) in FACS buffer and incubated for 20 minutes with a mix of 4 fluorophore-conjugated antibodies binding CD45 (#25-0451-82, Invitrogen, 1:160), CD31 (#102418, Biolegend, 1:40), CD11b (#101216, Biolegend, 1:160), and TER-119 (#116222, Biolegend, 1:80). For phenotypic profiling of cancer cells within mouse tumors, fluorophore-conjugated antibodies binding EpCAM (CD326, #118233, Biolegend, 1:50) were used. Cells were then stained with Fixable Viability Stain 780 (FVS780; #565388, BD Biosciences) as a live/dead marker. Cells were subsequently fixed according to the 10x Genomics fixation protocol.

### Tissue histology and immunofluorescence

For cryosection immunofluorescence, tumors harvested at experimental endpoints were fixed in 10% neutral buffered formalin (Richard-Allan Scientific), embedded in OCT (#23-730-571, Fisher Scientific), and cut into 10 µm sections. Sections were fixed in acetone for 10 minutes, washed in PBS, and blocked for 30 minutes in PBS with 0.1% Triton-X, 0.2% BSA, and 5% donkey serum (#D9663, Sigma-Aldrich). The slides were incubated with primary antibodies (Supplementary Table 2) overnight at 4°C in blocking buffer. The slides were incubated with secondary antibodies (Thermo Fisher Scientific, 1:500) for 2 hours at room temperature.

For manual paraffin immunofluorescence or immunohistochemistry, tumors harvested at experimental endpoints were fixed in 10% neutral buffered formalin (Richard-Allan Scientific), embedded in paraffin, and cut into 5 µm sections. The slides were heated for 1 hour at 60°C, deparaffinized, rehydrated with an alcohol series and incubated for 20 minutes in a pressure cooker on 95°C in Tris-EDTA antigen retrieval buffer (#E1161, Sigma-Aldrich). For immunofluorescence, the tissues were subsequently blocked in PBS with 0.1% Triton X-100, 2% BSA, and 5% donkey serum (#D9663, Sigma-Aldrich). The slides were incubated with primary antibodies overnight at 4°C in blocking buffer (**Table S4**). Slides were incubated for 2 hours at room temperature in secondary antibodies (Thermo Fisher Scientific, 1:500).

For immunohistochemistry, tissues were blocked in BLOXALL Endogenous Peroxidase and Alkaline Phosphatase Blocking Solution (SP-6000, Vector laboratories) for 10 minutes and incubated in 2.5% Normal Horse Serum Blocking Solution (S-2012, Vector laboratories) for 1 hour before incubating in primary antibodies overnight at 4C. Slides were incubated for 1 hour at room temperature in HRP-conjugated secondary antibodies (MP-7401-50, Vector Laboratories). Color was developed using ImmPACT (TM) DAB HRP Substrate (SK-4105, Vector laboratories). Slides were counterstained using with Richard-Allan Scientific; Gill; 1, 2, 3 Hematoxylin (#72404, Thermo fisher Scientific) for 3 minutes and Richard-Allan Scientific; Signature Series Bluing Reagent (#7301, Fisher Scientific) for 10 seconds and mounted on cover slips with Permount™ Mounting Medium, Liquid (#SP15100, Fisher Scientific).

For multiplexed immunofluorescence (Leica Bond RX), samples were pretreated with EDTA-based epitope retrieval ER2 solution (Leica, AR9640) for 20 minutes at 95°C. 4-plex antibody staining and detection were conducted sequentially. Antibodies used for multiplexed IF are listed (**Table S4**). The primary antibodies were incubated for 1h at RT. For rabbit antibodies, Leica Bond Polymer anti-rabbit HRP (included in Polymer Refine Detection Kit (Leica, DS9800) was used; for the goat antibodies, a rabbit anti-goat (Jackson ImmunoResearch303-007-003) secondary antibody was used as linkers for 8 min before the application of the Leica Bond Polymer anti-rabbit HRP for 8 min at RT. After that, Alexa Fluor tyramide signal amplification reagents (Life Technologies, B40953, B40958) or CF® dye tyramide conjugates (Biotium, 92172, 96053, 92174) was used for detection. After each round of IF staining, epitope retrieval was performed for denaturation of primary and secondary antibodies before another primary antibody was applied. After 1 hr incubation, Leica Bond Polymer anti-rabbit HRP was applied followed by Alexa Fluor tyramide conjugate 488, 543, 594, and 647 (Life Technologies, B40953, B40958), or CF® dye tyramide conjugate 430 for signal amplification. At each round, epitope retrieval was performed for denaturation of primary and secondary antibodies before the following primary antibody was applied.

All tissues were counterstained with DAPI (#D9542, Sigma Aldrich, 5 µg/ml) for 5 minutes and mounted on cover slips with Mowiol mounting reagent (#475904, Millipore Sigma). Slides were scanned on a Panoramic Scanner (3DHistech) with a 20X/0.8NA or 40X objective.

### Human tissue samples and COMET multiplex immunofluorescence

5µm tissue section was trimmed from a FFPE block and placed at the center of a clean glass slide. The slide was air dried and baked at 42°C for 3 hours and stored in a desiccator. Epredia PT Module was used to deparaffinize and retrieve epitopes (Epredia Dewax and HIER Buffer L; Epredia Dewax and HIER Buffer H). Slide was then washed twice with 1X Multistaining buffer (BU06) and loaded on to the COMET. Appropriate volumes of primary antibodies, secondary antibodies, 5µg/ml DAPI (D3571, Thermofisher), Multistaining buffer, Quenching buffer (BU08-L), Imaging buffer (BU09), and Elution buffer (BU07-L) were freshly made and loaded into the fluidics compartment of the instrument. 12 mm x 12 mm FOV was captured in a tiled fashion only where the tissue was auto detected. The primary antibodies were used at the following dilutions: 1:150 dpERK (CST, 4370), 1:100 Arginase-1 (CST, 93668), 1:200 S100A2 (Abcam, ab109494), 1:150 Claudin 18.2 (Abcam, ab241330), 1:100 FAP (Abcam, ab207178), 1:150 Keratin-17 (CST, 12509), 1:200 TFF-1 (Abcam, ab92377), 1:150 Gata-6 (R&D systems, AF1700), 1:200 CD68 (Abcam, ab213363), 1:100 CD3e (Abcam, 16669), 1:100 CD8 (Thermofisher, MA5-14548), 1:300 CD206 (Abcam, ab64693),1:150 CD11c (CST, 45581), 1:100 Tenascin-C (R&D systems, MAB2138), 1:200 CD31 (Abcam, ab182981), 1:250 Pan-CK (CST, 3984). The secondary antibodies were used at the following dilutions: 1: 200 Donkey anti-Rabbit AlexaFluor Plus 555 (Thermofisher, A32794), 1:300 Donkey anti-Rabbit AlexaFluor Plus 647 (Thermofisher, A32795), 1:300 Donkey anti-Goat AlexaFluor Plus 647 (Thermofisher, A32849), 1:150 Donkey anti-Rat AlexaFluor Plus 555 (Thermofisher, A48270), 1:300 Goat anti-Syrian Hamster AlexaFluor 647 (Thermofisher, A21451).

### Image analysis

Image analysis was processed in QuPath v.0.6.0 (*72*). ROIs were drawn to include pancreas tumor sections and exclude other tissues on each slide. Cell-detection was conducted using QuPath’s built-in “Positive cell detection”. Distance between positive cells was calculated using the built-in “Distance to annotations 2D” function once populations were defined.

### Quantitative PCR (qPCR)

RNA was isolated from FACS-isolated cell populations using the RNeasy Plus Micro kit (#74034, Qiagen) per the manufacturer’s instructions. Cells were sorted into RLT buffer with beta-mercaptoethanol then frozen at - 80 C until isolation. RNA was isolated from FACS-isolated cell populations using the RNeasy Plus Mini kit (#74134, Qiagen) per the manufacturer’s instructions. cDNA was synthesized with the PrimeScript RT Reagent kit (#RR037B, Takara). qPCR was performed in technical triplicates with 1 µl of cDNA (diluted 1:10 if necessary) using the PowerUP SYBR mix (#A25778, Applied Biosystems) and analyzed on the QuantStudio 7 Flex Real-Time PCR System. The DDCT method was used to quantify relative gene expression, normalized to *GusB* or *ActinB*. The oligonucleotides used for qPCR amplification in this study are listed in **Table S4**.

### Validation of cytokine knockout

For protein level analysis, cell culture media were collected after 24-72 hr of conditioning, centrifuged to remove particulates and stored at –20 °C. The ELISA assays were conducted using mouse Duoset® ELISA kits (R&D Systems) according to the manufacturer’s instructions. In lines where protein level analysis was not feasible due to low baseline protein expression, genomic DNA was extracted from cells and region of the expected genomic disruption was amplified by PCR and purified using standard procedures. Purified PCR samples were sequenced by sanger sequencing. Genomic disruption was confirmed by analysis of sequences with a sequence trace decomposition tool (*73*).

### EVE Cytokine Profiling

Tissues were flash frozen in liquid nitrogen at the time of harvest and stored at −80°C. Tissue samples were manually homogenized in TG tube dounces in 30 µL of T-PER™ Tissue Protein Extraction Reagent (Thermo Scientific #78510), then lysed in a final volume of 100 µL T-PER supplemented with 1:100 protease inhibitor (Thermo Scientific #78441) for 20 minutes on ice. Tissue homogenates were centrifuged, and supernatants were collected. Total protein concentration was measured using the DC protein assay (Bio-Rad #500-0116). Samples were normalized to 1 mg/mL and submitted to Eve Technologies for the Mouse Cytokine Discovery Assay® (MD44 or MD68).

### Xenium data generation and analysis

FFPE blocks were sectioned and processed according to 10XG user guidelines (CG000580, CG000582, CG000584). Briefly, 5µm tissue sections were trimmed from FFPE blocks and placed within the fiducial frame of the Xenium slide (PN-1000460). The slides were air dried, baked at 42°C for 3 hours and stored in a desiccator. Tissues were then deparaffinized, rehydrated and decrosslinked using Xenium Sample Prep Reagents (PN-1000460). Tissues were hybridized overnight using a custom probe set (480 gene panel, ID#44CXV9). The probes were ligated and amplified *in situ*. Tissues were quenched to remove autofluorescence and counterstained with DAPI. Slides with corresponding decoding file were loaded and imaged on the Xenium instrument.

### Panel and gene set design

We designed a 480-gene Xenium probe panel to capture various PDAC transcriptional states, ligands and receptors enriched in basal cells, immune and stromal states relevant to the TME, as well as downstream signaling response genes to infer pathway activity (**Table S1**).

Our previous characterization of PDAC transcriptional states in this mouse model from Pitter *et al.* (*12*) guided selection of features that discriminate classical (*Lgals4*), basal (*Slc4a11*), and mesenchymal (*Lgals1*) states, in addition to ligands with state-biased expression, including *Pdgfb*, *Csf2*, and *Tgfb* for basal and *Ihh*, *Il18*, and *Wnt4* for classical states. To characterize non-cancer cells in the PDAC TME, we relied on several studies including Reyes *et al.* (*27*), Burdziak *et al.*(*74*), Schlesinger *et al.* (*75*), as well as in-house TME cell data from our model (not shown). Marker genes were selected to discriminate key cell types and previously described transcriptional subsets of TME cells (i.e. *Arg1*^+^ macrophages). Probe design was carried out with Schlesinger *et al*., Pitter *et al*., and in-house data covering a broad range of cancer cell and TME states, using the Xenium Panel Designer (10x Genomics) with assistance from the company. Genes that were too highly expressed and posed a risk of optical crowding were removed.

We designed gene sets in the panel to quantify levels of distinct transcriptional programs across individual cancer cells. Three gene sets defined the major PDAC cancer cell states based on overlap with established transcriptional subtypes in human PDAC studies, including Raghavan *et al*. (*76*), Moffitt *et al*. (*8*), and Collisson *et al*. (*7*) as well as our prior studies in mouse models of PDAC, including Pitter *et al*.: classical-PDAC (*Agr2*, *Cldn18*, *Epcam*, *Lgals4*, *Prom1*, *Spink4*, *Tspan8*), basal-PDAC (*Cd44*, *Cd9*, *Cldn6*, *Itga2*, *Krt17*, *Lamb3*, *Lamc2*, *Nt5e*, *Slc4a11*), and mesenchymal-PDAC (*Lgals1*, *Prrx1*, *Prrx2*, *S100a4*, *Stmn2*, *Twist1*, *Inhba*).

To characterize cytokine signaling pathway activity, we selected features from CytoSig (*77*), the Immune Dictionary (*78*), and ImmGen cytokine treatment studies (*79, 80*) that were relatively specific for cytokines linked to specific putative signaling pathways and were broadly induced in target cells. Negative feedback regulators of the pathway (e.g. *Socs3* for the STAT3 pathway) were prioritized, as they are often most strongly and immediately induced upon signal transduction, and are directly related to the reception of a specific signal. Nine signaling gene sets are represented in the panel, covering NF-kB, TGFB/SMAD, type I IFN, type II IFN, STAT3 related, Hedgehog, MAPK, IL4/STAT6 related, and WNT pathways. All gene sets include at least three genes, except WNT, which only consists of *Axin2* to focus on canonical pathway activity.

The panel also targets *CreER*, *GFP* and *tagBFP2*, which are important for phenotyping and confirming experimental consistency in our mouse model. Probes for these were designed with custom sequence input to the Xenium Panel Designer and assistance from the company.

### Spatial transcriptomics data preliminary processing

Spatial transcriptomics was performed on 15 tissue sections from untreated controls (*n* = 7), and from tissue collected at day 2 (*n* = 4), day 4 (*n* = 2), and day 6 (*n* = 2) after basal ablation using the Xenium platform (10x Genomics) with our custom panel. One sample comprised two ROIs (two separate Xenium images); therefore, metadata were organized by biological replicate (‘sample_name’) and a unique identifier for each Xenium image (‘sample_id’). Thirty-two probes were removed from analysis because they were not relevant to cell barcodes used in this study. Cell segmentations generated by the Xenium Onboard Analysis v3.2.07 software from nuclei (10x Nucleus) or cells based on ‘multimodal’ cell membrane staining (10x Cell) were converted to per-sample AnnData objects by summing transcript counts for each gene with a quality score (*qv*) > 25.

### Transcript assignment and cell segmentation with segger

Data derived from *in situ* hybridization-based gene expression profiling are rife with contaminating transcripts. Among other things, inaccurate cell segmentation, transcript diffusion, and z-plane contamination from overlapping cells can cause transcripts to be assigned to the wrong cell of origin. Although restricting analysis to transcripts that overlap well-defined DAPI signal (nuclear segmentation) is often considered ‘stringent’, nuclei are also contaminated by diffusing transcripts, and only contain a fraction of the cell’s transcriptome, leading to sparser and less accurate measurements.

We recently developed segger (*24*), a graph-attention network method that uses a heterogeneous cell-and-transcript graph to refine transcript assignments by learning similarities between transcripts and cellular boundaries in a joint embedding space. Segger leverages gene co-expression across the tissue as well as spatial proximity to inform similarity scores between transcript pairs and between transcripts and neighboring cells through its graph-attention network. Because the learned similarity scores are based on gene co-expression and physical position, they can more accurately assign transcripts to cells in the absence of defined cell membrane boundaries, and also refine mappings of transcripts that drift into neighboring nuclei or cells through diffusion or z-plane contamination. We used an updated implementation of segger (0.1.0) to refine our transcript-to-cell mappings (https://github.com/dpeerlab/segger).

We removed transcripts with *qv* < 25 and cells with fewer than 10 assigned transcripts based on DAPI-co-localized nuclear counts before entering raw Xenium data as input to segger, which was run for 20 epochs (or more if loss curves did not converge), using triplet loss, 128 dimensions, and max-nodes-per-tile of 55,000. Only cell IDs present in both 10x and segger processing were retained. Transcript outputs were summed for each cell ID and converted to AnnData objects, constituting 3,951,594 cells and 448 features across 15 samples as input for cell typing.

Segmentation quality was assessed by checking 1) the shape of the transcript similarity distribution across all transcripts, 2) the fraction of transcripts assigned to cells compared to 10x Nucleus and 10x Cell assignments, and 3) the removal of contaminant genes.

Only transcripts passing a segger similarity score threshold are assigned to a cell. Since segger is run, threshold values can vary for each transcript, as well as for the same transcript across samples, which allows segger to adapt to changes in gene co-expression and local contextual information. In representative samples, marker genes had high similarity scores in the cell type they marked, as expected (**Fig. S11A,B**). While thresholds for a given transcript differed by sample, the shape of the similarity distributions was largely the same, supporting the use of sample-specific thresholds to distinguish likely true counts from contamination.

The threshold for each gene can be tuned to achieve greater purity at the expense of total counts. To optimize this tradeoff, we varied similarity thresholds for a set of marker transcripts across representative samples, setting initial values as the minimum of Li (skimage.filter.threshold_li) (*81*) or Yen (skimage.filter.threshold_yen) (*82*) thresholding on the similarity score histograms (segger default), and removing unassigned transcripts. Additional thresholds were set by increasing the baseline threshold by 10–60% of the difference between the baseline and the maximum similarity value for that transcript, in 10% increments.

We selected six samples covering a range of cell type compositions, timepoints, and Xenium runs; and we used *GFP*, *tagBFP2* and *CreER* reporter transgenes that are expressed exclusively in pancreatic epithelial cells as ground truth for contamination, computing the fraction of contaminant transcripts out of total counts for all non-epithelial cells at each threshold. Segger-segmented counts generally exhibited far lower levels of transcript contamination, with similar or higher total count recovery than 10x Cell or 10x Nucleus (**Fig. S11C**). Increasing similarity score thresholds did remove additional contamination, but at the expense of total counts, which varied by sample. To maximize counts while dramatically limiting contamination, we therefore decided to use initial threshold values for our analyses.

On final segmentations, segger increased total counts 1.19–2.07 fold (median 1.51), while reducing mean contamination fraction to a median of 61% of contamination levels in 10x Nucleus segmentations (**Fig. S11D,E**).

### Sample quality control metrics

Per-sample quality control metrics were computed to assess data quality across 16 Xenium images (15 tissue sections) comprising 3,964,218 cells and 640 million transcripts. Metrics included median transcript counts per segger-assigned cell (130 ± 60, range 22–231), median counts per nucleus (59 ± 23), mean transcript quality value score (37.8 ± 0.5), and fraction of transcripts passing the segger similarity threshold (0.77 ± 0.05). Negative control metrics assessed background signal: mean control probe counts per cell (0.02 ± 0.02), mean control codeword counts (< 0.01), and fraction of non-specific transcripts (< 0.003). Non-specific transcripts were computed as the fraction of all transcripts that corresponded to the 32 probes in our panel that should not have cognate RNAs in these samples.

### Cell type assignment

We clustered all cells to assign ‘lineage’, then reclustered within broad lineages to assign ‘cell_annotation’, and finally within more granular lineages to assign ‘cell_annotation_granular’, and computed differentially expressed genes for each cell annotation compared to all others for that annotation resolution (**Table S2**). All analyses were performed using scanpy (v1.11.5), anndata (v0.12.11), Phenograph (*30*) (v1.5.7), and rapids-singlecell (v0.13.4) with GPU acceleration via cuML (v26.2.0).

### Assignment of major lineages

Cells were first assigned to epithelial, lymphoid, myeloid or stromal lineage, based on global clustering using PhenoGraph (*resolution* = 0.6) on log1p-normalized segger counts from all cells with at least 10 counts, and principal components (PCs) explaining 80% of variance. Lineage assignments were based on marker genes: *Epcam*, *GFP*, *tagBFP2*, *Tubb3* (epithelial/neuronal); *Vim*, *Col5a1*, *Kdr*, *Pecam1*, *Pdgfrb* (stromal); *Ptprc*, *Cd3e*, *Cd19*, *Il7r*, *Mzb1* (lymphoid); and *Ptprc*, *Csf1r*, *Itgam*, *Csf3r* (myeloid).

#### Common sublineage preprocessing pipeline

Cells within each lineage were clustered for more refined cell typing. We filtered segger counts to retain cells with total counts ≥ 20 and total genes ≥ 10—relatively permissive quality control filters that ensured cell types with low library size (i.e. neutrophils) were not excluded. Transgene counts were removed prior to dimensionality reduction, so that clustering would only reflect endogenous gene expression. Dimensionality reduction and PhenoGraph clustering were performed in each lineage using PCs explaining 70% variance on log1p-normalized segger counts, *k* = 50, UMAP with *min_dist* = 0.1 and 2000 epochs, Leiden clustering *resolution* = 1 and *min_cluster_size* = 150 cells. Additional PhenoGraph clustering at *resolution* = 10 was used for majority voting approaches at later stages of quality control (overclustering).

Cells were flagged as putative low quality if more than 50% of cells within the high resolution PhenoGraph clusters had fewer than 25 counts, and they were subsequently removed if they did not share biological structure, such a neutrophil grouping, and lacked distinct marker genes. Populations that co-expressed marker genes of different lineages were denoted as ‘contaminated’ and removed from analysis. For each lineage, differential expression within clusters was performed with Wilcoxon rank-sum tests on log1p-normalized segger counts in scanpy (sc.tl.rank_genes_groups), and results were filtered by log_2_ fold-change ≥ 0.5 and Benjamini-Hochberg-adjusted *p* < 0.01. Cells were identified as ‘cancer-neighbors’ if they were within 60 µm of a cancer cell, which helped to identify cancer-associated populations.

Because samples were collected at different times, we used annotations from an initial clustering to classify subsequent samples using label transfer by a Celltypist (*83*) model trained on our Xenium data at that stage in sample collection. The model was trained in a standard way, using a 90 train : 10 test split stratified by cell type on segger counts that were library size–normalized and multiplied by a size factor of 10,000 followed by log1p transformation. All panel features except *GFP*, *tagBFP2* and *CreER* were used as input. The majority *predicted_label* within overclustering-defined clusters was used for downstream analysis and refinement.

### Epithelial/neuronal lineage annotation

Epithelial/neuronal cells were manually annotated as acinar, ductal, acinar-to-ductal metaplasia (*30*), cancer epithelial, mesothelial/antigen-presenting CAF (apCAF), tuft-like, islet, Schwann cell, and neuron based on marker gene expression within defined clusters. Marker gene expression patterns were derived from our previous work and other PDAC scRNA-seq datasets(*12, 74, 75, 84*). Marker genes included *Phgdh*, and *Igf1* (acinar); *Pou2f3* (tuft-like); *Il6ra* and *Ncam1* (islet); *Fgfr3*, *Prox1*, and *Sox9* (ductal); *Msln* and *H2-Aa* (mesothelial/apCAF); *Slc4a11*, *Lgals4*, *Lamc2*, *Onecut2*, and *Kras* (transformed cancer epithelial cells); and *Tubb3*, *Trpv1*, *Stmn2*, *S100b* (neuronal-associated). ADM cells were identified as a population bridging normal acinar cells and malignant/ductal populations with high *Lcn2*, *Sox9* and *Kras* expression, that retained *Igf1* and reduced *Phgdh* acinar marker expression. In addition, we identified a population of cancer epithelial cells with high neuron-associated gene (*Trpv1*, *Tubb3*, *L1cam*) expression that occurred across multiple samples, which we annotated as cancer epithelial neuronal-like. It is unclear whether this represented a highly enervated population or the cancer cells themselves expressed some of the marker genes.

Following general annotations, several sublineages were clustered further. Antigen-presenting CAFs were difficult to differentiate from mesothelial cells (from which they derive) and were thus annotated as mesothelial/apCAF. These cells expressed *Msln*, *Saa3*, *Sdc4*, and *H2-Aa*. Neuron and glia-associated cells were subclustered (50% variance PCs, *k* = 50, PhenoGraph *resolution* = 0.8, *min_cluster_size* = 300). Schwann cells were identified by S100b and L1cam expression, while neurons were identified by high *Tubb3*, *Stmn2*, and *Ngfr*. Cancer epithelial cells were further annotated into granular states (see ***Cancer cell state assignment***).

### Lymphoid lineage annotation

Lymphoid cells were separated into several sublineages for clustering. Natural killer (NK) and innate lymphoid cells (ILC1) were grouped due to transcriptional similarity. NK_ILC cells expressed *Ncr1, Eomes, Klrk1,* and *Gzmc*. We also identified other *Cd3e-*negative ILC populations expressing high levels of *Il7r*, including ILC2s (*Gata3, Il1rl1*) and ILC3s (*Rorc, Ccr6, Kit, Il22*).

We utilized our CellTypist predictions described previously to annotate T cells, as this discriminated T-cell subsets better than standard clustering. T cells were annotated into conventional CD4^+^ T cells (Tconv; *Cd4*, *Cd3e, Trac*), CD8^+^ T cells (*Cd8a, Cd8b1, Cd3e, Trac*), Treg (*Foxp3, Ctla4*), gamma delta T (gdT; *Trdc, Rorc, Il23r, Trac-*), and mucosal-associated invariant T cells (MAIT; *Trdc-, Trac, Cd3e, Rorc, Il23r, Cxcr6*). An additional population of innate-like T cells expressed *Trac* and *Cd3e* but was *Cd4-* and *Cd8a-*negative and expressed *Cxcr6, Zbtb16,* and *Klrk1*.

B-cell subclustering (49 PCs explaining 60% variance, *k* = 30, PhenoGraph *resolution =* 0.8, *min_cluster_size* = 1) yielded B cells (*Cd19* and *Cd79a*) and closely related plasma cells (*Mzb1* and *Irf4*). Plasma cells contained more mature plasma cells as well as plasmablasts (*Mzb1* expression and proliferation markers) as well as transitional plasma cells that expressed *Mzb1* but had not yet downregulated *Cd19*. We also identified lymphoid cells of mixed lineage expressing proliferation markers (*Mki67, Pcna)*, which we grouped and annotated as lymphoid_proliferating.

### Myeloid lineage annotation

Myeloid cells were split into macrophage/monocytes, conventional dendritic cells (cDC), and neutrophils/granulocytes for subclustering. Plasmacytoid dendritic cells (pDC) can derive from myeloid and lymphoid lineages and were sufficiently distinct to be annotated in isolation (*Bcl11a, Irf8, Mzb1*).

cDCs were isolated from other myeloid cell types by high *H2-Aa, Itgax,* and *Zbtb46* expression. These were divided into subsets using marker genes *Xcr1* and *Irf8* (cDC1); *Irf4, CD2091,* and *Sirpa* (cDC2) and *Ccr7* (mature DC enriched in immunoregulatory molecules (mregDC)). cDC2 populations also displayed features of monocyte-derived dendritic cells (moDC), including *Itgam* and *Ccr2;* however, we lacked marker genes to completely distinguish these subsets and the cDC2 annotation therefore likely encompasses both subsets.

Neutrophils were identified as a population with high *Csf3r, Itgam* and *Il1r2* expression and lower library size than other myeloid populations. A small population (∼500 cells) that was transcriptionally similar to neutrophils expressed *Kit* and *Il1rl1*, suggesting it could be a distinct population such as mast cells, but did not have enough cells for robust phenotyping and was therefore labeled as general granulocyte.

Macrophages and monocytes were isolated as a large population expressing higher levels of *Aif1, Mrc1, Mafb,* and *Csf1r* relative to other myeloid cells (**Fig. S12**). Macrophages and monocytes were combined and labeled macrophage (the dominant population) due to their transcriptional similarity and the difficulty of separating them within our data. We subclustered these cells (48 PCs explaining 60% variance, *k* = 30, PhenoGraph *resolution* = 0.8) to assign granular state labels based on the expression of key marker genes in our panel. We identified populations which mainly resided in the lymph node, neighboring cancer cells, or non-cancerous tissue; however, transcriptional states were not definitively restricted to these boundaries. A population largely associated with lymph node included macrophages expressing *Timd4* (macrophage_Timd4). Interstitial populations, those reflecting populations typically present in normal pancreatic epithelium and along connective tissue of the pancreas, were identified by less frequent contact with cancer cells (within 60 µm) and higher expression of *Lyve1, Folr2, H2-Aa,* and *Clec10a* (ref). Several interstitial populations were identified, including those expressing interstitial markers alone (macrophage_interstitial) or with *Cd163* (macrophage_interstitial_Cd163) or *Retnla* (macrophage_interstitial_Retnla). Among the interstitium-associated macrophages, macrophage_interstitial_Cd163 was most frequently associated with cancer cells and expressed lower levels of *H2-Aa*. One macrophage population had an equal number of neighbors and non-neighbors to cancer cells and expressed low levels of interstitial associated genes. This population was identified by high expression of *Saa3* and labeled macrophage_Saa3.

Macrophage populations that tended to localize near cancer cells were termed tumor-associated macrophage (TAM). *Arg1*^+^ TAMs (TAM_Arg1) expressed *Arg1, Itgam,* and *Plaur*, and similar populations expressed high levels of additional marker genes including *Axl* (TAM_Axl), *Fabp5* (TAM_Fabp5), AP1 components *Junb* and *Fosl2* (TAM_AP1), and *C1qb*, which was more similar transcriptionally to interstitial populations (TAM_C1q). Other TAM populations expressed lower *Arg1* and higher levels of *Itgax* and *H2-Aa* (TAM_Itgax), or neighbored cancer cells highly expressing interferon-stimulated genes (ISGs) including *Irf7, Rsad2,* and *Stat1* (TAM_IFN).

### Stromal lineage annotation

Fibroblasts and pancreatic stellate cells (PSCs) were annotated using consensus signatures from the PDAC literature (*85–87*). Fibroblasts were identified by *Vim*, *Pdpn*, *Col5a1* and *Col15a1* expression. After subclustering (70% variance PCs, *k* = 50, PhenoGraph *resolution* = 0.8, *min_cluster_size* = 100), granular annotations were first assigned using myofibroblastic CAF (myCAF) and inflammatory CAF (iCAF) signatures from Elyada *et al*. (*85*). This identified a myCAF population expressing high levels of *Tnc* and *Fap* (myCAF_FapTnc), and related states expressing either high levels of Wnt receptor *Fzd1* and downstream target *Axin2* (myCAF_Wnt), or hedgehog signaling–related genes *Gli1* and *Ptch1/2* (myCAF_Gli1). We found a population mapping to the iCAF signature that expressed *Dpt* and *Pdgfra* which we called iCAF, a population resembling adventitial fibroblasts and expressing high levels of *Pi16* (fibroblast_Pi16), and a population similar to fibroblastic reticular cells (FRCs) that was mainly located in lymph nodes and expressed high levels of *Cxcl12* and *Cxcl13* (fibroblast_FRC-like). Additional populations expressed high levels of interferon response genes *Rsad2* and *Irf7* (fibroblast_IFN); *Saa3* (fibroblast_Saa3); or *Serpine1* (fibroblast_Serpine1); or lacked clear marker genes (fibroblast_unknown).

PSCs clustered separately from traditional fibroblasts, and expressed high levels of *Pdgfrb*, *Notch3*, and *Acta2* and lower levels of collagen components *Col5a1* and *Col15a1*. Vessel-associated mural cells expressed high *Notch3*, *Acta2*, and *Eln* but lower *Pdgfrb* levels compared to PSCs. Other *Notch3*- and *Pdgfrb*-negative populations associated with a PSC/mural cell cluster were defined as mesenchymal-like stroma because they lacked clear defining markers, but they expressed higher levels of *Itga5* and *Itgb1*; subpopulations of these expressed high levels of *Fgfr2* or localized near blood vessels. We also identified mural cells that clustered more closely with vascular or lymphatic endothelial cells. Pericytes clustered with vascular endothelial cells and expressed high levels of *Notch3*, *Acta2*, and *Pdgfrb*. Smooth muscle cells (SMCs) formed a distinct population that clustered closest to vascular endothelial cells and expressed high *Acta2* and *Stmn2,* but lacked traditional endothelial markers. Mural cells that clustered more closely with lymphatic endothelial cells expressed high levels of *Pdgfra*, *Pdgfrb*, and *Notch3* while lacking expression of *Pecam1*.

Endothelial cells were identified by *Pecam1* and *Kdr* expression. Capillary-associated vascular endothelial cells expressed *Ackr1* and *Plvap* and lymphatic endothelial cells expressed *Prox1* and *Lyve1*.

Adipocytes were mapped to a small population that clustered closely to endothelial cells, expressed *Fabp4* and lacked traditional endothelial markers.

Mesenchymal cancer cells clustered closest to fibroblasts in our data. These could be distinguished based on the expression of PDAC mesenchymal markers *Vim*, *Prrx1*, *Prrx2*, *Twist1*, and *Lgals1*. Moreover, mesenchymal cancer cells express our GFP reporter, because they derive from the epithelial lineage, facilitating their distinction from traditional fibroblasts.

### Cancer cell state assignment

While cancer cell states fall along a continuum, assigning broad classical, basal, or mesenchymal labels was useful for tracking the frequency of these categories after basal ablation or in relation to our niche embeddings, and for restricting and filtering our data for targeted analysis (see ***niche trajectory pruning*** sections). To do this, we assigned labels to one of the three main states using a CellTypist classifier trained on cancer cells from untreated samples (*n* = 7). Untreated samples were used to prevent basal ablation–induced variation from affecting our annotations, thus giving a more common reference across conditions.

In practice, mesenchymal cancer cells clustered distinctly from epithelial states, such that cluster membership was sufficient for their annotation. Epithelial cancer states failed to form a distinct population upon clustering, so they required the classifier model to distinguish them accurately. The classifier was still trained with mesenchymal cells to confirm our clustering approach, and to allow us to assess mixed cancer states (i.e. basal with mesenchymal features).

To select cells and features for training and assign labels, we concatenated cancer_epithelial and cancer_mesenchymal cells and subset to untreated samples (520,181 cells from 7 samples). We created a knn graph (*k* = 50) using PCs explaining 50% variance and then computed diffusion maps (*29*) using Palantir (*88*) to capture the main axes of variation among cancer cells. A small population of *Spink4-* and *Tspan8*-high cancer cells were removed before computing the final DCs. Pearson correlations between DC values and gene expression revealed that DC1 defines an epithelial–mesenchymal axis that is positively correlated with mesenchymal markers *Vim, Prrx1,* and *Twist1* and most negatively correlated and *Epcam*. DC2 was found to represent a classical–basal axis positively correlated with *Lgals4, Tspan8,* and *Agr2*, and negatively correlated with *Slc4a11, Sox11,* and *Lamc2*. Training labels were assigned using DC thresholds to capture cells that were not co-expressing multiple cancer state programs: mesenchymal (DC_1 > 0.75), basal (DC_2 < -0.30 and not mesenchymal), classical (DC_2 > 0.20 and not mesenchymal). In order to only train on genes most informative to cancer state, the top 20 positively and negatively correlated genes with each DC (57 unique genes) were selected as features for training.

To assign cancer state labels to all 671,050 cells in the study, including cells from post-ablation conditions, a CellTypist logistic regression classifier was trained using an 80 train : 20 test split to confirm performance, achieving 99% accuracy. The *predicted_labels* output from CellTypist was used for state assignment, representing the cancer state with highest classification probability among cancer epithelial cells. The final assignments resulted in 353,751 classical, 146,975 basal, and 170,324 mesenchymal cells.

### Spatial neighbor graph construction

We constructed spatial neighbor graphs to analyze cellular niches. Our analyses either used a fixed 60-µm radius or the 32 most spatially proximal neighbors (specified in each relevant analysis section below). The 32 nearest-neighbor resolution was used for our Wormhole niche embedding, to focus on local relationships (∼35-µm radius) and reduce niche overlap, and the 60-µm radius was used when further reductions in sparsity were desired, such as computing ligand–receptor correlations across individual niches, or for stringent filtering (***Niche trajectory pruning*** sections). Spatial neighbor graphs were constructed using Squidpy (*89*) with squidpy.gr.spatial_neighbors. Per-cell neighbor counts by cell type were computed by summing connections across each cell type’s columns in the connectivity matrix. Neighbor fractions were computed by dividing by the total number of neighbors.

### Selection and exclusion of tissue areas

To focus analyses on the pancreatic tumor microenvironment and exclude lymph node regions, we identified lymph node and parenchymal regions per sample with a strategy similar to the *in-silico* dissection performed in Reyes *et al.* (*27*). Parenchyma was defined as all epithelial cells (acinar, ADM, cancer_epithelial, cancer_mesenchymal, ductal, islet, tuft-like) plus any cell within 200 µm of an epithelial cell. Only 2% of cells (66,509 cells) were outside the parenchyma by these criteria. Lymph nodes were identified by constructing a spatial neighbor graph of all cells within a 30-µm radius of those cell types known to be overrepresented in lymph nodes relative to parenchymal tissue (B_cell, plasmablast, CD8_T, Tconv, Treg). Using this neighbor graph, we defined groups of the selected immune cells that were spatially connected (i.e., connected components; scipy.sparse.csgraph.connected_components), and classified components exceeding 250 cells (i.e., dense aggregates) as lymph nodes. Lymph node annotations were expanded by marking all cells within 50 µm of identified lymph node cells to ensure exclusion from our niche analysis. This identified 460,881 cells (15% of total) in 6 of 15 samples as residing within a lymph node area.

### Differential expression

We tested for differential gene expression between control (day 0) and basal ablation (days 2–6) tissue sections separately in macrophages and fibroblasts to understand expression shifts after basal cell removal **(Fig. S4A**). Differential expression was performed with edgeR’s quasi-likelihood test (*90*) using segger counts summed across cells of each type within each sample (*n* = 5 control; *n* = 8 DT treated; 128,612–313,098 cells per condition per cell type). Size factors were estimated via calcNormFactors() to use as an offset during testing, followed by dispersion estimation and quasi-likelihood model fitting (glmQLFit). FDR correction was performed using the Benjamini-Hochberg method.

To reduce spurious associations due to contaminating transcripts, we annotated each gene with the percent of cells that expressed it for each cell type (at least 1 count). To highlight genes associated with basal niches, we computed Spearman correlations to the untreated niche embedding DC1 axis (see ***Gene–trajectory correlations***), and identified basal-correlated genes as those with *ρ* > 0.7. Volcano plots highlighted genes that were differentially expressed at an adjusted *p* < 0.1, log_2_ fold-change > 0.5, and were expressed in at least 4% of cells (42 and 71 genes in fibroblasts and macrophages, respectively, out of 448 panel genes tested).

### Generation of niche embeddings with Wasserstein Wormhole

*Overview.* Niches are inherently distributional entities. Each cellular neighbor expresses gene programs to different extents, which cannot be fully captured using discrete averages. Moreover, tissue remodeling after cancer cell state ablation is a highly dynamic process in which niche structure changes gradually in response to the perturbation, requiring a niche similarity metric that can faithfully incorporate such complexity for inferring remodeling trajectories. Optimal transport–based methods enable the use of similarity (distance) metrics that retain the distributional nature of the underlying measurements and thus more accurately reflect spatial biology, but they are prohibitive to compute for tens of thousands of pairwise distances, let alone the millions in our study. We thus employed Wasserstein Wormhole (0.3.0, https://github.com/dpeerlab/WassersteinWormhole)(28), our transformer-based autoencoder that embeds cells in a latent space wherein Euclidean distances approximate optimal transport distances, allowing for a dramatic reduction in computational cost. Using this approach, we were able to relate the niches of cancer cells to one another, either across the classical–basal axis in untreated tumors, or in response to cancer cell-state ablation, constructing continuous trajectories of cellular recruitment and remodeling (**Fig. 4A,B**).

We used a modified implementation of Wormhole that includes changes to feature scaling and distance computation, and that incorporates gene program–based modules rather than individual genes as input features for greater robustness (details of the implementation in this study appear here; a full description and benchmarking of the generalized spatial ecosystem framework will be provided separately; reflected in current GitHub version 0.3). To construct gene modules, we first computed local gene–gene correlations in transcriptomic space using Hotspot (*91*), then partitioned the correlation matrix into modules using a symmetric version of non-negative matrix factorization (symNMF) (*92*), allowing for gene membership in multiple modules. Next, we computed PCs within each module across all input cells (which we term modPCs) and used these per-cell loadings to generate Wormhole models.

#### Wormhole analysis details

Modules were computed for all cell types and timepoints at once (downsampled to 50% to reduce computational complexity), defining global groups of co-expressed genes across cells. For Hotspot, we used a depth-adjusted negative binomial model with raw segger-assigned counts as input. An unweighted knn graph (*k* = 300) was constructed using distances based on 93 PCs (explaining 70% of variance in log-normalized expression space). Moran’s I autocorrelation identified locally variable genes with FDR < 0.05 out of all panel genes (445 genes), and local correlations were computed for these genes and clipped to zero for further analysis. NMF was run on the resulting local correlation matrix with *k* = 25 modules (*92*). For each gene, the loading values were square-sum normalized and assigned to all modules up to α = 0.4, a cumulative sum threshold determined to balance the specificity of gene modules with the pleiotropic activity of certain genes.

modPCs were computed across all input cells to our untreated or ablation response analysis separately using the same defined set of 25 gene modules. Principal component analysis was applied independently within each module to generate per-cell features, by subsetting cells to module genes and computing PCs on log-normalized expression using randomized singular value decomposition without centering. The resulting modPCs, explaining up to 70% variance, were retained and concatenated to form the final feature matrix.

Separate Wormhole models were trained for untreated and ablated conditions, based on niches containing 32 nearest neighbors (and excluding the anchor cell itself). We selected *n* = 24 latent dimensions in both models to capture relevant biological variation without introducing sample-specific artifacts, which can arise due to tumor compositional heterogeneity. As input, we used all parenchymal, non-lymph node anchor cells. In our more targeted analyses of cancer cell niches, we filtered anchor cells after model training to focus on specific continuous relationships of interest.

### Construction of niche trajectories

To characterize continuous niche remodeling, we used Wormhole embeddings as the substrate for trajectory analysis, focusing on the classical–basal niche axis in untreated tumors, and the classical response axis after basal-state ablation. A single tissue section contains niches at different stages of remodeling. By ordering niches directly in the learned ecosystem embedding space, we can reconstruct an unsupervised progression of remodeling states, identify graded transitions, reduce the impact of sample-to-sample variability, and aggregate niches at comparable stages of response for downstream analysis. To focus trajectory inference on tumor-associated epithelial remodeling, we restricted anchor cells to malignant and pancreatic epithelial populations. This reduced variation driven by stereotyped anatomical localization of unrelated cell types, such as mesothelial cells at tissue boundaries, while preserving the stromal and immune cells present within each epithelial-centered niche.

Because cancer cell transcriptional state is itself a major determinant of local niche structure, we applied trajectory-specific filters to isolate the remodeling axis relevant to each analysis. For the untreated classical–basal trajectory, we excluded niches with substantial mesenchymal cancer-cell content, allowing the analysis to focus on the epithelial classical–basal continuum. For the post-ablation classical response trajectory, we excluded niches containing basal or mesenchymal cancer cells, thereby isolating the secondary remodeling of residual classical niches after basal-state loss. These filters enabled diffusion components to recover the principal axes of niche variation corresponding to the biological processes under study.

### Untreated niche trajectory pruning

The untreated Wormhole model consisted of 1,231,448 total cells across 7 samples. Samples BT1791, BT1793, and Br1436 were excluded from the untreated trajectory analysis because they were dominated by extensive mesenchymal cancer-cell expansion and formed sample-specific regions in the niche embedding, rather than mixing with the shared classical–basal continuum observed across the remaining tumors.

To determine the untreated classical–basal niche trajectory, we first selected epithelial cancer cell anchors from the untreated Wormhole embedding, yielding 233,694 classical and 119,276 basal candidate anchors. To focus on resolving the classical–basal continuum, we excluded niches with >10% mesenchymal cancer cells within a 60-µm radius, removing approximately 25% of candidate anchors with substantial mesenchymal content, while retaining niches with low-level mesenchymal admixture that may represent early or local mesenchymal emergence. Following filtering, 90,903 niches were retained for analysis.

We visualized the niche embedding using UMAP with *k* = 50 neighbors defined by our 24-dimension niche latent. To remove small, disconnected islands that can distort diffusion component analysis, we applied HDBSCAN (*93*) clustering on the UMAP (*min_cluster_size* = 25, *cluster_size_threshold* = 500) and retained the large, connected niche population. This removed 1,592 niches, leaving 89,311 niches.

### Ablation trajectory pruning

To analyze niche remodeling after basal-state ablation, we reapplied Wormhole to samples collected at day 0 and days 2, 4, and 6 after DT treatment. The initial embedding included 2,117,666 cells from 13 samples. Samples BT1791, BT1793, B0 1401 1.veh, and BQ1955 were excluded because they were dominated by mesenchymal cancer-cell expansion or showed sample-specific response patterns that did not mix with the shared ablation trajectory, leaving 1,528,897 anchor cells across 9 biological replicates. For the global epithelial remodeling analysis, we selected all malignant and non-malignant pancreatic epithelial anchor cells from this ablation embedding (795,402 cells total), down-sampled to 25% to reduce computational cost, and visualized the 24-dimensional Wormhole latent space by UMAP using *k* = 75 neighbors. HDBSCAN clustering of the UMAP coordinates (*min_cluster_size* = 10, *cluster_size_threshold* = 300) removed 37 cells, forming a small, disconnected island, and the resulting embedding was used to visualize epithelial states, treatment timepoints, and niche composition (**Fig. 6B**).

To define the classical response trajectory after basal-state ablation, we focused on a stringent set of classical epithelial anchor cells, with the goal of resolving secondary remodeling of residual classical niches, rather than the direct effects of basal or mesenchymal cancer-cell loss. We therefore started from classical-labeled anchor cells in the non-outlier ablation samples (155,903 cells) and excluded any niche containing basal or mesenchymal cancer cells within 60 µm, retaining 59,243 anchors. Cancer-state labels only approximate an underlying continuous transcriptional space; thus, we further removed classical-labeled cells with evidence of mixed classical–basal identity, defined by a basal-PDAC signature score > 0.2. We also removed an *Agr2*-high gastric-like epithelial cluster, which represented a distinct epithelial program. After these filters, 47,825 classical anchor cells remained. UMAP was performed on the neighbor graph (*k* = 50) in the 24-dimensional Wormhole niche latent space, followed by HDBSCAN filtering of small off-trajectory islands (*min_cluster_size* = 25, *cluster_size_threshold* = 300). This removed 4,285 cells, leaving 43,540 classical niches for trajectory analysis.

### Diffusion component analysis

Following niche selection, we computed diffusion maps on the niche neighbor graph using Palantir (v1.4.4). An adjacency matrix was computed from the 24-dimensional Wormhole niche latent space using *k* = 50 neighbors, and the top 20 DCs were extracted. We applied the Palantir multiscale transformation, in which each eigenvector is scaled by λ/(1−λ), where λ is the corresponding eigenvalue, and the trivial first eigenvector is discarded. This multiscale representation avoids selecting a single diffusion scale and is particularly important for manifolds with large density differences, where abundant regions and rare remodeling states may otherwise be resolved at different optimal scales. The resulting scaled DCs captured continuous gradients in niche composition, with the first diffusion component (DC1) corresponding to a biological axis of interest (untreated classical–basal or post-ablation classical response axis). DC values were rank-transformed for visualization and downstream analysis to represent ordered progression across niches (**Fig. 4B**, **Fig. 6B**).

### DC1 bin summarization for trajectory analysis

To reduce sparsity in our trajectory analysis, we divided DC1 into 30 equally spaced bins, yielding ∼3,000 and ∼1,450 cancer cells per bin in the untreated and classical response settings, respectively. For each DC1 bin and cell type, counts were summed across cells, normalized to the total counts for that cell type, scaled to the median total count across bins, and log1p-transformed. The frequency of receptor-positive cells for each cell type out of all niche bin cells was also computed. A threshold of at least 2 counts was used to limit noise caused by segmentation artifacts (discussed in ***Transcript assignment and cell segmentation with segger***).

To summarize TME gene programs along each cancer-cell-defined trajectory, we selected all non-anchor cells in niches along the trajectory, and assigned each cell a DC1 value. The DC1 value was averaged from all cancer anchors within 60 µm of the cell, weighted by inverse physical distance, to account for membership in multiple niches. This value was used to determine which bin the cell occupied (bin boundaries were defined by all cancer-cell anchors). We quantified the fraction of each cell type out of all cells in each DC bin (**Fig. 4C**, **Fig. 6D, Fig. S3C**), or the fraction of granular macrophage and fibroblast states out of the total number of cells of that type in each niche bin (**Fig. S7E**).

### Gene–trajectory correlations

Spearman correlations were computed between DC1 bin position and summarized gene expression values in each cell type for all panel genes. The top positively and negatively correlated genes (*ρ* > 0.7) for each DC were identified to characterize the biological processes underlying each trajectory axis (**Fig 4.D-F, Fig. S3E-G, Fig. S7B,C**). Correlations were additionally filtered for genes with > 0 counts detected in at least 5% of cells for a given cell type to remove spurious associations.

### Ligand and receptor co-localization analysis

To prioritize ligand and receptor pairs that 1) covary along key axes of variation and 2) spatially co-localize in the tissue, we undertook a two–tier testing strategy.

Tier 1 uses cell type expression matrices in equally spaced bins along an axis—either the untreated classical–basal or classical response axis—to measure relationships between ligand–receptor (L–R) pairs along the trajectory. For each ligand and cognate receptor and ordered sender–receiver cell type pair, we fit a linear regression across DC1 bins. The predictor is the log-normalized expression of the ligand in the sender cell type within each bin, and the response is the fraction of receiver cells with detectable receptor expression within the same bin. This test reduces sparsity by summarizing across cells in each bin, and it groups niches relative to the axis of study, thus quantifying putative L–R interactions in relation to the biological variation we seek to explain.

Tier 2 measures spatial co-localization between L–R pairs across niches, in order to verify that the L–R associations which covary along our axis of interest also physically associate in the same niches (bin-level summarization in Tier 1 does not guarantee spatial association). To test this, we summarized ligand expression within niches by cell type, then computed the Spearman correlation between log-normalized ligand expression and the fraction of receptor-positive cells out of all cells in the niche, for each sender–receiver pair. We computed correlation values within each sample, to ensure consistent co-localization across biological replicates. For this analysis, we used cells within 60 µm of anchor cells (∼100 neighbors per anchor) to mitigate sparsity, rather than the 32 neighbors used to compute the niche embedding.

In the untreated setting (**Fig. 5A, S5A,B**), we tested 70 ligands and 63 receptors across 16 cell types (at least 500 cells per cell type), totaling 10,672 tests. We found 625 associations at a Benjamini-Hochberg–adjusted *p* < 0.01 in the tier 1 test, which was filtered for ligands and receptors expressed in >10% of cells of each type to remove low-frequency interactions. After applying our tier 2 filter of within-niche *ρ* > 0.1 (90th percentile) in at least 2 samples, 378 were removed, leaving 247 prioritized interactions representing 79 different L–R pairs. We focused on ligands whose expression was correlated with DC1 of the untreated niche embedding (Pearson *r* > 0.1), leaving 16 basal-enriched ligands with 63 L–R interactions and 5 classical-enriched ligands with 15 L–R interactions.

In the classical response setting (**Fig. S8B**), the total number of cancer anchor cells in the response trajectory was 43,540 (approximately 1,500 per bin), and the last 2 bins of DC1 were removed from analysis due to sample-specific behavior. We tested L–R relationships among 77 ligands and 73 receptors across 29 cell types (at least 200 cells per cell type), totaling 22,019 tests. The tier 1 test yielded 577 associations at an adjusted *p* < 0.05, and tier 2 filtering removed 317, leaving 260 significant associations, which were further filtered for receptors or ligands detected in at least 5% of cells of a given type. One association that was just below our receptor-positivity threshold was included (detected in 4.5% of macrophages)—*Mif* sent by cancer cells to *Cxcr4* in macrophages—because *Mif* expression in cancer cells is strongly associated with the response axis (*ρ* > 0.97), and the pair passed the tier 2 co-localization test in 4 separate samples. In addition, NK_ILC1 cell associations were retained if they had a mean *ρ* > 0.05 across samples for the tier 2 test, as they likely suffer from very few cells when conducting the co-localization test.

For the classical response axis, we also visualized total ligand expression and receptor-positive cell fraction from niche bins summarized across all cell types at once, in order to reduce sparsity and understand how total levels of certain chemokines related to cell-type-specific expression patterns. We did an additional Tier 1 test at the niche level to confirm chemokine L-R associations at this level (**Table S3**).

#### Gene associations with diffusion components

Along each niche trajectory, we prioritized genes whose expression was strongly associated with DC1 (either increasing or decreasing along the trajectory), based on Spearman correlation between bin-summarized per-cell-type expression for each gene and the ordered DC1 bin number.

In the untreated setting (**Fig. 4D-F; Fig. S3E-G**), 336 of 1334 genes tested per cell in cancer cells, macrophages, and fibroblasts had *ρ* > 0.7 and adjusted *p* < 0.01. These were filtered for ligands expressed in at least 10% of cells within a given cell type to remove spurious associations potentially driven by contamination.

In the classical response setting (**Fig. S7B,C**), 159 of 947 genes tested per cell type had *ρ* > 0.7 and adjusted *p* < 0.01. These were filtered to retain ligands expressed in at least 5% of cells within a given cell type. The lower threshold was designed to enhance capture of cytokines and other ligand genes that are lowly expressed but critical for immune response. We included aggregated type I IFN ligand correlations as well, despite their detection in only ∼1% of cells, as type I IFN activity was supported by associations with downstream response programs.

### Ligand associations with response programs

Our panel included sets of downstream response genes for signaling pathways (see ***Panel and gene set design***). To assess the activity of these gene programs, we *z*-scaled summarized expression values across bins for each gene, then averaged all genes in each response gene set to obtain a signature score. Response programs for cancer cells, fibroblasts, and macrophages correlated with DC1 (*ρ* > 0.5) were highlighted (**Fig. S7D**).

#### Differential effector molecule analysis in classical response

To determine which molecules could be mediating immune activity against classical cells, we compared the frequency of effector molecule–expressing cells in early (bins 3–13) and late (bins 17–27) portions of the DC1 trajectory. These ranges were selected to capture niches in windows prior to most remodeling or after basal ablation effects, while avoiding edge cases by excluding the bins at the extremes. To identify shifting genes, we performed Fisher’s exact tests on pooled counts to maximize power, given that many ligands critical for anti-tumor immunity (i.e. *Ifng*) are poorly captured at the transcript level. For each cell type and gene combination, the sum of cells with ≥ 2 counts for the gene within each early or late group was used to create a 2 x 2 contingency table, before applying the Fisher’s exact test (reporting Benjamini-Hochberg-corrected *p* values) and computing odds ratios using Haldane-Anscombe correction to avoid errors from 0-count cells (odds ratio > 1 indicates an increase in late bins compared to early) (**Fig. S8C**).

### Gene or cell fraction plotting over DC bins

When visualizing L–R pair expression with heatmaps, we smoothed expression values to reduce noise driven by discrete bin cutoffs, by taking the mean of expression between each bin and its two flanking bins, then *z*-scaled values per gene. To order ligands in each receptor group, we took the first derivative of the smoothed expression across bins and identified the first bin in which the derivative exceeded 0.05 times its maximum absolute value. Ligands that were negatively associated with the DC were plotted first.

We also used line plots so summarize trends of gene expression or cell type fractions along the classical response axis (**Fig. 6E,F; Fig. S7E**). Line plots were generated using loess smoothing (*span* = 0.3) on bin summarized gene expression or cell frequencies in the ggplot2 (v4.0.1) stat_smooth function. Shading indicates the 95% confidence interval.

### scRNA sequencing using the 10x Genomics Flex workflow

Single-cell suspensions were processed according to the 10x Genomics protocol for the fixation of cells and nuclei (CG000782 | Rev D). Briefly, suspensions of up to 10 million cells were spun down at 350 rcf for 5 min at 4°C and cells were resuspended in 1 mL of Fixation Buffer (4% Formaldehyde, 1X Conc. Fix and Perm Buffer B; 10x Genomics PN-2001301). Cells were fixed for 16–24h at 4°C, then spun down at 850 rcf for 5 min at room temperature and quenched with 1 mL of Quenching Buffer (1X Conc. Quench Buffer B; 10x Genomics PN-2001300). After quenching, 0.1 volumes of Enhancer (10x Genomics PN-2000482) and 10% glycerol were added for long-term storage.

Stored samples were processed by thawing at room temperature, centrifuging at 850 rcf for 5 min and resuspending in 1 mL 0.5X PBS – 0.02% BSA. Cell concentration and viability was assessed by AO-PI staining with a LUNA-FX7™ Automated Cell Counter (Logos Biosystems).

Where indicated, thawed and washed fixed cell suspensions were enriched by flow cytometry before downstream 10x Flex processing. Cancer cells were enriched by sorting dump-negative cells, defined as CD45⁻CD31⁻CD11b⁻TER119⁻. For lineage-tracing experiments, GFP⁺ cells were additionally enriched by flow cytometry when detectable by fluorescence and identified downstream by *GFP* transcript expression. Approximately 60,000–100,000 cells were sorted per sample.

Samples were pooled for hybridization. Up to 2 million cells were processed per hybridization following 10x Genomics user guide (CG000787 | Rev B). Hybridizations were set up in 40 µl of hybridization mix with 10 µl of Mouse WTA probes (10x Genomics PN-1000788) and supplemented with 2.5 µl of custom probes based on company recommendations (CG000621 | Rev D). Hybridizations were performed at 42°C for 16–24 hr. Hybridization samples were then diluted in Post-Hyb Wash Buffer and measured by AO-PI staining in a LUNA-FX7™ Automated Cell Counter (Logos Biosystems). For each experiment, we pooled an equal number of cells from each hybridization to have an equal distribution per sample. Cell pools were then washed 3 times in Post-Hyb Wash Buffer for 10 min at 42°C. Next, washed cells were resuspended in Post-Hyb Resuspension Buffer, filtered through a Miltenyi Biotec 30 µm filter and measured with the cell counter to determine the amount needed for the Chromium X run.

For the GEM encapsulation, we followed the 10x Genomics protocol and targeted 20,000 cells per sample up to 320,000 cells total for a pool of 16 samples. After loading the GEM-X FX Chip and running it on the Chromium X, GEMs were recovered and processed as indicated by 10x Genomics. After processing the GEMs, the product was pre-amplified and indexed to construct the sequencing library. All libraries were sequenced on an Illumina Novaseq X with standard dual indexing and demultiplexing. Raw .bcl files were processed using CellRanger (10x Genomics).

### Single-cell cancer cell-state analysis

#### Overview

To quantify how basal- and classical-derived cancer cells redistribute across transcriptional states during tumor evolution and after therapy, we derived compositional measures of classical, basal and mesenchymal cancer cell state and visualized distributional shifts in lineage-traced cells with ternary plots.

We assessed data from *KPF-basal* and *KPF-classical* autochthonous tumors, in which basal or classical cells were EGFP-labeled for 3 or 14 days of lineage tracing during unperturbed tumor evolution (**Fig. 2**). A second dataset was generated from orthotopic transplant of *KPF-basal* cell lines to establish synchronized tumor-bearing cohorts for treatment with vehicle, FOLFIRI, MRTX1133, or FOLFIRI plus MRTX1133 (**Fig. 7**). Across experiments, tumors were dissociated, and cancer cells were enriched by flow cytometry prior to fixation and sequencing. Briefly, each sample was preprocessed and normalized independently before jointly embedding samples from the same experiment and annotating cell type. *EGFP*-positive cells were then identified and scored for the three PDAC subtype signatures. Per-cell scores were converted into a softmax compositional score over the three states in lineage-marked cells at the harvest time point—thus making it possible to quantify transdifferentiation over time or state redistribution following therapy.

Analyses were performed using scanpy v1.12.1, anndata v0.12.16, and CellTypist v1.7.1. Doublets were detected using the Scrublet (*94*) implementation bundled in scanpy, and SciPy v1.17.1 was used for statistical fitting. For the transplant series, rapids-singlecell v0.15.2 with CuPy v14.1.1 was used for GPU acceleration.

#### Data preprocessing

For each sample, cells with fewer than 500 total counts and 200 detected genes were filtered, and doublets were scored and removed based on Scrublet with default thresholds. Data was library-size normalized to 10,000 counts per cell, followed by log transformation with a pseudocount of 1. The top 2000 highly variable genes (HVGs) were selected from the normalized data (sc.pp.highly_variable_genes, *flavor* = “seurat_v3”, *n_top_genes* = 2000) after excluding mitochondrial, ribosomal, and transgene-probe features, and the top 75 PCs on the HVGs (sc.pp.pca, *n_comps* = 75, *svd_solver* = “arpack”) were retained, explaining 60–70% of the variance in each sample. This was used to build a knn graph (*k* = 30) and UMAP visualization embedding (clustering *resolution* = 0.1) to identify the malignant compartment.

Each transgene was detected by three probes; thus, expression was quantified separately by summing the raw counts of each transgene’s three probe features per cell, library-size normalizing, and log-transforming with a pseudocount of 1.

#### Joint cell type annotation

Preprocessed samples from the same experiment were concatenated and re-embedded with batch-aware feature selection by adding *batch_key* = “sample_id” to the HVG computation, before computing PCs, generating a knn graph (*k* = 30), and visualizing with UMAP (*min_dist* = 0.3). The basal and classical GEMM series were processed using scanpy v1.12.1, with joint embeddings based on 100 PCs. Dimensionality reduction and clustering on the transplant series were performed with rapids-singlecell for GPU acceleration, using 75 PCs for the joint embeddings. PCs were chosen to capture 60-70% of the variance in both series.

CellTypist v1.7.1 (celltypist.annotate, *majority_voting* = true) was custom-trained on our Xenium panel and used for initial cell annotation, producing both per-cell and majority-vote labels. To identify the malignant compartment, the tumor and tumor-associated epithelial compartment was then subset into CellTypist labels cancer epithelial, cancer mesenchymal, acinar, ductal, ADM, mesothelial, tuft-like, and cancer epithelial neuronal-like. Mesothelial cells were included because they share markers (e.g., *Msln*) with malignant states and are commonly misassigned to them.

This subset was re-embedded (2000 HVGs, 100 PCs, *k* = 30, UMAP *min_dist* = 0.1) and clustered with Leiden (*resolution* = 1.0). Each cluster was then labeled by manual review of differentially expressed genes (Wilcoxon rank-sum test, sc.tl.rank_genes_groups) and canonical markers (gastric/classical: *Lgals4, Arg2, Tff2*; basal: *Slc4a11*; mesenchymal/EMT: *Vim, Lgals1, Twist1*; mesothelial: *Msln, Wt1, Adgrd1, Dcn*; hypoxic: *Vegfa, Ero1a*; intestine: *Pigr, Muc13, Alpi;* ADM/Acinar: *Cpa1, Rbpjl, Nr5a2*; ductal: *Prox1; Tuft: Pou2f3*; neuroendocrine: *Chga, Chgb, Neurod1*). Programs corresponding to gastric/classical, basal, and hypoxic states were collapsed into a single cancer epithelial label.

The final set of labels consisted of cancer epithelial, cancer mesenchymal, acinar, ADM, mesothelial, tuft-like, and intestine. Clusters that co-express distinct lineage programs were considered doublets and removed (complementing per-cell doublet removal with Scrublet during preprocessing). A few clusters within this compartment had non-epithelial identities, including mast cell, TAM (engulfed cancer transcripts), and erythroid, and were removed as contamination.

#### Reporter-positive cell assignment and cancer cell re-embedding

Cancer cells were defined as cells with cancer epithelial or cancer mesenchymal annotations. Due to the high levels of reporter transcript in the ambient RNA, we distinguished reporter-driven signal from background by fitting a gamma distribution (scipy.stats.gamma.fit, *floc = 0*) to the non-zero *EGFP* transgene expression of non-cancer cells; a cell was called *EGFP*⁺ if it was a cancer cell with transgene expression at or above an 80th percentile (scipy.stats.gamma.ppf, *q = 0.80*) *EGFP* positivity threshold, chosen to balance sensitivity and specificity against known negative controls.

Cancer cells (cancer epithelial and cancer mesenchymal) were subsetted, independently re-embedded (batch-aware HVG, PCA, *k* = 30, UMAP *min_dist* = 0.3) and Leiden clustered (*resolution* = 0.5). The resulting cancer-only annotated object for each experiment (*KPF-basal*, *KPF-classical*, and orthotopic transplant) served as the input to cell-state analysis.

#### Cancer cell state compositional scoring

For each experiment, cancer-only cells were each scored against basal, classical and mesenchymal signatures (see ***Panel and Gene Set Design*** for markers). Every gene was *z*-scored across cells; then, for each signature, the *z*-scores of its constituent genes were averaged per cell to yield a raw signature score. Because the signatures differ in gene number, and therefore in score variance, each of the three raw score columns were *z*-scored across cells to place them on a common scale. A row-wise softmax was applied across the three standardized scores (scipy.special.softmax) to obtain a non-negative per-cell compositional score over the three states that sums to one. Because the softmax maps each cell’s standardized scores onto the simplex, the values reflect how strongly each state is expressed relative to the population mean for that state.

Cell-state composition was visualized using per-cell compositional scores as barycentric coordinates (axes ordered classical–basal–mesenchymal) in ternary plots. Each plot is a per-cell scatter over the simplex, with points colored by local density (scipy.stats.gaussian_kde, *bw_method* = 0.30). Within each condition, cells were downsampled to the condition with lowest cell count using a fixed random generator so that point density was comparable across panels.

### Statistics and reproducibility

GraphPad Prism software v.9.5.1 was for statistical analyses or in-house scripts in Python, which are available from the corresponding author upon request. Statistical analyses were performed as indicated in the figure legends. For comparisons between untreated and treated samples at single time points, *p* values were determined using two-sided Welch’s *t* tests. When multiple pairwise comparisons were performed, *p* values were adjusted using Holm’s multiple comparisons correction. For comparisons across more than two groups, the statistical test used is specified in the corresponding figure legend. Investigators were not blinded to allocation during experiments or outcome assessment.

