## Supplementary Materials for "The basal cell state maintains pancreatic cancers by controlling an immunosuppressive circuit"

This file includes:

Figs. S1 to S12

Tables S1 to S4

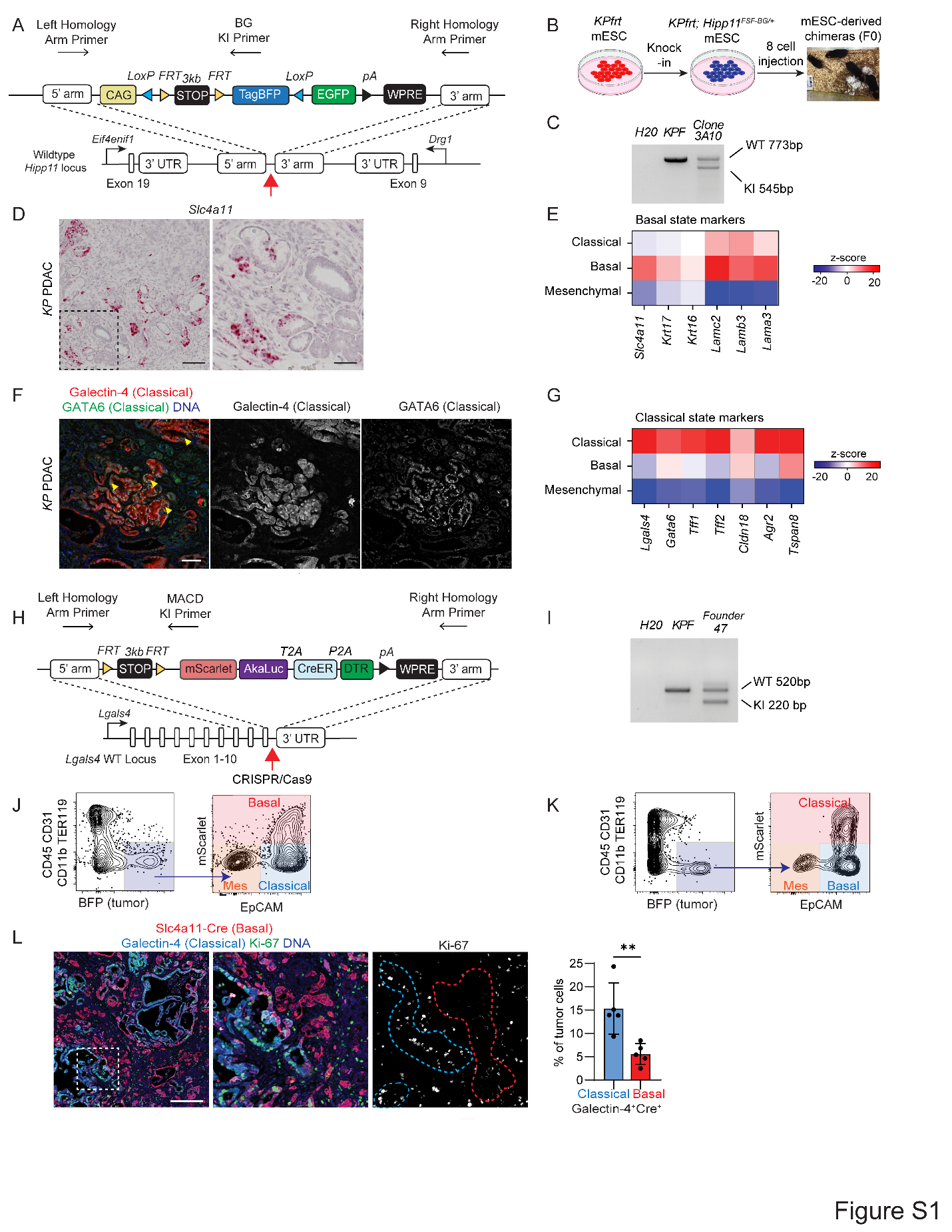

Fig. S1. Construction and validation of reporter and lineage tracing systems.

**(A)** *Hipp11^FSF-BG/+^* targeting strategy. The *FSF-BG* reporter construct was knocked into the *Hipp11* intergenic safe harbor locus (between *Eif4enif1* and *Drg1*) via CRISPR/Cas9 gene editing. *FlpO*-mediated recombination removes the *FSF* cassette to activate TagBFP, whereas Cre-mediated recombination removes the *LSL* cassette to permanently activate EGFP.

**(B)** Strategy to generate *Kras^FSF-G12D/+^; Trp53^frt/frt^; Hipp11^FSF-BG/+^* (*KPF*) reporter mouse. The *CAG-FSF-BG* construct was knocked into *KPfrt* mESCs, and correctly targeted clones were used to generate chimeric mice by 8-cell-stage embryo injection.

**(C)** Genotyping PCR of parental *KPfrt* mESCs and a correctly targeted ESC clone (Clone 3A10), from which chimeras were derived, showing background *KPfrt* mESC (WT) and knock-in (KI) alleles.

**(D)** *Slc4a11* mRNA expression in *KPfrt* PDAC. Scale bars, 100 µm (left) and 40 µm (right)

**(E)** Expression of key laminins and keratins, which mark the human basal subtype, across cancer cell states in unperturbed autochthonous *KP* PDAC tumors (*12*).

**(F)** Representative immunofluorescence in an autochthonous *KP* PDAC tumor showing colocalization of Galectin-4 with high levels of classical maker GATA6 (yellow arrowheads). GATA6 is also detected in stromal populations. Scale bar, 100 µm.

**(G)** Expression of key human-conserved classical markers across cancer cell states in unperturbed autochthonous *KP PDAC* tumors (*12*)

**(H)** *Lgals4^MACD/+^* targeting strategy. The FSF-MACD cassette was inserted in-frame after exon 10 of the *Lgals4* locus by CRISPR/Cas9 editing. The FSF-MACD reporter was introduced into *Kras^FSF-G12D/+^; Trp53^frt/frt^; Hipp11^BG/+^* embryos by zygote injection.

**(I)** Genotyping PCR of founder mice derived from injected embryos using primer pairs detecting WT and KI alleles.

**(J,K)** Flow cytometry analysis of *KPfrt; Slc4a11^MCD/+^; Hipp11^BG/+^* (H) and *KPfrt; Lgals4^MACD^*^/+^; *Hipp11^BG^*^/+^ (I) tumors. Tumor cells (BFP⁺; CD45⁻CD31⁻CD11b⁻CD11c⁻F4/80⁻ TER-119⁻) were sorted for cell state based on mScarlet and EpCAM expression. *n* = 5 mice for each genotype.

**(L)** Immunofluorescence (left) and quantification (right) of Ki-67 in *KPF-basal* PDAC tumors, comparing Cre⁺ basal and Lgals4⁺ classical cells. Ki-67 marks proliferating cells. *n* = 5 mice; **, *p* = 0.0046, Welch’s t-test.

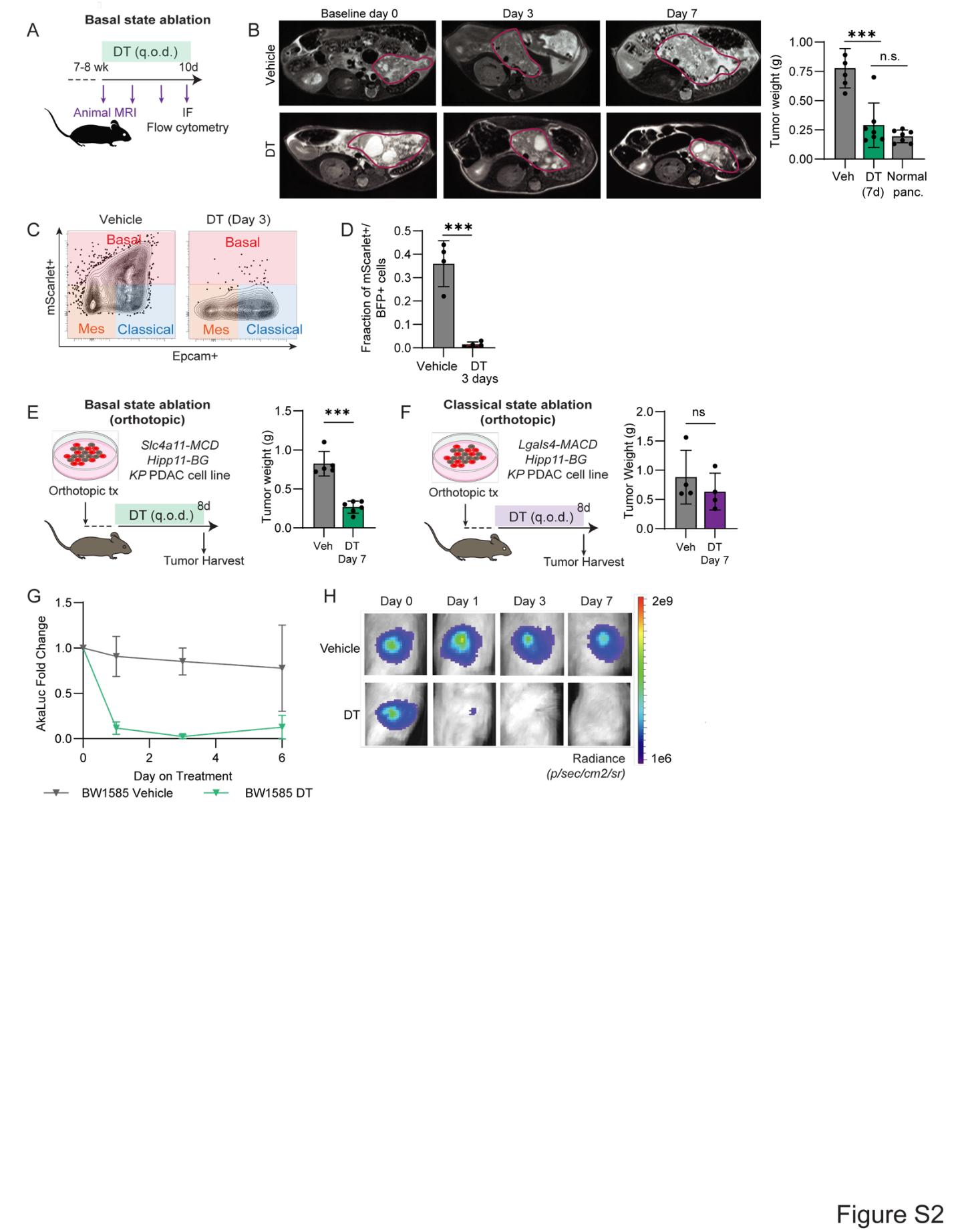

Fig. S2. Selective ablation of the basal cell state leads to rapid remodeling of the tumor ecosystem.

**(A)** Experimental strategy for DT-mediated basal-cell ablation in *KPF*-*basal* mice. Tumor volume was monitored by serial MRI; endpoint tumors were analyzed by flow cytometry and immunofluorescence.

**(B)** Representative MRI images showing tumor regression following DT-mediated basal-state ablation compared with normal saline control (left), with quantification and comparison to normal pancreas weights from age-matched non-tumor-bearing mice. *n* = 5–7 per group. *p* = 4.2*10^-4^, Welch’s t-test.

**(C,D)** Distribution of residual cancer cell states in DT and control-treated tumors by flow cytometry (C), with quantification (D). DT treatment leads to marked loss of mScarlet⁺ and presence of both EpCAM⁺ and EpCAM⁻ populations, consistent with specific loss of the basal state. *n* = 4 tumors per group. *p* = 5.4*10^-3^, Welch’s t-test.

**(E,F)** Orthotopic models used to compare basal-state (E) and classical-state (F) ablation. Only basal-state ablation results in a significant reduction in tumor weight. *n* = 4–7 tumors per group. ***, *p* = 5.1*10^-4^; ns, *p* = 0.41, Welch’s t-test.

**(G,H)** *In vivo* AkaLuc bioluminescence imaging of orthotopic *Lgals4*-MACD pancreatic tumors marking the *Lgals4*⁺ classical state, with quantification (G) and representative images (H), showing that DT-treated tumors lose the classical state population. *n* = 4 tumors per group.

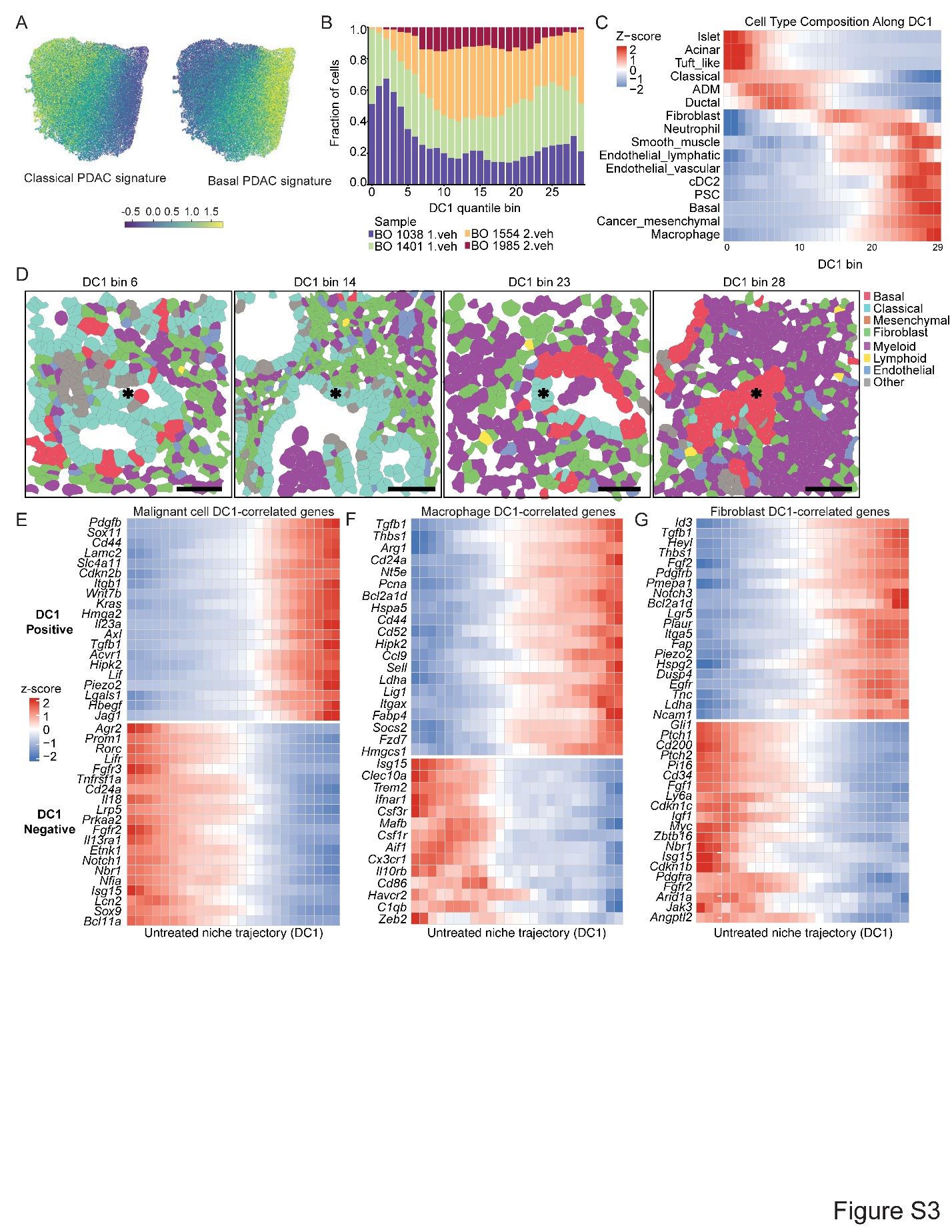

Fig. S3. Cellular and transcriptional programs associated with the classical–basal PDAC niche axis.

**(A)** UMAP visualization of Wormhole-embedded cancer cell niches from vehicle-treated *Slc4a11-*MCD tumors, colored by classical or basal transcriptional program.

**(B)** Distribution of vehicle-treated sample composition in cancer cell niches that are ordered and binned across the dominant diffusion component (DC1), showing broad sample mixing along the classical–basal niche axis.

**(C)** Cell-type composition across neighborhoods ordered along the classical–basal niche axis.

**(D)** Annotated Xenium data (Methods) showing representative cellular neighborhoods across DC1 bins spanning the classical (left) to basal (right) axes. Scale bar, 50 µm.

**(E–G)** Top positive-ranked and negative-ranked cancer cell genes (E), macrophage genes (F), or fibroblast genes (G) associated with DC1 (Spearman correlation *r* > 0.7 between binned expression and bin number).

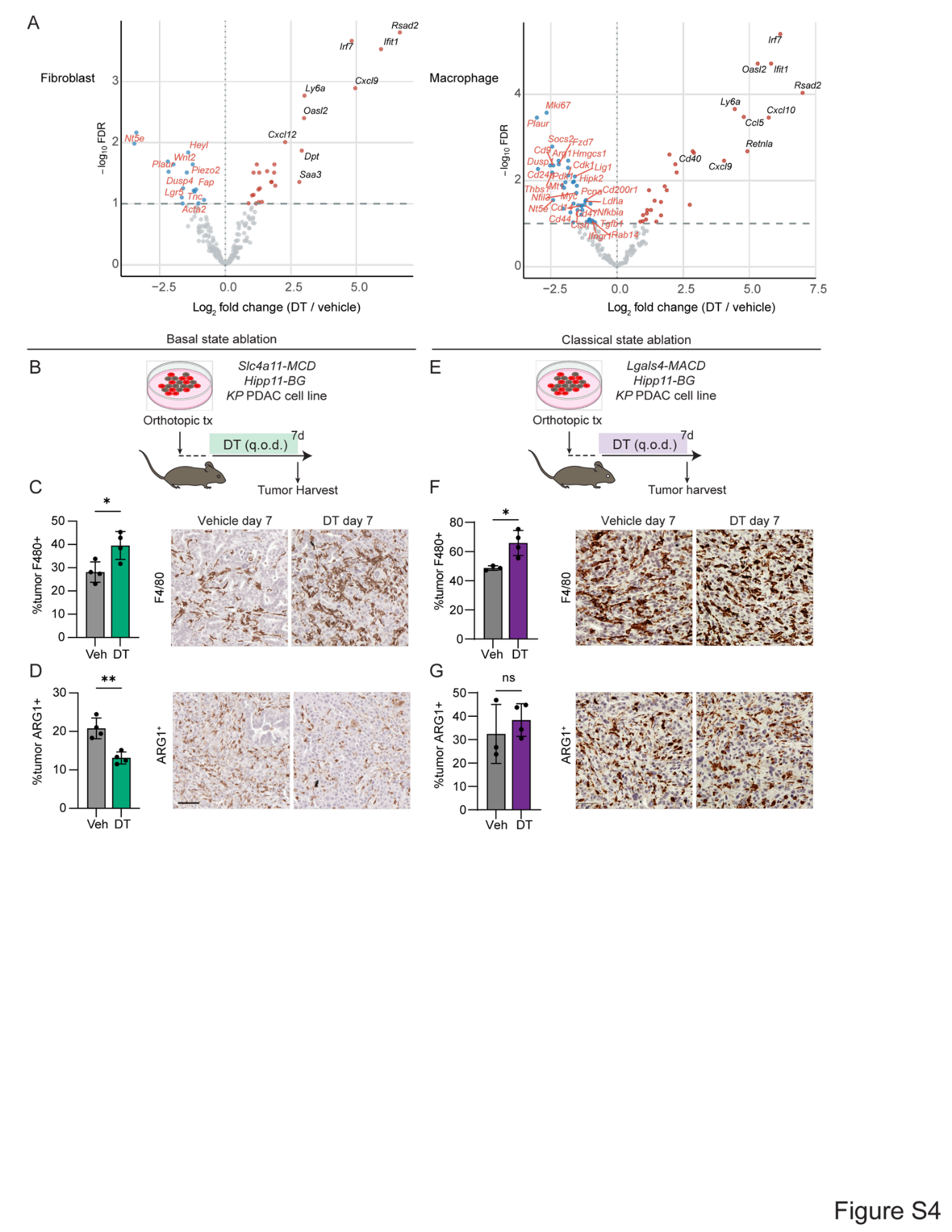

Fig. S4. A classical–basal cell axis organizes tumor microenvironment niches and is disrupted by basal cell ablation.

**(A)** Genes differentially expressed in all samples from day 2–6 DT-treated tumors (*n* = 5 control; n = 8 treated) compared to vehicle-treated controls for fibroblasts (left) and macrophages (right), showing significance (log_10_ FDR calculated for each cell type) of differential expression and fold change between treated and control. Genes highlighted in red are highly correlated with basal niches along DC1 (Spearman *r* > 0.7) and differentially expressed (adjusted P < 0.1, log_2_ fold-change > 0.5); genes highlighted in black are top differentially expressed genes after ablation.

**(B)** Experimental design for comparing ablation of *Slc4a11*-MCD^+^ cells in DT and control orthotopic tumors.

**(C)** Immunohistochemical staining and quantification of F4/80^+^ cells in vehicle-treated and DT-treated *Slc4a11-MCD* tumors. *n* = 4 mice per condition; error bars, s.d.; *, *p* = 0.0240, Welch’s *t*-test.

**(D)** Immunohistochemical staining and quantification of Arg1^+^ cells in vehicle-treated and DT-treated *Slc4a11-MCD* tumors. Scale bar, 50 μm. *n* = 4 mice per condition; error bars, s.d.; **, *p* = 0.0047, Welch’s *t*-test.

**(E)** Experimental design for comparing ablation of *Lgals4*-MACD^+^ cells in DT and control orthotopic tumors.

**(F)** Immunohistochemical staining and quantification of F4/80^+^ cells in vehicle-treated (*n* = 3) and DT-treated (*n* = 4) *Lgals4-MACD* tumors. Error bars , s.d.; *, *p* = 0.0253, Welch’s *t*-test.

**(G)** Immunohistochemical staining and quantification of Arg1^+^ cells in vehicle (*n* = 3) and DT (*n* = 4). Error bars, s.d.; ns, *p* = 0.5159, Welch’s *t*-test.

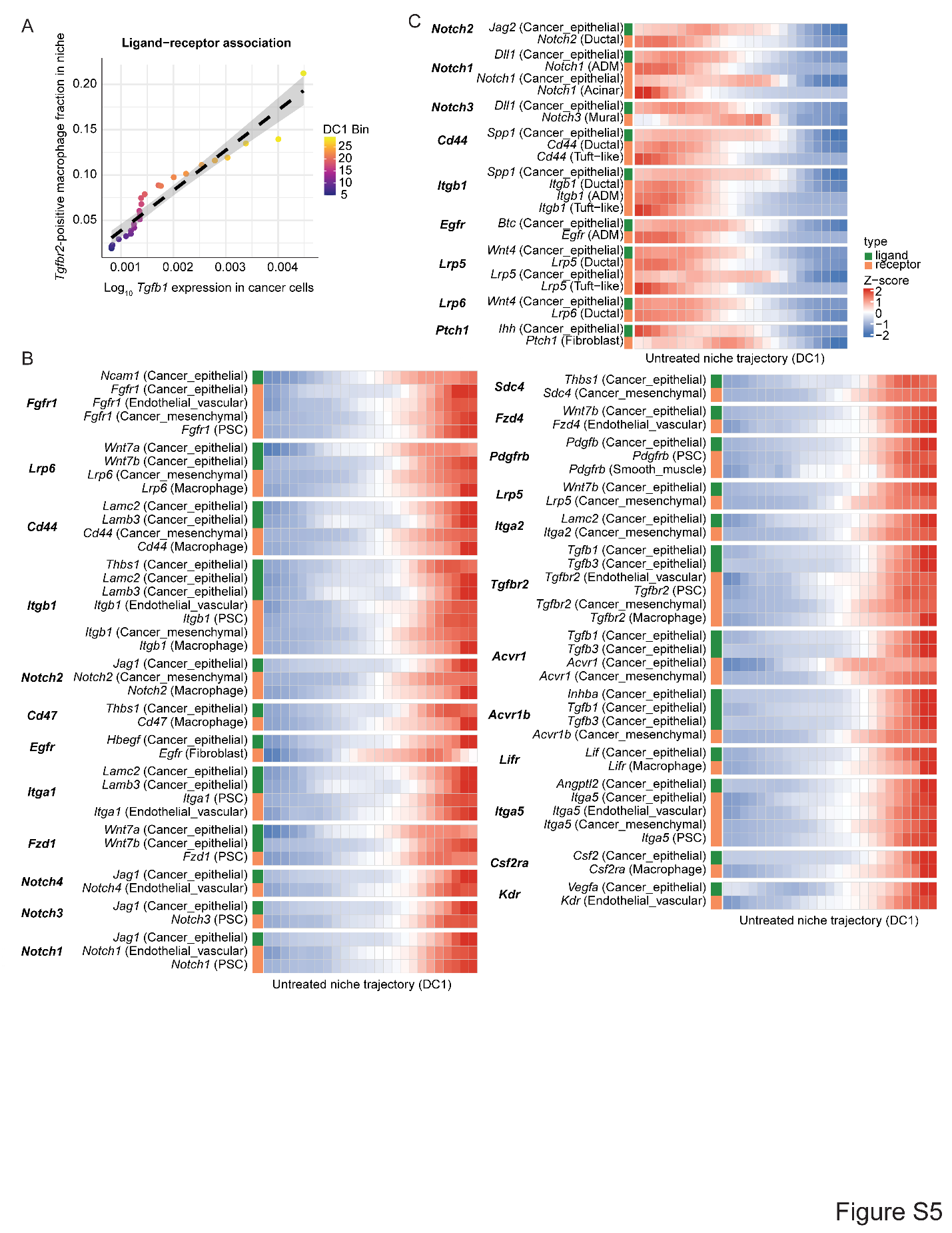

Fig. S5. Basal cancer cells nominate paracrine cytokine circuits.

**(A)** Example of ligand–receptor association, showing that expression of *Tgfb1* by cancer cells and fraction of cells (within the niche bin) positive for its cognate receptor *Tgfbr2* by macrophages in the same niche increase together. Both go up as niches range from classical-anchored to basal-anchored.

**(B,C)** Significant predicted sender–receiver relationships (Methods) showing source cell and ligand expression or cognate receptor-positive cell frequency along the untreated niche trajectory (DC1), for basal-associated (B) or classical associated (C) interactions.

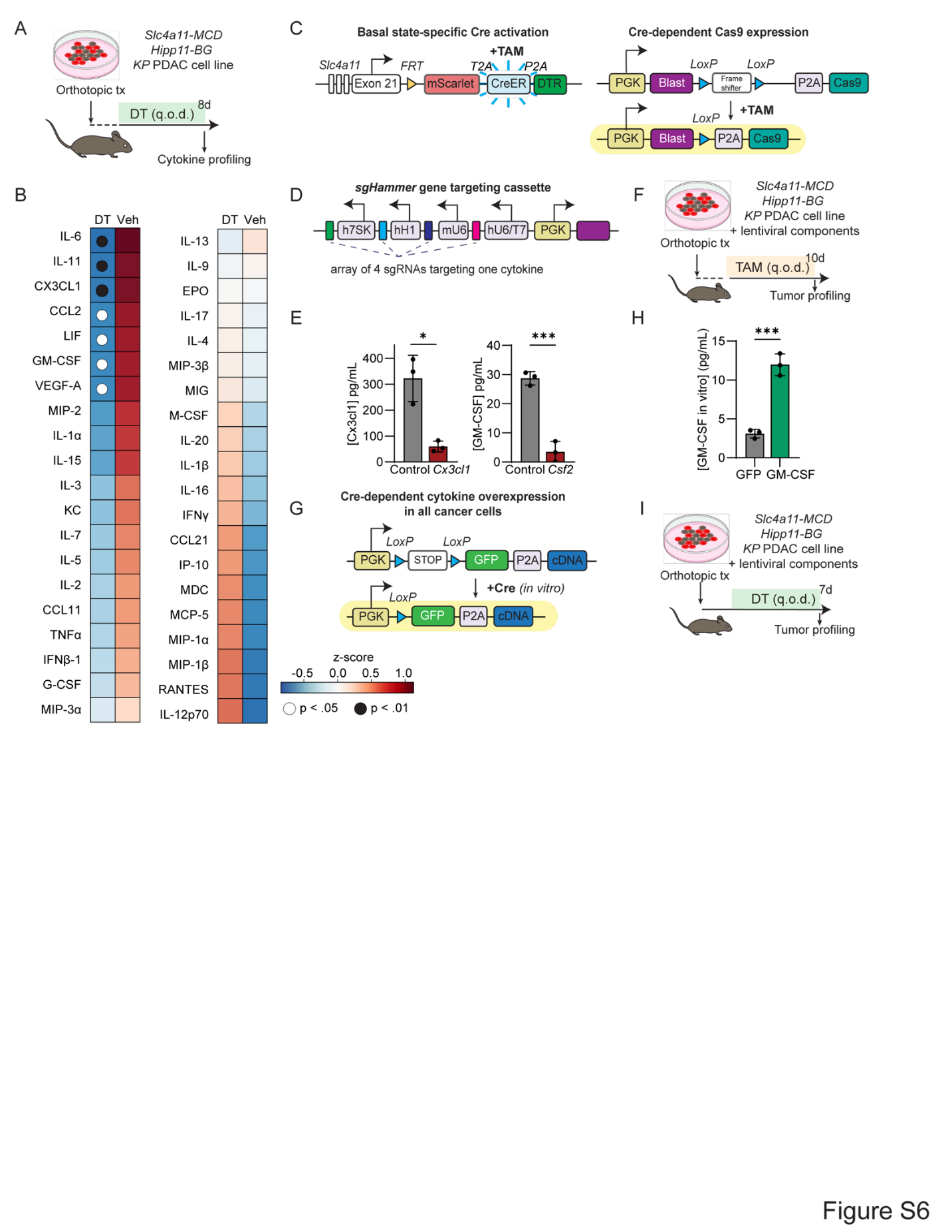

Fig. S6. GM-CSF produced by basal cancer cells enforces an immunosuppressive niche.

**(A)** Experimental design for cytokine profiling of vehicle- and DT-treated tumor tissues using the Eve Technologies 44-plex cytokine assay.

**(B)** Scaled mean cytokine concentrations in tumors treated with vehicle (Veh, *n* = 3) or DT (*n* = 4). Significant changes are indicated by circles with corresponding *p*-values.

**(C)** Inducible basal-state-specific Cas9 system. *In vivo* tamoxifen administration activates Cre recombinase specifically in *Slc4a11*-*MCD*⁺ (basal) cells, leading to Cas9 expression.

**(D)** sgHammer construct, consisting of an array of four sgRNAs targeting a single cytokine.

**(E)** Mean CX3CL1 and GM-CSF concentration by ELISA from *Slc4a11-MCD* supernatant of cells expressing *sgCtrl*, *sgCx3cl1*, or *sgCsf2* after *in vitro* Cas9 activation by CMV-Cre. Error bars, s.d.; *, *p* = 0.0308; ***, *p* = 0.0005, Welch’s t-test.

**(F)** Experimental design for *in vivo* basal-state-restricted cytokine knockout. Orthotopic tumors were treated with tamoxifen (q.o.d. for 10 days) to activate Cas9 in *Slc4a11-MCD*⁺ basal cells.

**(G)** Cytokine overexpression construct. *In vitro* Cre activation induces expression of a cDNA encoding the cytokine of interest.

**(H)** Mean GM-CSF concentration by ELISA from *Slc4a11-MCD* supernatant of cells expressing GFP or *Csf2* cDNA in all cells. Error bars, s.d.; **, *p* = 0.0037, Welch’s t-test.

**(I)** Experimental design for cytokine overexpression studies. Cell lines expressing the cytokine of interest or GFP control were used to generate orthotopic *Slc4a11-MCD* tumors. Tumors were then treated with DT or vehicle control and profiled at endpoint.

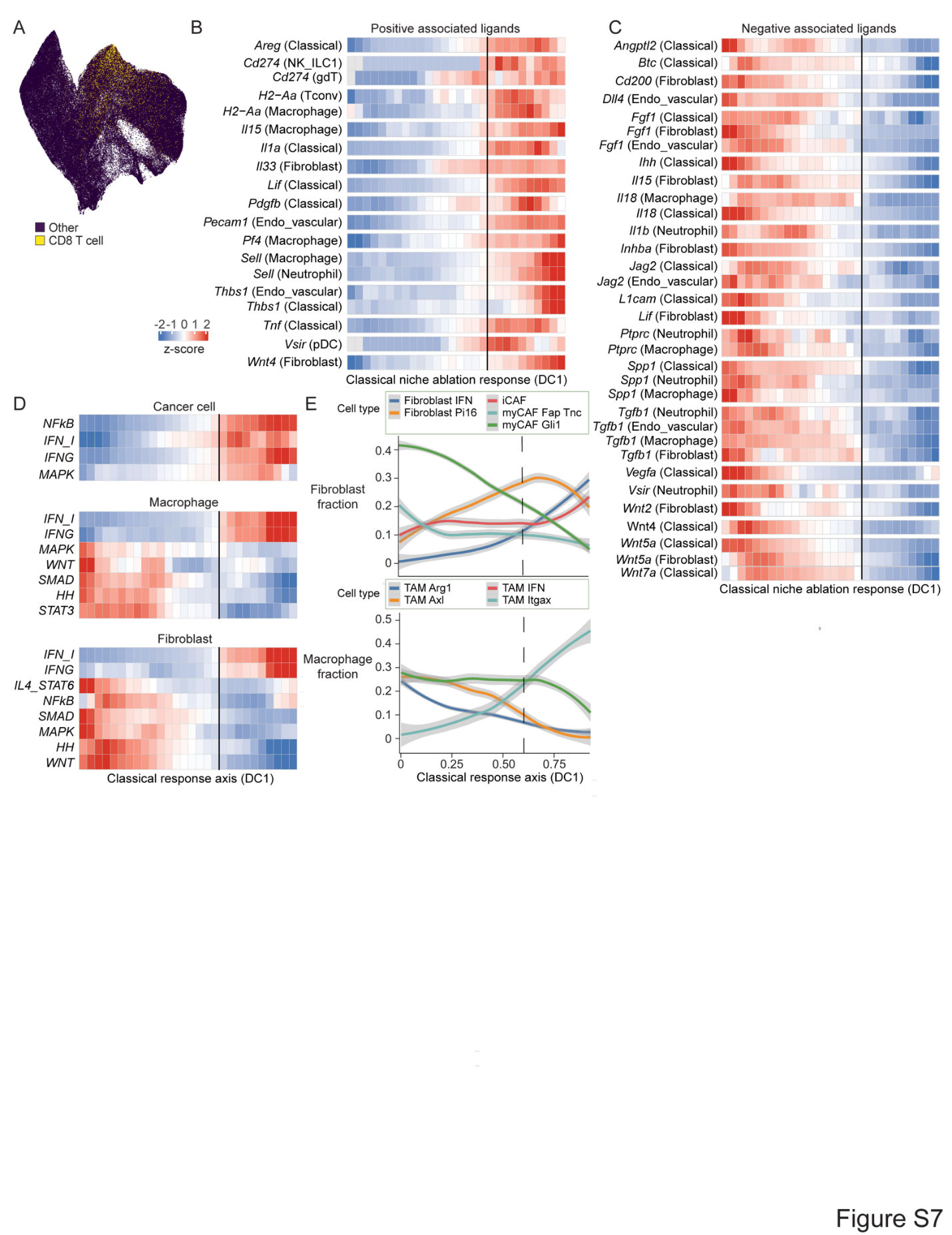

Fig. S7. Classical response–associated cell-state and signaling programs in basal ablation time series.

**(A)** UMAP embedding of all niches as in Fig. 6A, labeled by presence or absence of CD8 T cells within the niche.

**(B,C)** Scaled expression (*z*-scored by row) along classical niche response axis, showing ligands with expression that is positively (B) or negatively (C) associated with classical response axis bin number (Spearman *r* > 0.7), and their source cell. Line corresponds to bin at which the untreated fraction drops below 10% of niche bin fraction.

**(D)** Scaled expression (z-scored by row) along classical niche response axis, showing signaling response gene programs that are correlated with classical niche response axis bin in cancer cells, macrophages and fibroblasts. Line corresponds to bin at which the untreated fraction drops below 10% of niche bin fraction.

**(E)** Fraction of fibroblast (top) or macrophage (bottom) granular cell states out of all fibroblasts or macrophages, respectively, along the classical response niche axis. Dashed line corresponds to bin at which the untreated fraction drops below 10% of niche bin fraction.

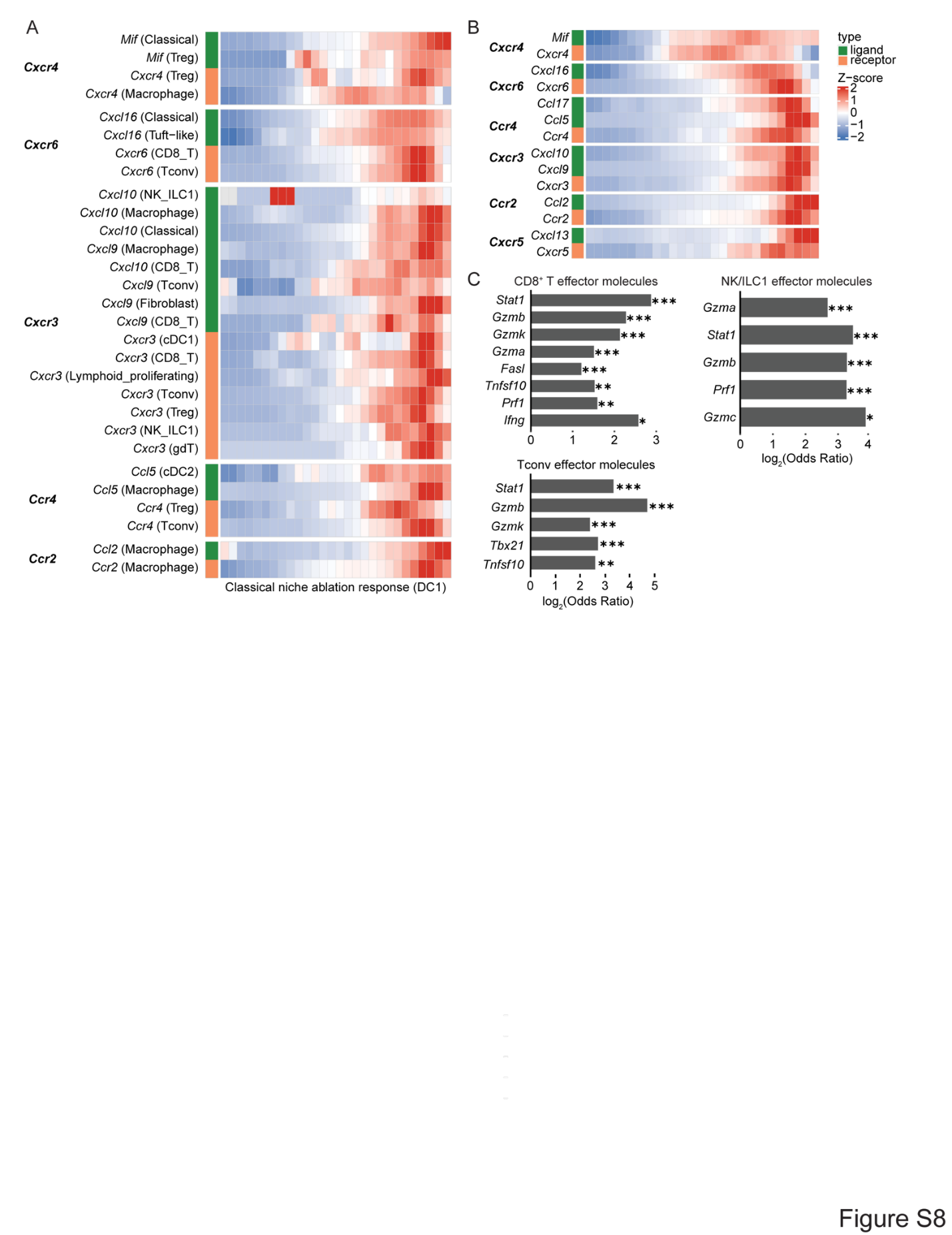

Fig. S8. Classical response–associated chemokine and effector molecular relationships in basal ablation time series.

**(A,B)** Scaled expression (*z*-scored by row) along classical niche response axis, showing significant cell-type-level (A) or niche-level (B) chemokine ligand–receptor relationships (see Methods for significance determination).

**(C)** Log_2_ odds ratio of effector-molecule-positive cell frequency in early (3-13) compared to late (17-27) bins in the classical response axis. *, *p* < 0.05; **, *p* < 0.01; ***, *p* < 0.001, Fisher’s exact test.

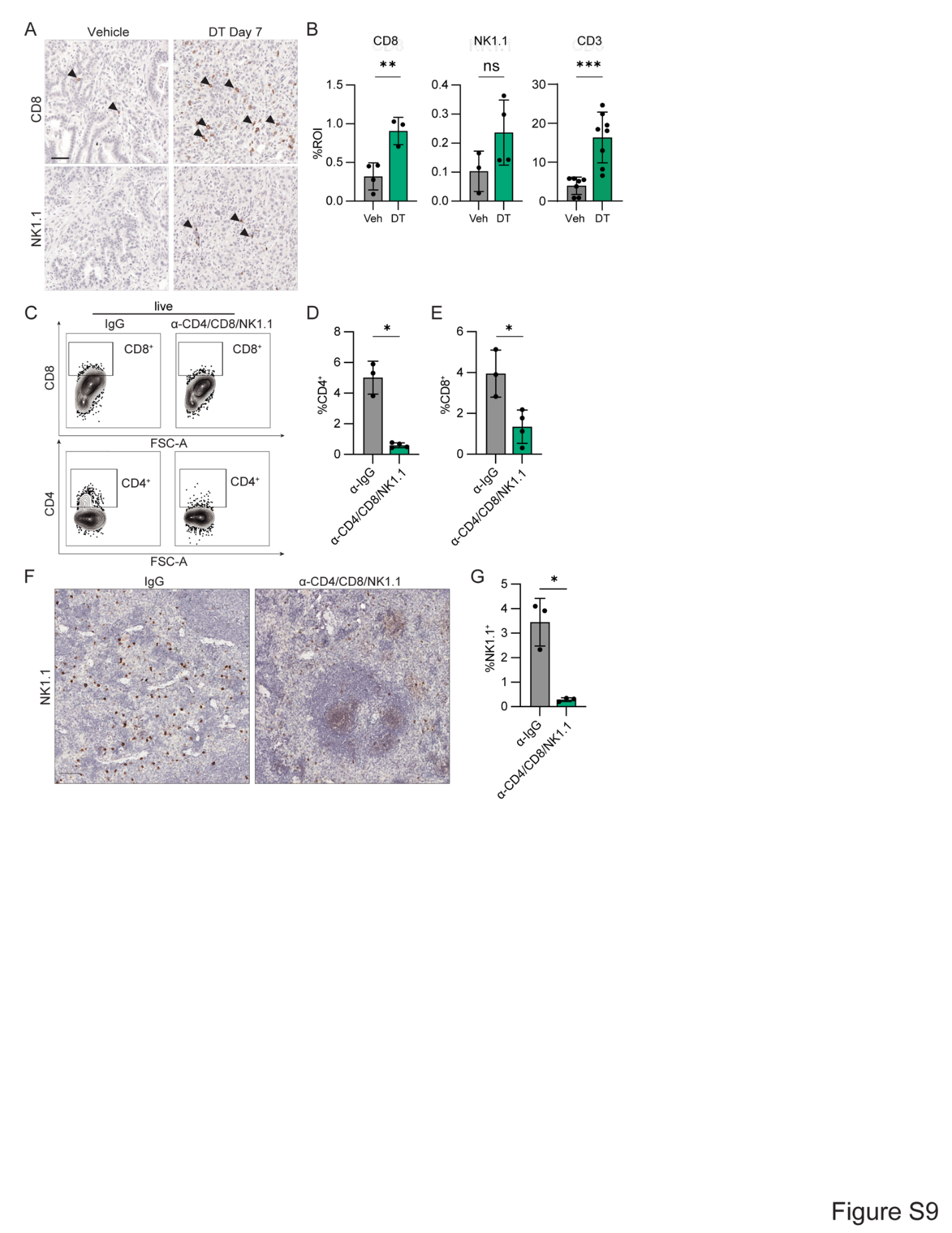

Fig. S9. Basal state ablation induces sequential immune recruitment and T/NK cell-mediated tumor killing.

**(A)** Representative IHC images of CD8 and NK1.1 staining in F1 orthotopic tumors 7 days after vehicle or DT treatment. Basal-state ablation is associated with increased infiltration of CD8⁺ T cells and NK cells.

**(B)** Mean cytotoxic lymphocyte marker-positive cell percentage per ROI following basal-state ablation compared with vehicle control. Error bars, s.d.; **, *p* < 0.01; ***, *p* < 0.001; ns, *p* > 0.05, Welch’s *t*-test.

**(C)** Representative flow cytometry analysis of orthotopic *Slc4a11*-MCD tumors treated with IgG (*n* = 3) or depletion antibodies (*n* = 4).

**(D, E)** Mean percentage of all live cells expressing CD4 (D) or CD8 (E) in IgG (*n* = 3) or CD4/CD8/NK1.1 (*n* = 4) treated tumors. Error bars, s.d.; *, *p =* 0.0174 (D) and 0.0358 (E), Welch’s t-test.

**(F, G)** Immunohistochemical staining and quantification of NK1.1^+^ cells in spleens of mice treated with anti-IgG (*n* = 3) or anti-CD4/CD8/NK1.1 (*n* = 3). Error bars, s.d.; *, *p* = 0.0296, Welch’s *t*-test.

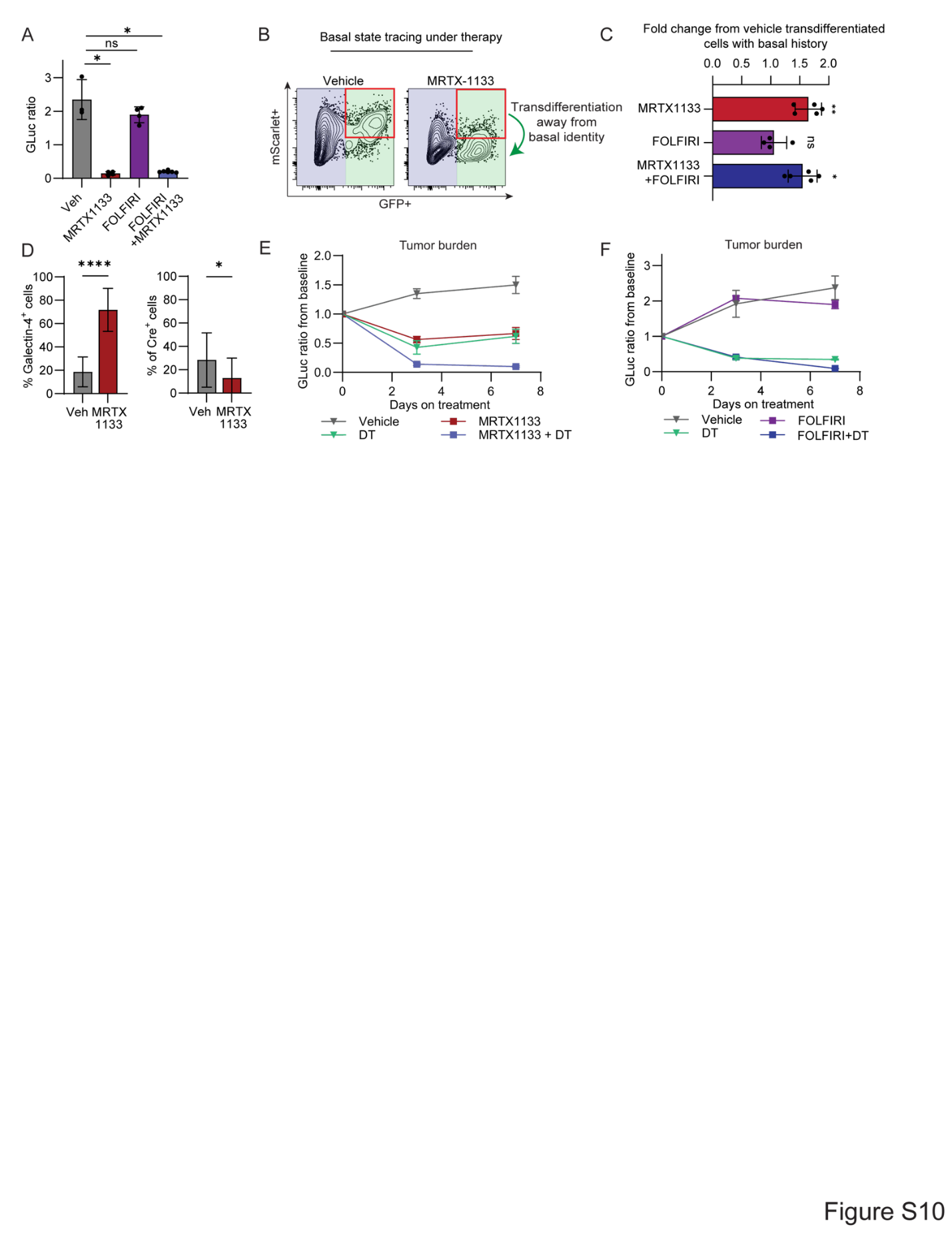

Fig. S10. Therapeutic targeting of the basal state.

**(A)** Mean tumor burden at day 7 of chemotherapy, KRAS inhibitor treatment, or both, measured by GLuc in orthotopic *KPF-basal* tumors. *n* = 3–6 mice per group; error bars, s.d.; Welch’s t-test with correction for multiple comparisons. *, *p* = 0.0230 (MRTX1133 vs. Vehicle), 0.0246 (FOLFIRI + MRTX1133 vs Vehicle); ns, *p* = 0.318, Welch’s t-test.

**(B)** Representative flow cytometry plots of live tumor cells gated as CD45⁻CD31⁻CD11b⁻TER119⁻DAPI⁻ in basal lineage-traced tumors after treatment. A GFP⁺mScarlet⁻ population emerges under MRTX1133 treatment, indicating exit from the basal state.

**(C)** Quantification of basal-state exit in lineage-traced tumors, measured as mScarlet⁻GFP⁺ cells/total GFP⁺ cells and shown as fold change relative to vehicle. *n* = 4–5 tumors per group; error bars, s.d. ns, adjusted *p* = 0.645; *, adjusted *p* = 0.0111; **, adjusted *p* = 0.00483, unpaired Welch’s t-test with Holm correction for multiple comparisons.

**(D)** Mean percentage of GFP^+^ cells overlapping with Slc4a11-Cre or Galectin-4 staining in basal lineage-traced autochthonous tumors after 7 days of vehicle or MRTX1133 treatment. Each point represents an individual tumor lesion or region. *n* = 13–17 lesions per group; error bars, s.d.; *, *p* = 0.0452; ****, *p* = 1.84 × 10⁻⁸, unpaired Welch’s t-test.

**(E,F)** Mean tumor burden measured over time by GLuc in response to basal ablation with KRAS inhibitor therapy (E) or chemotherapy (F), or those treatments alone. *n* = 7–10 mice (E) or 3–7 mice (F) per group; error bars, s.e.m.

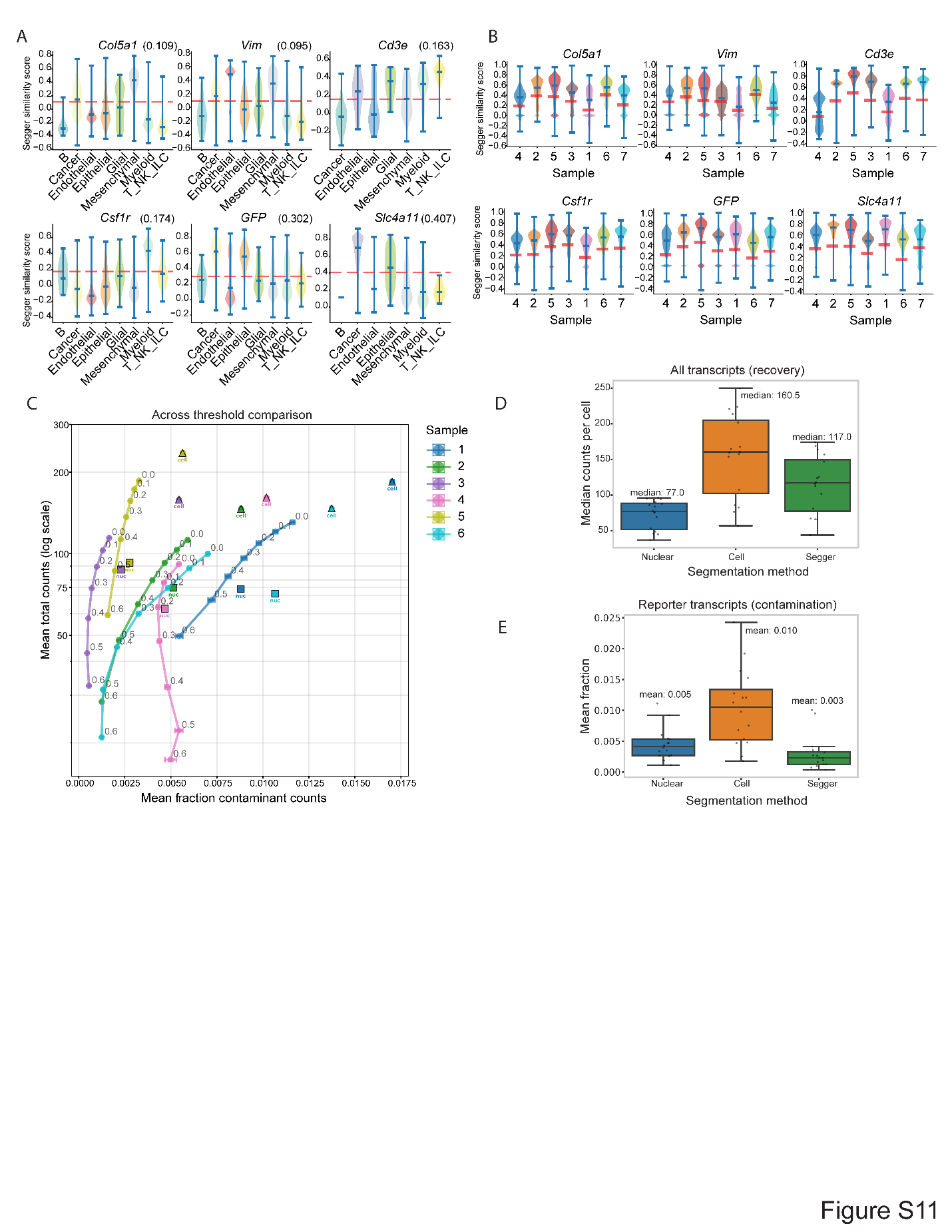

Fig. S11. Benchmarking of cell segmentation and transcript assignment.

**(A)** Segger similarity score distributions by cell lineage (*n* denotes number of cells of each type) for select marker genes within a representative sample. Dashed red line indicates threshold for assignment to a given cell. A high similarity score indicates that a transcript is strongly associated with a cell boundary based on local context, cell identity, and gene co-expression across the tissue.

**(B)** Segger similarity score distributions across samples for select marker genes show automated thresholds adapting to sample-specific similarity score distributions in a consistent way. Red lines indicate automated assignment threshold values for each sample.

**(C)** Segger transcript assignment thresholds affect purity and transcript recovery. Thresholds were increased by 10–60% of the difference between the automated thresholding and maximum similarity score values across all genes in six representative samples. Squares and triangles indicate metrics for 10x Genomics software default nuclear and cell segmentations, respectively. Fraction of contaminant counts was computed by the number of GFP, tagBFP2, and CreER counts in non-epithelial cells.

**(D,E)** Median per-cell counts of all transcripts (D), or mean fraction of GFP, tagBFP2, and CreER transcripts in non-epithelial, non-cancer cells (E) across all samples (*n* = 16).

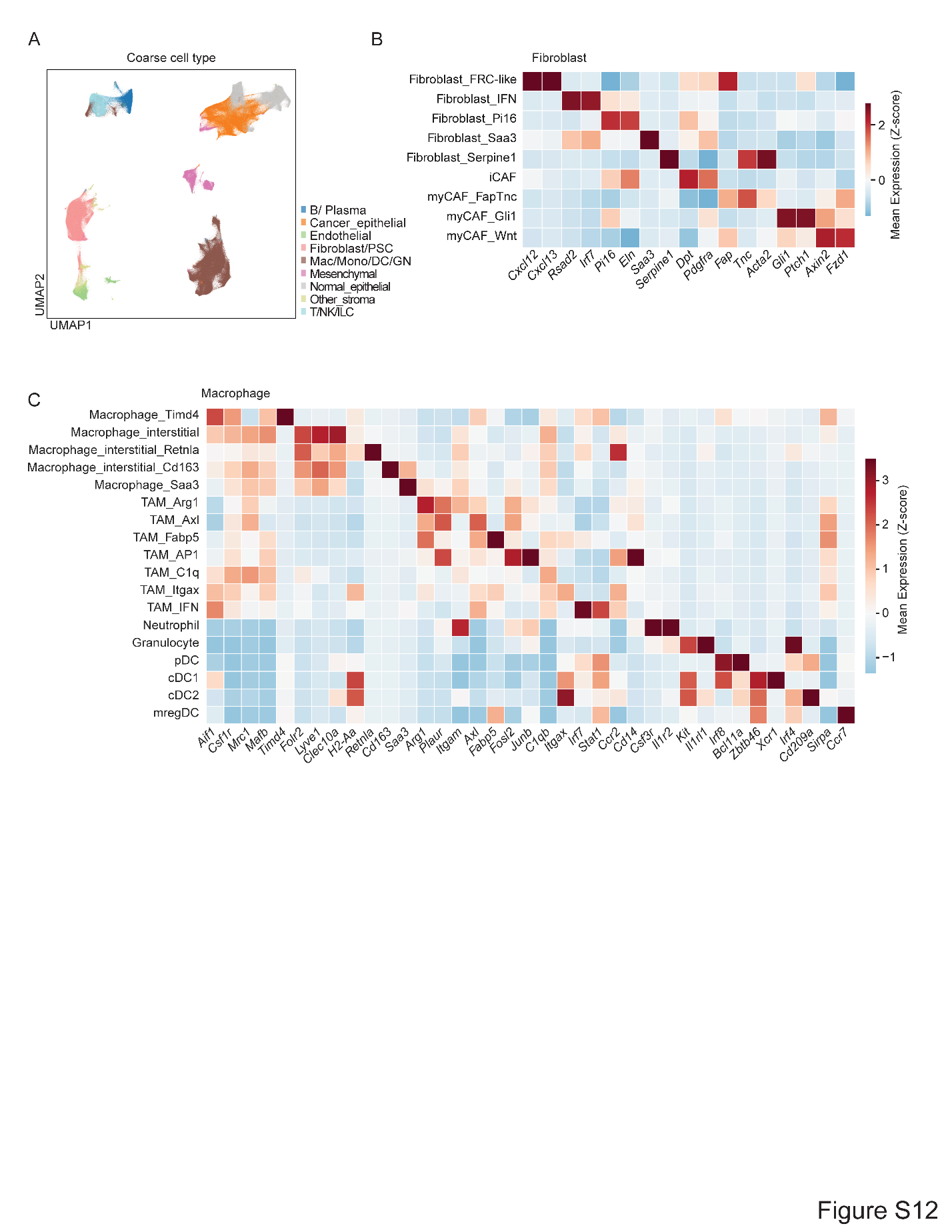

Fig. S12. Xenium cell typing.

**(A)** Single-cell transcriptional embedding of Xenium data, annotated by broad cell lineage (*n* = 3,091,255 cells).

**(B, C)** Marker gene expression in fibroblast (B) and myeloid (C) granular cell annotations. Mean log1p-normalized expression values within each cell annotation were *z*-scored across annotations for visualization.

Table S1. Xenium probe panel, cancer cell state gene sets, sample-level metadata and quality-control metrics.

Table S2. Differentially expressed genes supporting Xenium cell-type annotation, macrophage and fibroblast expression changes over the basal ablation time course, and cytotoxic lymphocyte effector frequency over the classical-response trajectory.

Table S3. Genes and putative ligand–receptor interactions correlated with diffusion component 1 in untreated tumor niches and along the classical-response trajectory.

Table S4. Experimental reagents used in this study, including primers, antibodies, and vectors.
